# A eukaryote-wide perspective on the diversity and evolution of the ARF GTPase protein family

**DOI:** 10.1101/2020.10.31.363457

**Authors:** Romana Vargová, Jeremy G. Wideman, Romain Derelle, Vladimír Klimeš, Richard A. Kahn, Joel B. Dacks, Marek Eliáš

## Abstract

The evolution of eukaryotic cellular complexity is interwoven with the extensive diversification of many protein families. One key family is the ARF GTPases that act in eukaryote-specific processes, including membrane traffic, tubulin assembly, actin dynamics, and cilia-related functions. Unfortunately, our understanding of the evolution of this family is limited. Sampling an extensive set of available genome and transcriptome sequences, we have assembled a dataset of over 2,000 manually curated ARF family genes from 114 eukaryotic species, including many deeply diverged protist lineages, and carried out comprehensive molecular phylogenetic analyses. These reconstructed as many as 16 ARF family members present in the last eukaryotic common ancestor (LECA), nearly doubling the previously inferred ancient system complexity. Evidence for the wide occurrence and ancestral origin of Arf6, Arl13 and Arl16 is presented for the first time. Moreover, Arl17, Arl18 and SarB, newly described here, are absent from well-studied model organisms and as a result their function(s) remain unknown. Analyses of our dataset revealed a previously unsuspected diversity of membrane association modes and domain architectures within the ARF family. We detail the step-wise expansion of the ARF family in the metazoan lineage, including discovery of several new animal-specific family members. Delving back to its earliest evolution in eukaryotes, the resolved relationship observed between the ARF family paralogs sets boundaries for scenarios of vesicle coat origins during eukaryogenesis. Altogether, our work fundamentally broadens the understanding of the diversity and evolution of a protein family underpinning the structural and functional complexity of the eukaryote cells.

**Significance:** ARF Family GTPases are crucial regulations of a diversity of cellular compartments and processes and as such the extent of this system in eukaryotes reflects both cellular complexity in modern eukaryotes and its evolution. Strikingly, a comprehensive comparative genomic analysis of the protein family is lacking, leaving its recent and ancient evolution poorly resolved. We performed a comprehensive molecular evolutionary analysis, reconstructing a highly complex ARF family complement in the Last Eukaryotic Common Ancestor, including a number of paralogs never before identified as such, and we find resolved relationships between the paralogs. This work has implications for cellular evolution from eukaryogenesis to cellular complexity in metazoans.

## Introduction

Understanding how the eukaryotic cell evolved in all its complexity is one of the greatest open questions in evolutionary biology. Eukaryogenesis involved both the origin of new genes and the diversification of key building blocks (Dacks et al. 2016; Eme et al. 2017). Among the different building blocks, particular groups of proteins radiated early in the evolution of eukaryotes and are represented by a large number of pan-eukaryotic orthologs, presumably with conserved functions. One of the largest groups of proteins, acting in an incredibly diverse array of cellular pathways, is the Ras superfamily of GTPases. This superfamily is frequently equated with familiar and extensively studied eukaryotic “small GTPases”. However, the more appropriate, i.e. evolutionary, definition conceives it as a major monophyletic subgroup of the vast TRAFAC class of GTPases that also includes prokaryotic representatives, larger proteins combining a Ras-related GTPase domain with other functional domains, and – surprisingly to many in the field – the alpha subunits of heterotrimeric G-proteins (Leipe et al. 2002). Because of its central role in so many fundamental cellular functions, understanding the origin and evolution of this complex superfamily of proteins is necessary for uncovering the processes by which eukaryotes evolved and diversified.

The internal classification of the Ras superfamily is unsettled. In many overviews, especially those concentrating on the eukaryotic small GTPases, the content of the superfamily is pigeonholed into five major families (Ras, Rho, Rab, Ran, Arf/Sar; Colicelli 2004; Rojas et al. 2012), but this scheme ignores the prokaryotic superfamily members (Wuichet and Søgaard-Andersen 2014), multi-domain proteins (such as the ROCO family; Bosgraaf and Van Haastert 2003), and various other lineages clearly distinct from or not easily classified into the well known families, such as the Gtr/Rag family (Klinger, Spang et al. 2016) or RJL proteins (Elias and Archibald 2009). Understanding the diversity and the evolutionary origin of the Ras superfamily in eukaryotes is a challenging task, given the presence of tens to hundreds of Ras superfamily genes in each extant eukaryote genome (Rojas et al. 2012). Disregarding potential (presently unknown) cases of horizontal gene transfer from prokaryotic sources into particular eukaryote lineages, the wealth of Ras superfamily genes in eukaryotes ultimately derives from a set of genes present in the Last Eukaryote Common Ancestor (LECA). Several evolutionary analyses have attempted to reconstruct LECA’s complement of particular Ras superfamily subgroups and detail the downstream innovation within eukaryotes. Prominent examples include analyses of the Rab (Diekmann et al. 2011; Elias et al. 2012; Klöpper et al. 2012) and Ras families (van Dam et al. 2011), and some isolated lineages like RJL (Elias and Archibald 2009), Miro (Vlahou et al. 2011), or RABL2 (Eliáš et al. 2016). These investigations demonstrated that a large number of functionally investigated paralogs were present in the LECA, emphasizing the role of loss or streamlining of genomic complement in many eukaryotic lineages. They also identified ancient LECA paralogs of unknown function that have been lost in lineages leading to conventional model systems but which are present in diverse eukaryotic lineages of ecological and medical importance. Paralogs with such an evolutionary distribution were recently coined jotnarlogs (More et al. 2020). Finally, these studies also inevitably shed light on the diversification of GTPases in the post-LECA expansion phase. For example, divergent paralogs of unclear evolutionary relationships are found in various taxa (e.g., Pereira-Leal 2008), most likely resulting from rapid sequence evolution of lineage-specific paralogs linked to their neofunctionalization. Additionally, the inherently small nature of the GTPases makes them particularly susceptible to molecular tinkering, such as accretion of additional domains or gain/loss of motifs mediating specific post-translational modifications (e.g., Záhonová et al. 2018).

Not yet addressed in a comparable evolutionary framework is the ARF protein family. This large protein family is comprised of the “true” ADP Ribosylation Factors (i.e., Arfs), as well as Arf-like proteins (Arls), Arf-related protein 1 (Arfrp1), and Sar1. Clearly related are the beta subunits of the signal recognition particle receptor (SRβ; Schwartz and Blobel 2003). Sequence analyses have also revealed that an Arf-like ancestor, modified by insertion of a novel α-helical region into its GTPase domain and high sequence divergence, gave rise to the alpha subunits of heterotrimeric G-proteins (abbreviated Gα; Neuwald 2007; Anantharaman et al. 2011). The distinction between Arf and Arf-like (Arl) proteins was originally made based upon activity in the cholera toxin-catalyzed ADP-ribosylation of the stimulator of adenylyl cyclase, Gα_s_, as all tested Arfs retain this functionality while the Arls did not (Tamkun et al. 1991; Clark et al. 1993). However, this activity has proven of very limited utility in studies of cellular functions for ARF family members as greater appreciation of both the size of the family in model organisms as well as the diversity of functions became clear. Thus, little if any weight should be given to whether a gene is named as an Arf, an Arl, an Arfrp1, or a Sar. The ARF family is functionally heterogeneous and comprises proteins involved in membrane vesicle formation (Arfs, Sar1), other aspects of vesicle traffic and maintenance of membranous organelle morphology (e.g., Arl1, Arl5, or Arfrp1), microtubule dynamics and mitochondrial fusion (Arl2), and cilium biogenesis and function (Arl3, Arl6, Arl13) (Gillingham and Munro 2007; Donaldson and Jackson 2011; Francis et al. 2016). Members of this family are critical to these diverse cellular activities and dysfunction results in numerous human diseases. Family members are generally considered to be single-domain small GTPases. Post-translational modifications (N-terminal myristoylation or acetylation) are also often critical to the protein’s localization and function.

An early phylogenetic study on the ARF family, limited by a lack of taxonomic breadth in available genomic sequences, provided an early estimate of the ancient complexity of the family in LECA and identified putative lineage-specific expansions in metazoans (Li et al. 2004). The analyses showed that LECA contained at least eight ancient groups of orthologs inferred from representatives being present in metazoans and at least one non-opisthokont (protist or plant) eukaryote. This analysis also demonstrated that some of the metazoan family members lacked close relatives in other eukaryotes, suggesting that lineage-specific expansions related to metazoan multicellularity occurred. Perhaps most familiar is expansion yielding the well-known and founding members of the family, Arfs 1-5. These have been shown as deriving from a single ancestral gene (here referred to as Arf1 for simplicity) which duplicated prior to choanoflagellates, yielding Arfs 1-3 (sometimes named Class I Arfs but for convention referred to here as Arf1) and Arfs 4-5 (sometimes named Class II Arfs but for convention referred to here as Arf4), with each of those diversifying into five Arf paralogs around the whole genome duplications in the vertebrate lineage (Manolea et al. 2010). However, since these early studies, several family members from the target species (including humans) have been identified (Kahn et al. 2006) and methods of phylogenetic analyses of protein sequences have advanced, including the development of the ScrollSaw approach facilitating analyses of complex paralog-rich families (Elias et al. 2012). Thus, the time is ripe for obtaining a much better picture of the evolution of ARF family than in the previous studies.

To this end, we assembled, extensively curated and phylogenetically analysed a dataset of ARF family sequences from a taxonomically broad selection of eukaryotic species. This enabled us to revise the set of ancestral eukaryotic ARF family paralogs, which has now expanded to between 14 and 16 genes. Two paralogs, described here for the first time, are not represented in well studied models and point to hitherto unstudied molecular functions mediated by the ARF family. We observed an unexpected diversity of domain architectures challenging the dogma that ARF family proteins are only small and single-domain proteins. Our analyses also unveiled a range of predicted post-translational modifications (PTMs), including but not limited to well-established N-terminal myristoylation, and other molecular adaptations that facilitate membrane association as a central feature of ARF family biology. Finally, we identified well supported relationships between the paralogs, which have implications for the inferred function of the primordial family members during eukaryogenesis.

## Results and Discussion

### A comprehensive dataset and phylogeny of the ARF family

We first gathered all ARF family sequences (including SRβ but excluding the highly divergent Gα proteins) from a broad diversity of eukaryotes, exploiting both publicly available and privately curated genomes and transcriptomes. We did not rely solely on predicted protein sequence sets but also checked the genome and transcriptome assemblies to ensure maximal accuracy when it comes to statements about the absence of particular genes in different taxa. All sequences were carefully validated, as described under Materials and Methods, and when needed, edited (by modifications of the originally predicted gene models or by changes in the assembled nucleotide sequences based on inspection of raw sequencing data) to ensure maximal quality and completeness of the data. Our final dataset, provided as supplementary dataset 1 (Supplementary Material online), included >2,000 manually curated sequences from 114 species (supplementary table 1, Supplementary Material online). The number of ARF family genes in individual species ranged from 5 in the yeast *Schizosaccharomyces pombe* to 70 in the rotifer *Adineta vaga* (this high number apparently reflecting the tetraploid origin of its genome; Flot et al. 2013).

The genes were initially annotated based on their similarity to previously characterized or named ARF family genes in model organisms scored by BLAST. While this procedure enabled us to recognize candidate groups of orthologs and to assign most of the genes into these groups, the assignment of many sequences was uncertain or unclear and a more rigorous method for establishing orthologous relationships – phylogenetic analysis of a multiple sequence alignment – was required to corroborate the proposed groups of orthologs and to possibly identify additional ones not readily apparent from sequence-similarity comparisons. Such an analysis of the whole dataset was impractical, if not impossible, for its size and the existence of divergent sequences that tend to disrupt the results of phylogenetic inference. We therefore utilized of the ScrollSaw protocol previously developed to deal with a similarly complex family of Rab GTPases (Elias et al. 2012) and applied by others to resolve deep relationships within protein families (e.g., Vosseberg et al. 2021). This protocol enables one to infer a “backbone” phylogeny of a protein family by concentrating on preselected sequences likely representing slowly-evolving members of the main clades of the family conserved across distantly related organismal lineages. Briefly (see Materials and Methods for details), we divided the sampled species into 13 groups corresponding to major eukaryotic lineages, and for each pair of groups we identified all pairs of sequences (the two sequences representing the two different groups) that had mutually minimal genetic distances calculated by the maximum likelihood method from a multiple sequence alignment. We then gathered all the sequence pairs of all the comparisons, removed redundancies, and inferred trees from the full resulting dataset (supplementary fig. 1, Supplementary Material online) or after pruning sequences from selected species to further decrease the complexity of the analysis (fig. 1; supplementary fig. 2, Supplementary Material online). This resulted in a taxonomically rich and generally well resolved final phylogeny, which enabled us to infer various aspects about the evolutionary and diversity history of the ARF family in eukaryotes.

**Fig. 1.**
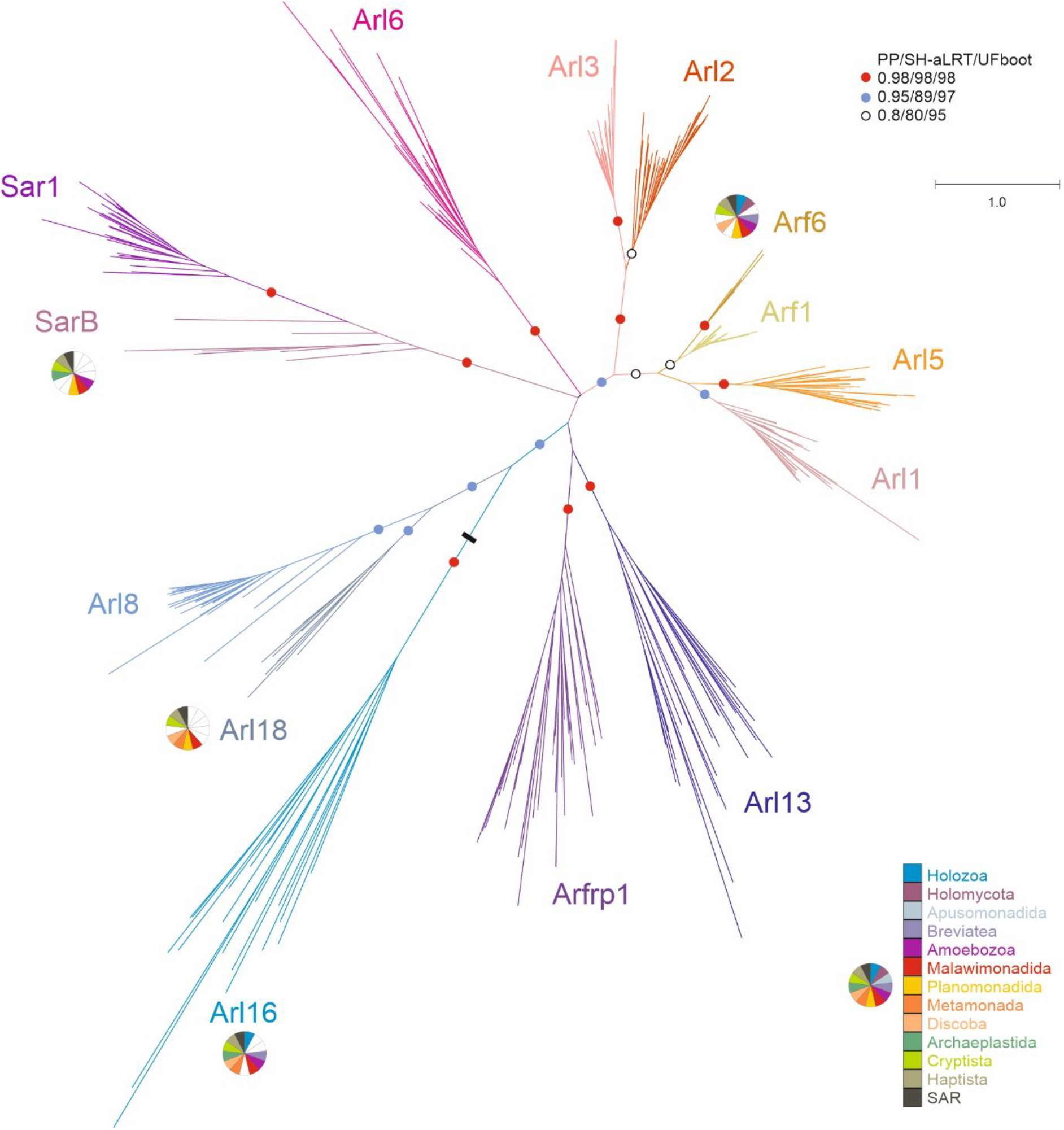
Maximum likelihood phylogenetic tree of the ARF family based on a reduced ScrollSaw dataset. The tree was inferred using IQ-TREE with LG+I+G4 model (the model selected by the program itself) based on a multiple alignment of 348 protein sequences. Brach support was evaluated with MrBayes (posterior probability, PP) and with IQ-TREE using the SH-aLRT test and the ultrafast (UF) bootstrap algorithm (both 10,000 replicates), as described under Materials and Methods. Dots at branches represent bootstrap values as indicated in the graphical legend (top right), the black bar indicates the position of the root of the tree as determined with the MAD method. The bar on the top corresponds to the estimated number of substitutions per site. The pie charts indicate the occurrence of Arf6, Arl16, Arl18 and SarB in main eukaryotic lineages (indicated by different colours explained in the graphical legend in the lower right). The remaining paralogs have ubiquitous distribution (i.e., are present in all main lineages analysed). A full version of the tree is provided in supplementary fig. 2, Supplementary Material online.

### LECA possessed an extensive array of ARF family paralogs

Dissection of the “ScrollSaw” trees indicated the existence of 13 potentially monophyletic groups (Sar1 and SarB are counted as a single putative clade for the moment, see below). Each group is represented by genes from all or a majority of the major eukaryote lineages, in all cases spanning both putative principal clades of eukaryotes (Opimoda and Diphoda) defined by the most recent hypothesis on the position of the root of the eukaryote phylogeny (Derelle et al. 2015). As such these groups all are candidates for separate ARF family paralogs differentiated before the radiation of extant eukaryotes and perhaps present in the LECA, provided that they are monophyletic (i.e. that the root of the ARF family tree lies outside of them). Our trees are inherently unrooted due to the absence of a suitable outgroup, as other GTPases, including the presumably most closely related group, SRβ, are too divergent and their inclusion into these analyses limits the resolution of the trees. Hence, to formally rule out the possibility that the root lies in any of the 13 putative clades, we employed the outgroup-independent minimal ancestor deviation (MAD) method (Tria et al. 2017), which placed the root onto a branch separating the Arl16 group from all other groups combined (fig. 1). We also note that the rooting outside any of the 13 groups implies a much simpler evolutionary scenario than a root positioned into any of the groups, so hereafter we treat the 13 groups as clades. Most of them have high statistical support (posterior probability, SH-aLRT support, and ultrafast bootstrap values greater than or equal to 0.98, 98, and 98, respectively) (fig. 1). An exception is the clade denoted Arf1 and comprising prototypical Arf sequences, but there is little doubt that it constitutes a coherent group of orthologs. The weak signal for its monophyly may stem from a very slow evolution of Arf1 sequences (apparent also from very short branches in the tree) having precluded accumulation of paralog-specific sequence features that would enable strong phylogenetic separation from the related, more rapidly evolving (and much more strongly supported) paralogs. Nevertheless, a focused analysis restricted to Arf1, Arf6, Arl1 and Arl5 allowed us to use a protein alignment with more positions and recovered Arf1 as a supported monophyletic clade (supplementary fig. 3, Supplementary Material online).

The existence of two separate clades of Arfs originated before the divergence of metazoans, fungi, and plants was hypothesized previously but not convincingly demonstrated (Li et al. 2004). We show that mammalian Arf6 has robustly supported orthologs in various protists spanning the phylogenetic breadth of eukaryotes. The existence of a separate eukaryotic Arf6 clade is further supported by comparison of intron positions in Arf genes (supplementary fig. 4, Supplementary Material online). In contrast, as expected, the mammalian Arf1-Arf5 proteins (class I and II Arfs) all cluster into the Arf1 clade. Our analyses further demonstrate that the metazoan Arl16 has orthologs present in diverse protists and thus represents a novel ancient ARF family paralog. Another previously unrecognized ancient paralog, which we propose to call Arl18, was missed because it is not represented in metazoans and has no characterized or named member. It is most closely related to Arl8, yet the separation of Arl8 and Arl18 is apparent not only from the phylogenetic analysis (fig. 1; supplementary fig. 2, Supplementary Material online) but also from their distinct exon-intron structures (supplementary fig. 5, Supplementary Material online).

Two additional ancient eukaryotic ARF family paralogs seem to exist, although they were not unambiguously supported by our phylogenetic analyses. The broader clade including Sar1 proteins and their relatives has a somewhat unusual internal structure with a strongly supported subclade, comprised of typical Sar1 proteins found in all taxa investigated, and a more basal paraphyletic group of proteins representing different Sar1-like paralogs from phylogenetically diverse protist lineages (fig. 1; supplementary fig. 2, Supplementary Material online). These are not simply divergent Sar1 orthologs, as they always co-occur with a *bona fide* Sar1 in each species analyzed, and multiple lines of evidence suggest they constitute a separate ancient paralog of their own, which we call SarB (adopting the name proposed before for a respective *Dictyostelium discoideum* representative; Week et al. 2003). Specifically, some intron positions in SarB genes are exclusive for this group and not shared with Sar1 (supplementary fig. 6, Supplementary Material online) and the functionally important Walker B motif of SarB generally exhibits a conserved tryptophan residue shared by other ARF family members and Gα proteins, as opposed to a phenylalanine residue typical for Sar1 proteins (Vetter 2014; supplementary fig. 7, Supplementary Material online). Furthermore, a ML tree with SarB sequences constrained to form a clade could not be rejected by AU test, as opposed to trees imposing topologies that would correspond to the origin of SarB genes by multiple independent duplications of Sar1 genes proper (supplementary table 2, Supplementary Material online). Hence, it is most parsimonious to interpret SarB as a *bona fide* ancient ARF family paralog different from Sar1, with the phylogenetic signal for its monophyly virtually vanished over the eons. Such a situation is not uncommon in phylogenetic analyses of families of short proteins with an inherently limited phylogenetic signal. For instance, a similar behaviour was previously observed with the highly conserved Rab1 GTPase paralog, whose undoubted monophyly was also difficult to recover (Elias et al. 2012).

The second additional potential ancient paralog, here proposed to be called Arl17, is present in various protists, certain fungi, and a single metazoan lineage, and its representative contain one to three non-identical copies of a novel conserved domain C-terminal to the GTPase domain (fig. 2; supplementary fig. 8, Supplementary Material online). The novel ~100 residue, C-terminal domain displays no discernible homology to previously described domains (even when tested by the highly sensitive HHpred searches), but occurs also in other (non-Arl17) proteins from some opisthokonts and bacteria, either as a stand-alone protein (e.g., EGF92317.1) or in combination with various non-GTPase domains (e.g., XP_004347279.1). Despite their unique domain architecture, no Arl17 sequences passed the ScrollSaw filter, hence they are absent from the tree presented in fig. 1, and although forming a clade in phylogenetic analysis, statistical support for their monophyly is lacking (fig. 2). Still, the most parsimonious interpretation of our analyses is that Arl17 is an ancient ARF family GTPase that was present already in LECA and had evolved from a duplication of the Arf1 gene, but the tendency of the GTPase domains in Arl17 proteins to be very divergent (supplementary fig. 9, Supplementary Material online) has weakened the signal for their monophyly.

**Fig. 2.**
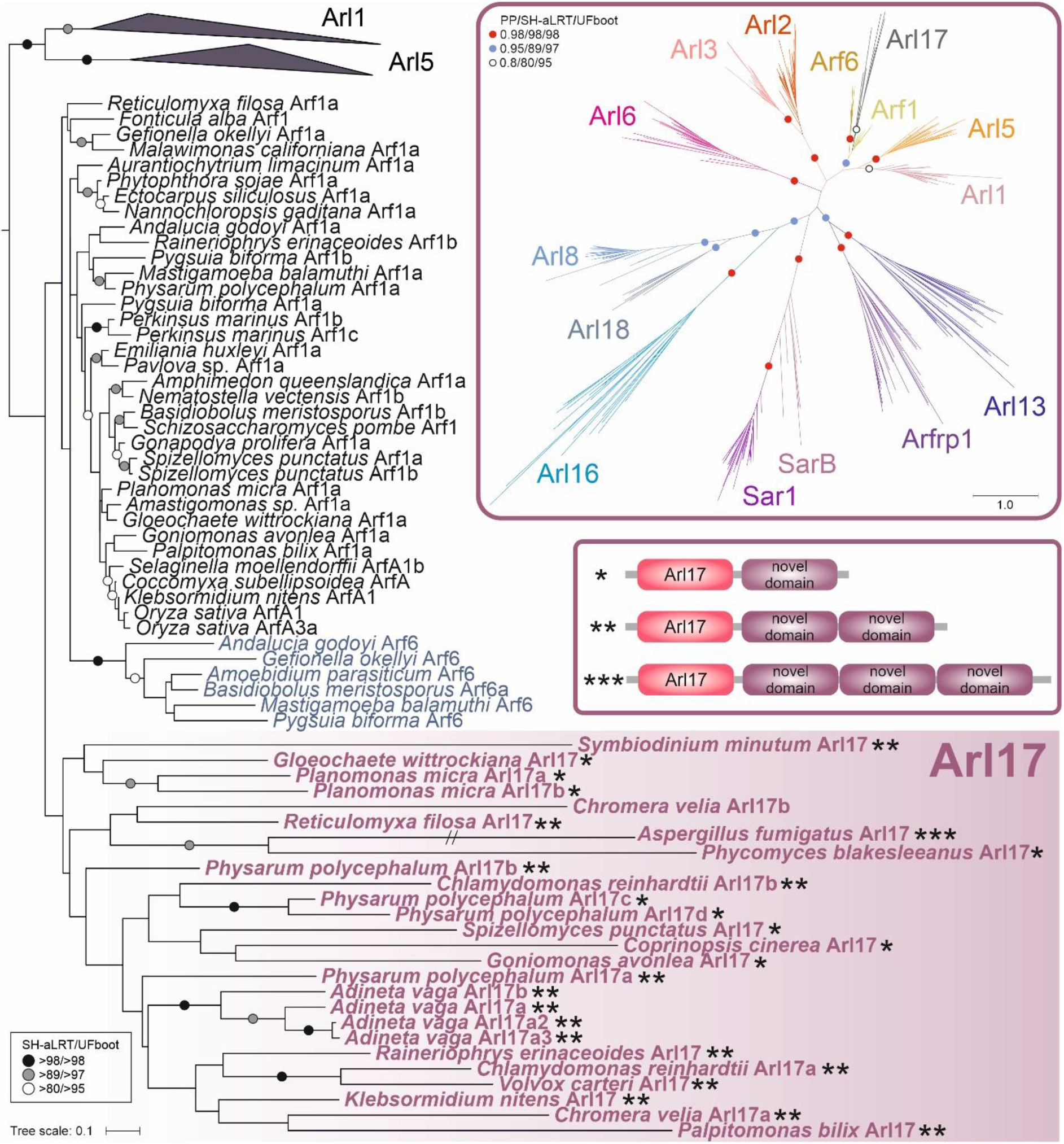
Phylogenetic analysis and domain architecture of Arl17. The tree shown is a result of a ML analysis of all Arl17 sequences and a subset of the reduced “scrollsawed” dataset restricted to Arf1, Arf6, Arl1, and Arl5 sequences (the latter two collapsed as triangles), altogether 127 protein sequences. The alignment was trimmed manually. The tree was inferred using IQ-TREE with LG+I+G4 model (the model selected by the program itself) with the ultrafast bootstrap algorithm and the SH-aLRT test (both 10,000 replicates). Dots at branches represent bootstrap values as indicated in the graphical legend (top right). The upper inset shows the ML tree inferred from a full reduced “scrollsawed” dataset combined with a subset of Arl17 sequences (picking one representative per each major eukaryote group), altogether 356 protein sequences. The tree was inferred using the same approach as the tree shown in fig. 1. The inset beneath provides a schematic representation of three different variants of the Arl17 domain architecture (correspondence to specific proteins in the tree is indicated by the asterisks). The exact architecture of the *Ch. velia* Arl17b protein could not be determine due to incompleteness of the genome assembly.

Having established the main lineages of the ARF family, we attempted to assign all other genes in our full dataset (i.e. those that were excluded by the ScrollSaw protocol) into them by considering sequence similarity scored by BLAST, comparison to lineage-specific profile HMMs by HMMER, and by targeted phylogenetic analyses. The majority of genes in our dataset could be allocated with confidence to a specific, ancient ARF family paralog, enabling us to evaluate the pattern of retention of the ancient paralogs in modern eukaryotes and to map the presumed gene losses to the eukaryote phylogeny (fig. 3; supplementary table 3, Supplementary Material online). Nevertheless, a relatively small number of genes (160 out of > 2,000 sequences) remained unclassified. A majority of these likely correspond to taxon-specific duplications of the standard ARF family members that have diverged substantially, obscuring their actual evolutionary origin. Some cases, however, may represent excessively divergent, unrecognized direct orthologs of the widespread genes. For example, several unclassified genes showed potential affiliation to Arf6, yet without significant support in phylogenetic analyses. These sequences all share one or more intron positions specific to Arf6 (supplementary fig. 4, Supplementary Material online), supporting their annotation as highly derived Arf6 genes. Future studies with a more comprehensive sampling may help resolve cases such as these.

**Fig. 3.**
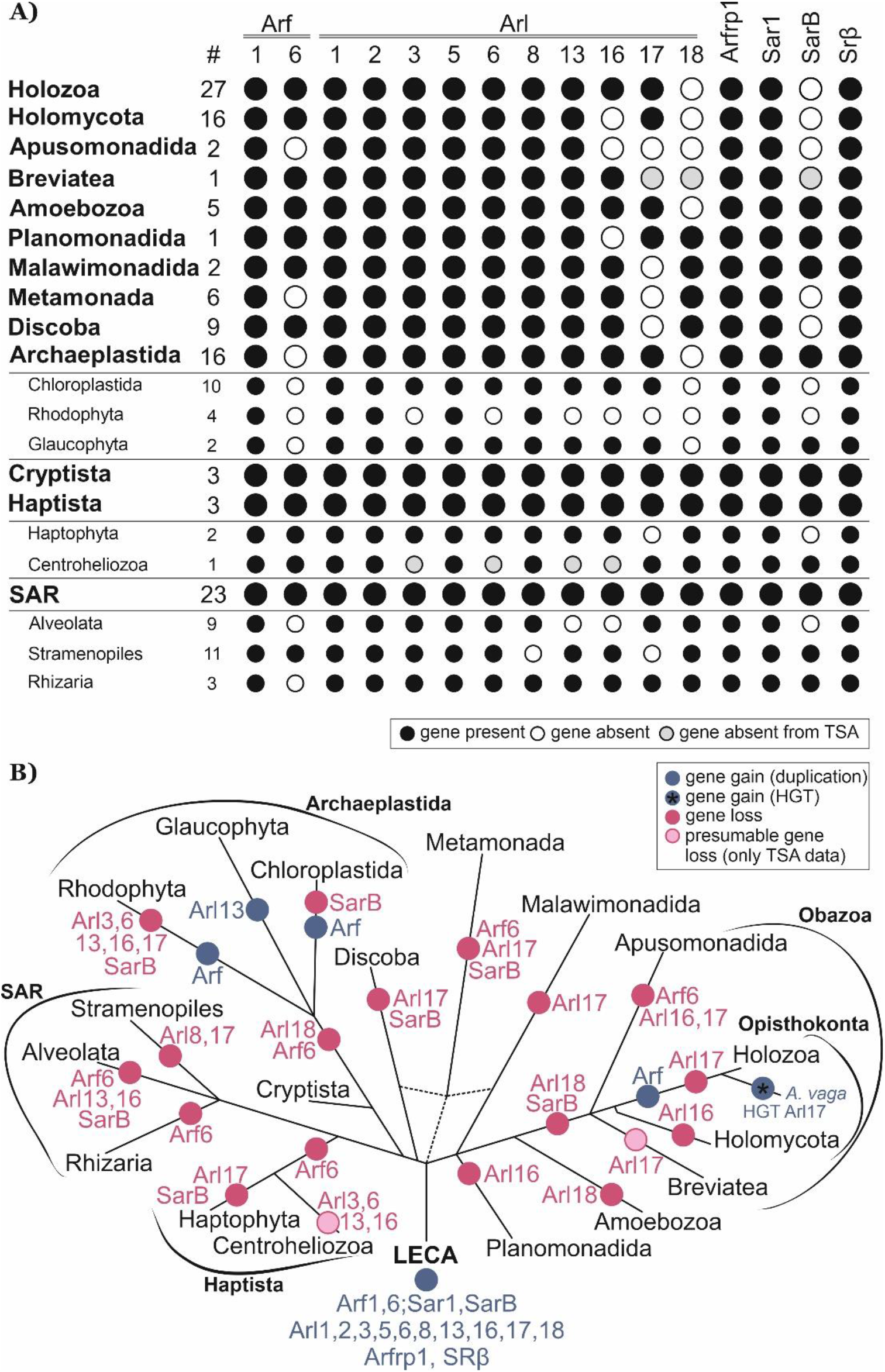
Retention of ancient paralogs of the ARF family in main lineages of eukaryotes. (**A**) Black circle: the paralog is present in at least one member of the lineage. White circle: the gene is absent from the lineage (evidenced by genome sequence data). Grey circle: the gene was not found in the transcriptome data available (lineages with transcriptome assemblies only). The hashtag (#) indicates the number of species included in the analysis. (**B**) Gene gains (blue circles) and losses (pink circles) mapped onto the eukaryote phylogeny. Only duplications specific to whole lineages listed in the picture are considered. The acquisition of Arl17 via HGT in rotifers (here represented by *A. vaga*) is indicated with a blue circle with an asterisk within.

### Complex cellular repertoire inferred from the LECA complement

The analyses presented above indicate that the LECA possessed at least 15 ARF family genes; Arf1 and 6, Arl1, 2, 3, 5, 6, 8, 13, 16, 17, and 18, Arfrp1, Sar1, and SarB. In addition, it certainly encoded SRβ, excluded from our ScrollSaw analysis (hence absent from the trees in fig. 1 and supplementary fig. 2, Supplementary Material online) due to its marked divergence from the (core) ARF family and because SRβ orthologs can be unambiguously recognized by sequence similarity. Eight of these clades (Arf1, Arl1, 2, 3, 5, and 8, Arfrp1, and Sar1) were previously recognized as likely ancient (Li et al. 2004) and the existence of orthologs of the metazoan Arl13 in protists was also noted (e.g., Miertzschke et al. 2014), although perhaps never documented by phylogenetic analyses. Our analysis thus indicates that the complement of ARF family paralogs in LECA may have been twice as big as previously identified, and further strengthens the idea that the LECA was a fully-fledged eukaryotic cell making broad use of complex molecular machinery.

The cellular functions of many of the 16 ARF family GTPases in the LECA in principle can be considered from what has been learned about their descendants in modern eukaryotes, although our present knowledge about the function of various GTPases comes from a limited number of phylogenetically biased model eukaryotes (primarily metazoans and the yeast *Saccharomyces cerevisiae*, i.e. the opisthokonts) and it is not always certain to what extent we can generalize from them to eukaryotes as a whole. In addition, each ARF family member studied in any depth in mammalian cells has been found to act in more than one pathway and typically with multiple downstream effectors (Kahn et al. 2009; Sztul et al. 2019), often making it difficult to assess which of these are ancient and which were acquired later. Finally, we recognize that any inferences about ancient functional roles relies on an assumption of functional homology across eukaryotes and an assumption of parsimonious retention of pleisiomorphic traits. From a large assessment of membrane-trafficking proteins that have been tested in model systems from across the eukaryotic tree, this assumption of functional homology appears to be justified (Klinger et al. 2016), but does warrant being explicitly named. With this caveat in mind, we summarize the key findings about the different paralogs to paint a hypothetical picture of the cellular engagement of the ARF family members in the LECA.

Most of the ARF family paralogs clearly play a role in the endomembrane dynamics. As a subunit of the receptor of the signal recognition particle, SRβ mediates co-translational import of proteins into the ER (Schwartz and Blobel 2003). Sar1 also associates with the ER and recruits subunits of the COPII coat complex to promote budding of transport vesicles from the ER (Miller and Barlowe 2010). Four paralogs – Arf1, Arfrp1, Arl1 and Arl5 – are physically and functionally associated with the Golgi/*trans*-Golgi network (TGN). One key function of Arf1 (including the metazoan Arf1 to Arf5) is to recruit different types of vesicle coats (COPI, AP-1/clathrin, AP-3) to different parts of the Golgi (Jackson and Bouvet 2014). Arl1 and Arfrp1 (confusingly called Arl3p in the yeast *S. cerevisiae*) are functionally linked, the latter shown to be critical for Arl1 recruitment to the *trans*-Golgi in both yeast and mammalian cells (Panic et al. 2003; Setty et al. 2003; Zahn et al. 2006). Arl1 recruits several effectors (e.g. golgins, arfaptins, and Arf-GEFs) to the *trans*-Golgi network (TGN) and is important for endosome-to-TGN traffic (Yu and Lee 2017). The function of Arl5 is less-well understood, but it may partly overlap with that of Arl1, as it also localizes to the *trans*-Golgi (Houghton et al. 2012), and both the fly Arl5 and the yeast Arl1 each interact with the GARP tethering complex (Panic et al. 2003; Rosa-Ferreira et al. 2015). In contrast to the Golgi localizing and acting members of the ARF family, Arf6 acts predominantly at the cell surface and endosomes to mediate endosome recycling, cell motility, and membrane extensions, which together influence cell division, lipid/cholesterol metabolism, and changes in actin dynamics (D’Souza-Schorey and Chavrier 2006; Cotton et al. 2007; Funakoshi et al. 2011; Schweitzer et al. 2011). Arl8 has been implicated in controlling lysosomal motility and traffic in metazoan cells (Khater et al. 2015). Its localization to the vacuolar membranes in *A. thaliana* (Heazlewood et al. 2007) suggests that functional association of Arl8 with the lysosomal/vacuolar compartment is ancestral and conserved.

Three paralogs, Arl3, Arl6, and Arl13 have been implicated in flagellar function (Fisher et al. 2020). Arl3 has been proposed to regulate the delivery of N-myristoylated and prenylated proteins to the cilium (Fansa and Wittinghofer 2016; Stephen and Ismail 2016). Arl6 (also called BBS3) regulates the function of the BBSome (a protein complex involved in intraflagellar transport; Mourão et al. 2014). Arl13 is involved in ciliary protein import and export, purportedly mediated by its activity as a positive regulator (guanine nucleotide exchange factor, GEF) for Arl3 (Gotthardt et al. 2015; Ivanova et al. 2017). Arl2 shares some effectors with Arl3 and is probably involved in traffic of lipidated proteins (Van Valkenburgh et al. 2001; Fansa and Wittinghofer 2016), but it has its own specific agenda, as it regulates the assembly of αβ-tubulin dimers (Al-Bassam 2017; Francis et al. 2017a; Francis et al. 2017b) and mitochondrial fusion (Newman et al. 2017).

Only a single study addressing the function of Arl16 has been published, reporting that the mammalian Arl16 inhibits the function of the RIG-I protein, involved in the defence against RNA viruses (Yang et al. 2011), but more specific functional insights are lacking. Functions for of the newly discovered paralogs SarB, Arl17, and Arl18 are completely unknown, as these paralogs are missing from all common model eukaryotes and thus represent examples of “jotnarlogs”, proteins that are present across eukaryotes, but missing in well-studied cell biological models (More et al. 2020). This adds further credence to the proposal that this is a substantial evolutionary cell biological phenomenon and highlights the gap in our understanding of the cell biology of the ARF family in eukaryotes. Nevertheless, some clues as to the function of these proteins are provided by the phylogenetic relationship to other paralogs, as relatedness within the ARF family appears to signify some level of functional similarity, despite exceptions. Indeed, the aforementioned functional aspects shared by the pairs Arl2-Arl3 and Arl1-Arl5 are reflected by close relationship of the paralogs in the pairs (fig. 1). Likewise, the related Arf1 and Arf6 paralogs, although different in terms of the intracellular localization and effectors they deploy (Jackson and Bouvet 2014), share the same class of GEFs and GTPase activating proteins (GAPs), though to a very incompletely characterized extent (Casanova 2007; Kahn et al. 2008; Sztul et al. 2019). Hence, by analogy we speculate that Arl18 may have similar functional attributes as its closest paralog Arl8 (e.g. it may likewise function in the lysosomal/vacuolar sector of the endomembrane system), and that SarB functions similarly to the canonical Sar1 protein in the secretory pathway (Sato and Nakano 2007; Melville et al. 2020). The specific relationship of Arl17 and true Arfs may be less informative concerning the function of the latter, given the unique domain architecture of Arl17 proteins and the generally divergent nature of their GTPase domains (compare the branch lengths of Arl17 sequences in the tree in fig. 2).

### Phylogenetic profiles of some ancestral eukaryotic ARF family paralogs illuminate differential simplification of endomembrane system functions in eukaryote evolution

A detailed scrutiny of the taxonomic distribution of some of the ancestral ARF family paralogs in extant eukaryotes provides interesting insights into the variation of their roles in cell functions across eukaryotes. While a hallmark of the ARF family perhaps is that members are commonly found to be active in multiple, distinct pathways in the same cells (Francis et al. 2016; Sztul et al. 2019), here we discuss their known or predicted functionalities with respect to their best known activities, recognizing the limitations that result.

Arfs (specifically the Arf1 paralog), Sar1, and SRβ are all found in every eukaryote sampled (with one exception in case of SRβ, most likely due to incompleteness of the data; supplementary table 3, Supplementary Material online), indicating that they belong to the functional core of the eukaryotic protein toolkit. Nearly ubiquitous is Arl2, being absent only from *Entamoeba histolytica*. Inspection of genomes of other *Entamoeba* species suggest that Arl2 loss is not an artefact and predates the radiation of the genus. Given the role of Arl2 in the assembly of tubulin dimers and in mitochondrial fusion (Francis et al. 2016), its absence in *Entamoeba* may be related to a unique combination of traits of this taxon including divergent tubulin sequences and a highly reduced microtubular cytoskeleton (Roy and Lohia 2004; Meza et al. 2006), and a simplified mitochondrion (i.e., a mitosome; Makiuchi and Nozaki 2014).

Five of the ancestral paralogs functionally linked to the endomembrane system (based on data from model eukaryotes) show various degrees of patchiness in their occurrence (fig. 3A; supplementary table 3, Supplementary Material online). Arl1, Arl5, and Arfrp1, all associated with the Golgi apparatus, have been preserved in all main eukaryote lineages sampled, but have been lost from some more terminal branches. Arl1 is missing from the fission yeast (*S. pombe*), diplomonads, and some apicomplexans. Arfrp1 is absent from the same set of species plus two more (the highly reduced endosymbiotic kinetoplastid *Perkinsela* sp. CCAP 1560/4 and the tiny green alga *Micromonas commoda*). The similar patterns of loss of these two GTPases may reflect the fact that they were shown to work in the same functional cascade (see above). How Arl1 functions in the absence of Arfrp1 in *Perkinsela* or *Micromonas* remains an open question but may reflect the multiplicity of pathways each GTPase may influence and the potential differences in their means of localization and activation. Arl5 is missing from many more eukaryotes, including even some metazoans (e.g., the flatworm *Schmidtea mediterranea*). A minimum of 20 independent losses of Arl5 is required to explain its distribution in our dataset (supplementary table 3, Supplementary Material online), suggesting that this GTPase is a less critical component of the basic infrastructure of the eukaryotic cell. In accord, disruption of the Arl5 gene in *Drosophila melanogaster* does not alter the fly’s viability or fertility (Rosa-Ferreira et al. 2015). Arl5 is closely related to Arl1 and the two GTPases may share some effectors (see above). It is thus possible that Arl5 loss is facilitated by partial functional redundancy with Arl1.Similar to Arl5, the distribution of Arl8 in extant eukaryotes has been shaped by multiple (at least 14) independent losses, including one in the lineage leading to the main eukaryotic taxon Stramenopiles (fig. 3B; supplementary table 3, Supplementary Material online). Comparison of phylogenetic profiles of Arl8 and the related uncharacterized paralog Arl18 reveals that the former paralog has been retained more frequently than the latter, but in a few taxa (e.g., stramenopiles) Arl18 occurs in the absence of Arl8 (fig. 3A; supplementary table 3, Supplementary Material online). It would be interesting to investigate whether a level of functional redundancy might allow Arl18 to have taken over some of the Arl8 functions in these organisms. The presence of both Arl8 and Arl18 in model systems like *Tetrahymena thermophila* and *Trypanosoma cruzi* (supplementary table 3, Supplementary Material online) provides a chance that functional dissection of these closely related paralogs is possible.

The patchy distribution of Arf6 is somewhat surprising, at least in part because it contrasts with the near universal distribution of Arf1 paralogs. While Arf6 is perhaps most commonly associated with endocytosis and plasma membrane dynamics (see above) we speculate that perhaps it is its role in pericentriolar localization of specific subsets of recycling endosomes that are required for midbody formation and abscission (Fielding et al. 2005; Wilson et al. 2005; Turn et al. 2020) that vary amongst species. The nature and composition of centrioles, as well as associated components are known to vary, including losses or differences in Archaeplastida and SAR (Nabais et al. 2020).

The unexpected discovery of the sporadically distributed, yet potentially ancestral SarB paralog (figs 1 and 3A; supplementary table 3, Supplementary Material online) raises an interesting possibility of a specific elaboration of the ER function in the LECA lost for some reason(s) by most major eukaryotic groups. Direct functional characterization of SarB in suitable model organisms is necessary before the causes behind the retention/loss pattern of the gene may be understood. However, it is interesting to compare SarB with the recently uncovered complexity of the ancestral set of paralogs of the COPII coat complex, including the Sec24III paralog as patchily distributed as SarB (Schlacht and Dacks 2015). The phylogenetic profiles of SarB and Sec24III do not overlap well (e.g., SarB is missing from Chloroplastida and Sec24III is absent from diatoms), so we are not suggesting a specific functional link between these two proteins. Nevertheless, the existence of both proteins implies the existence of an interesting degree of variation in the COPII vesicle formation at the ER in different eukaryotes.

### Arl17 provides a rare example of horizontal transfer of a Ras superfamily gene

The newly recognized functionally uncharacterized Arl17 group of ARF family protein is unusual not only because of its unique domain architecture (fig. 2), but also due to its very patchy taxonomic distributions (fig. 3A; supplementary table 3, Supplementary Material online). Based on our current sampling, Arl17 is completely missing from several major eukaryotic clades (Malawimonadida, Metamonada, Discoba, Stramenopiles, Haptophyta, and Rhodophyta), whereas its occurrence in the other groups is typically sporadic. Particularly interesting is identification of a group of four closely related Arl17 homologs in the rotifer *A. vaga* (Fig. 2; supplementary table 3, Supplementary Material online), which is the sole representative of the densely sampled Holozoa clade possessing Arl17 (supplementary table 3, Supplementary Material online). Transcriptome data from *A. vaga* relatives indicate that Arl17 is not restricted to a single rotifer species (data not shown), ruling out contamination in the *A. vaga* genome data. Hence, the isolated occurrence of Arl17 in a rotifer lineage strongly indicates gain via horizontal gene transfer (HGT) from a protist or fungal lineage, with subsequent gene duplications (at least partly accounted for by tetraploidy of the *A. vaga* genome, see above). Indeed, analyses of rotifer genomes revealed propensity of these peculiar microscopic animals for gene gain from various sources, and three of the four *A. vaga* Arl17 paralogs were included in the list of HGT candidates in the *A. vaga* genome (Flot et al. 2013). To our knowledge, this is the first convincing case of a eukaryote-to-eukaryote HGT in the whole Ras superfamily. Even though phylogenetic analysis of the Arl17 GTPase domain did not shed light on the origin of rotifer’s Arl17 (fig. 2), a specific relationship to Arl17 proteins from *Physarum polycephalum* is suggested by a phylogeny inferred for the different copies of the C-terminal novel domains (supplementary fig. 10, Supplementary Material online), suggesting that rotifers acquired Arl17 from an amoebozoan.

### Expansion of the ARF family in Holozoa

Given the prominent position of metazoan model systems (humans, *Mus musculus, D. melanogaster*, *Caenorhabditis elegans*) in research on the ARF family, we carried out a separate analysis concentrating on the family members in widely sampled representatives of Metazoa and their closest protist relatives, together constituting the taxon called Holozoa. Analogously to our eukaryote-scale ScrollSaw analysis described above, we compared 18 groups of sequences corresponding to the main holozoan lineages (phyla). This approach narrowed our original holozoan dataset of nearly 550 sequences to ~320 sequences and phylogenetic analysis of this reduced dataset revealed a set of strongly supported clades that provided a basis for defining ARF family paralogs conserved across the main holozoan or metazoan lineages (fig. 4). All ancient eukaryotic paralogs represented in this taxon, except Arf1, form supported clades (note that Arl17 failed to pass the ScrollSaw step as it is present only in rotifers). Furthermore, six additional groups could be identified based on this analysis, namely Arf4 (class II Arf), Arl4, 10, 15, 19 and TRIM23. Most of them are named according to previously annotated vertebrate genes (Gillingham and Munro 2007). An exception is a novel group, here named Arl19, which is not a resolved clade, but seems to represent a coherent evolutionary lineage based on additional evidence (see below). Analysis of intron positions in a subgroup of ARF family genes corresponding to Arfs and their closest relatives supported the delimitation of the main groups, but also suggested that several sequences initially annotated as Arf1 (based on BLAST searches) may constitute a novel conserved group in unicellular holozoans and several invertebrate lineages (supplementary fig. 11, Supplementary Material online). Specifically, this group is characterized by three unique intron positions, and a focused phylogenetic analysis supported its monophyly and separation from Arf1 and other clades (supplementary fig. 12A, Supplementary Material online). We thus named this novel clade Arl20.

**Fig. 4.**
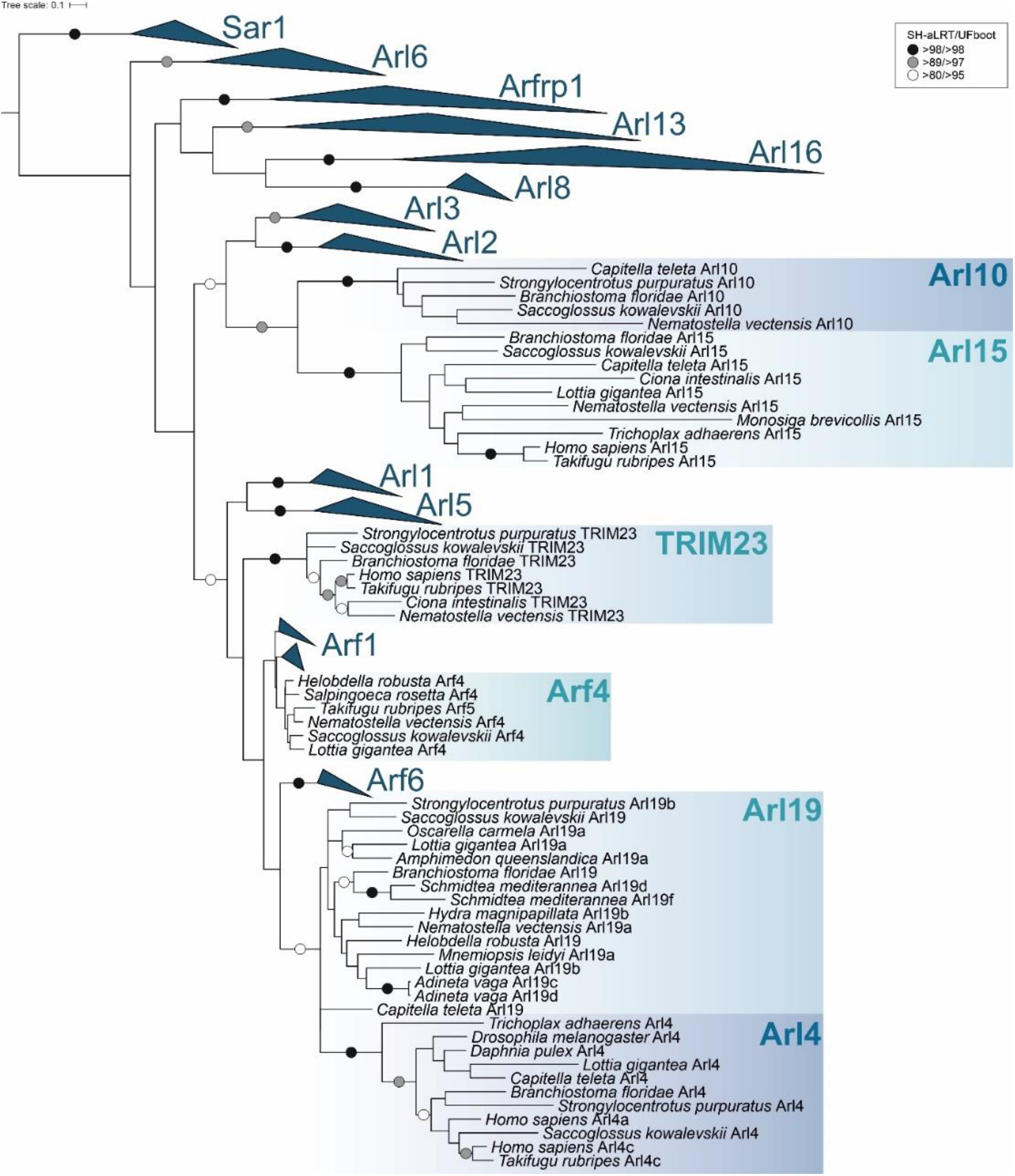
Maximum likelihood phylogenetic tree of the ARF family based on a ScrollSaw dataset in Holozoa. The tree was inferred using IQ-TREE with LG+I+G4 model (the model selected by the program itself) from a multiple alignment of 323 protein sequences with the ultrafast bootstrap algorithm and the SH-aLRT test (both 10000 replicates), as described under Materials and Methods. Dots at branches represent bootstrap values as indicated in the legend shown in the bottom left. Eukaryotic ancestral paralogs are collapsed as triangles.

Establishment of novel ARF lineages provided a basis for the assignment of sequences excluded by the ScrollSaw protocol by the same approach as described for the ancient eukaryotic paralogs. Moreover, inspection of the exon-intron structures facilitated assignment of some of the problematic genes. For example, *Takifugu rubripes* harbours several standard Arfs and one additional Arf-like paralog (TruArf4L in supplementary table 1, Supplementary Material online) with an almost equal similarity to the Arf1 and Arf4 groups. Both phylogenetic and HMMER-based analyses were inconclusive concerning the origin of this gene, but the exon-intron structure of TruArf4L exhibits the pattern typical to the Arf4 group (supplementary fig. 11, Supplementary Material online), supporting annotation of this gene as a divergent representative of the Arf4 group. Combining such different forms of evidence allowed us to annotate the majority of sequences, to establish the phylogenetic distribution of the main groups, and to map their origins and losses onto the holozoan phylogeny (fig. 5; supplementary table 3, Supplementary Material online).

**Fig. 5.**
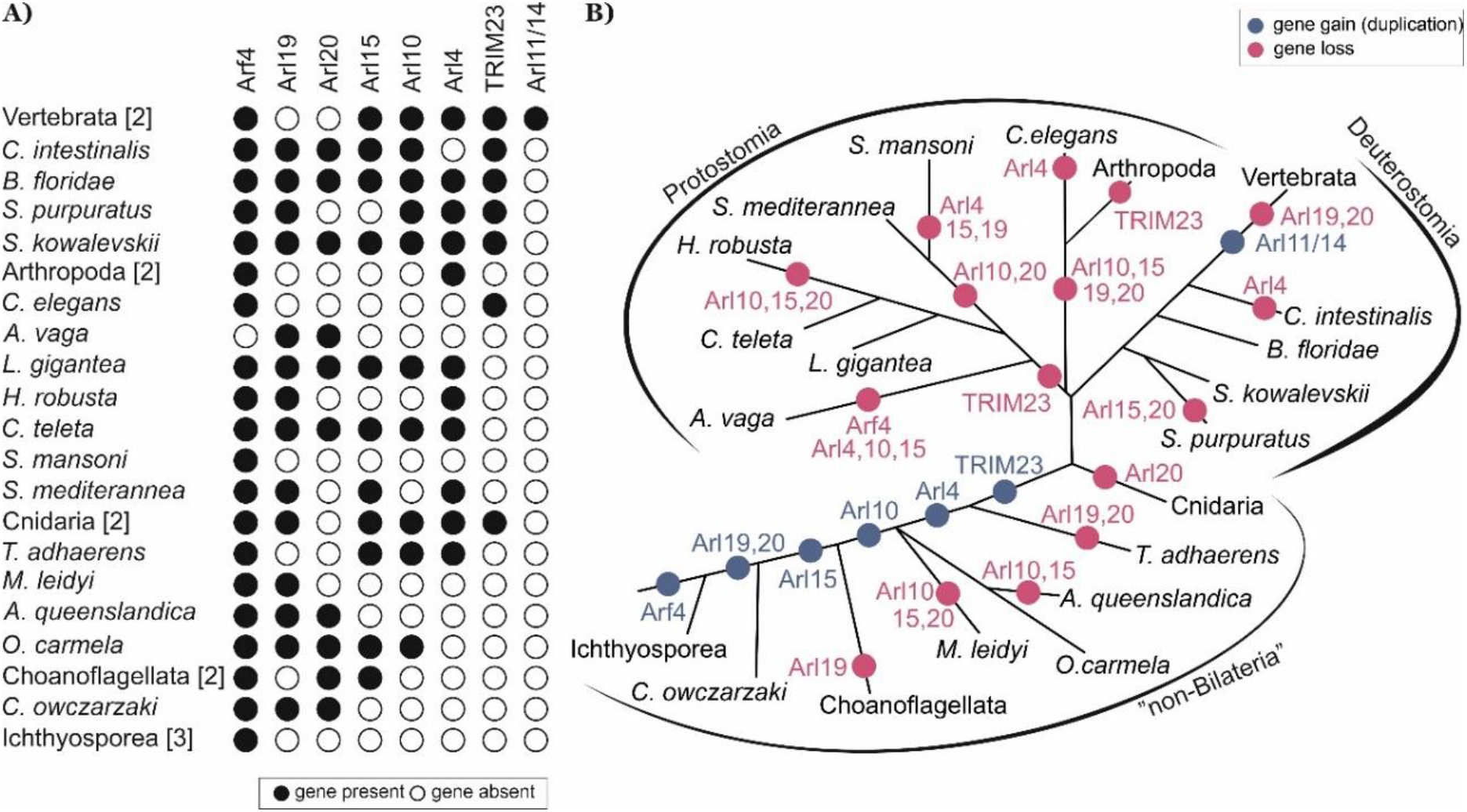
Retention of lineage-specific paralogs of the ARF family in main lineages of Holozoa. (**A**) Black circle: the paralog is present in at least one member of the lineage; white circle: the gene is absent from the lineage (evidenced by genome sequence data). Species with identical distribution are collapsed into higher taxa with the number of species indicated in the square brackets. (**B**) Gene gains (blue circles) and losses (pink circles) mapped onto the holozoan phylogeny.

Altogether we could recognise seven groups that apparently originated after the split of the holozoan lineage from their relatives (Holomycota), that is in the holozoan stem itself (Arf4), at a later step but still before the divergence of Metazoa and their sister group choanoflagellates (Arl15, 19, 20), in the metazoan stem (Arl10), or after the divergence of the deepest metazoan phyla (Arl4, TRIM23). This stepwise build-up of complexity of the ARF family (fig. 5B) contrasts with a somewhat different evolutionary pattern documented for the Rab family, which experienced a wave of expansion concentrated in the metazoan stem lineage (Elias et al. 2012). The novel ARF family members in Holozoa apparently emerged by modification of duplicated copies of specific ancient eukaryotic paralogs, although the exact sources may be difficult to determine. Sequence similarity and phylogenetic analysis (fig. 4) point to the Arl2/3 clade as the most likely cradle of Arl10 and 15, but the position of these two paralogs is unstable in different phylogenies (e.g. supplementary fig. 12B, Supplementary Material online). Evidence is more solid for the origin of Arf4, Arl4, 19, 20 and TRIM23, suggesting these are offshoots stemming from Arf1/6-like ancestors (fig. 4; supplementary fig. 12, Supplementary Material online).

A common origin of Arf1 and Arf4 groups was already reported (Li et al. 2004; Manolea et al. 2010), but our analysis placed this event before the divergence of ichthyosporeans to the common ancestor of Holozoa (fig. 5B), which probably possessed Arf1, Arf4, and Arf6 as single-copy genes. While Arf4 and Arf6 seem to duplicate only sporadically, Arf1 is often present in more than one copy, suggesting a high propensity for duplication; this tendency is in fact seen for eukaryote lineage in general (supplementary table 3, Supplementary Material online). Phylogenetic analyses usually do not recover Arf1 and Arf4 as supported monophyletic clades (e.g., fig. 4), which is probably a result of their high sequence similarity reflected also in partial functional overlap of Arf1 and Arf4 (Jackson 2014; Jackson and Bouvet 2014). However, their separation is obvious from the comparison of the exon-intron structures of the respective genes (supplementary fig, 11, Supplementary Material online).

Two more holozoan or metazoan GTPase groups are likely evolutionarily derived from the ancestral Arf1 gene, yet have diverged to the point it seems inappropriate to call them “Arfs”. One is Arl20, a previously unrecognized group of genes sharing three specific intron positions (supplementary fig. 11, Supplementary Material online). Their relationship to Arf1 cannot be conclusively inferred from our phylogenetic analysis (supplementary fig. 12A, Supplementary Material online), but an intron position shared with Arf1 (and Arf4) and outcomes of similarity searches support this hypothesis. TRIM23 (also called ARD1) is an unusual protein including not only the GTPase domain, but also a block of domains characteristic for the TRIM family (RING-type E3 ubiquitin ligase, a tandem of BBbox domains, and the BBC domain forming a coiled-coil) at the N-terminus. The GTPase domain is highly similar to true Arfs (Vichi et al. 2005) and its specific relationship to Arf1 is obvious from the virtually identical exon-intron structure (of the gene part encoding the GTPase domain; supplementary fig. 11, Supplementary Material online).

Arl4 and the Arl19 group newly recognized here constitute a sister group to Arf6 in our trees (fig. 4; supplementary fig. 12A, Supplementary Material online). While Arl4 forms a highly supported monophyletic group, its placement disrupts the monophyly of Arl19, perhaps due to an insufficient phylogenetic signal that would unite all Arl19 sequences in the analyses. The origin of Arl4 and Arl19 from Arf6 is conceivable and there are also potential functional links between Arl4 and Arf6; e.g., mammalian Arl4 proteins can recruit the Arf6 GEFs cytohesins to the plasma membrane (Hofmann et al. 2007) and each GTPase can influence actin dynamics (Cotton et al. 2007; Li et al. 2007; Patel et al. 2011). The exon-intron structure of Arl4 and Arl19 are not helpful in unveiling their origin. Only a minority of Arl4 genes contain introns, the intron positions are not conserved between Arl4 genes, and do not match the rest of examined Arf genes (supplementary fig. 11, Supplementary Material online). This suggests that Arl4 may have originated through retroposition (Kaessmann et al. 2009), that is by integration of a reverse-transcribed mRNA into the genome of an early metazoan, with the few non-conserved introns gained secondarily and independently in different metazoan lineages. The exon-intron structure of Arl19 is rather puzzling, as several genes share an intron with Arf1 (supplementary fig. 11, Supplementary Material online), but the whole clade branches off close to Arf6 (fig. 4).

In addition to the aforementioned novel ARF family members broadly conserved across Holozoa or Metazoa, various metazoan lineages exhibit still other novelties suggesting further functional elaboration. Here we focus on vertebrates. First, the vertebrate ARF family complement has been expanded by duplications of Arf1 and Arf4, yielding the well-known two groups of paralogs (Arf1, 2, 3 versus Arf4 and 5). Together with multiple duplications of Arl4, vertebrates are thus endowed with a battery of lineage-specific paralogs that are generally highly similar in sequence and (presumably) function (supplementary tables 1 and 3, Supplementary Material online). Second, vertebrates have experienced duplication of the Arl10 gene inherited from their invertebrate ancestor, giving rise to two in-paralogs that diverged from each other to such an extent that they were not initially recognized as closely related and which is reflected in their different names: Arl9 and Arl10 (supplementary fig. 12B, Supplementary Material online). Finally, vertebrates encode two divergent ARF family members of a common origin, called Arl11 and Arl14, that seems to have evolved by duplication and divergence from Arl4 (fig. 5; supplementary fig. 13, Supplementary Material online). The functional significance of these novelties is unclear, owing to limited knowledge of the function of the respective proteins in any vertebrate species including humans. It is, however, important to stress that the vertebrate ARF family complement has been sculpted also by gene loss, as vertebrates (in contrast to their sister group tunicates represented in this study by *Ciona intestinalis*) lack Arl19 and Arl20 (fig. 5).

### The emergence of other major eukaryotic clades was accompanied by limited evolutionary novelty in the ARF family

Given the identification of multiple novel ARF family paralogs in Holozoa/Metazoa, we also applied the ScrollSaw protocol to other eukaryote groups to uncover possible lineage-specific innovations. Interestingly, while gene duplications specific to terminal organismal lineages are common in the ARF family, only three higher-level taxa – rhodophytes, glaucophytes, and Chloroplastida – seem to have evolved novel family members by gene duplication in their stem lineages (fig. 3B; supplementary table 4, Supplementary Material online). The genome of red algal ancestors underwent massive reductive evolution (Yoon et al. 2017), which is reflected also by their highly reduced set of Rab GTPases (Petrželková and Eliáš 2014) as well as of ARF family proteins (fig. 3; supplementary table 3, Supplementary Material online). Somewhat opposite to this trend, a novel ARF family member, here denoted ArlRhodo, is shared by distantly related rhodophyte taxa and apparently emerged before the radiation of the whole group. Their origin remains elusive, as the phylogenetic analysis placed ArlRhodo as a separate clade of the ARF family with no specific affinities to any of the ancestral clades (fig. 6A). By contrast, the glaucophyte innovation, in fact represented by multiple paralogs in individual glaucophyte species, can clearly be traced as a highly divergent offshoot of Arl13 (supplementary fig. 14, Supplementary Material online).

**Fig. 6.**
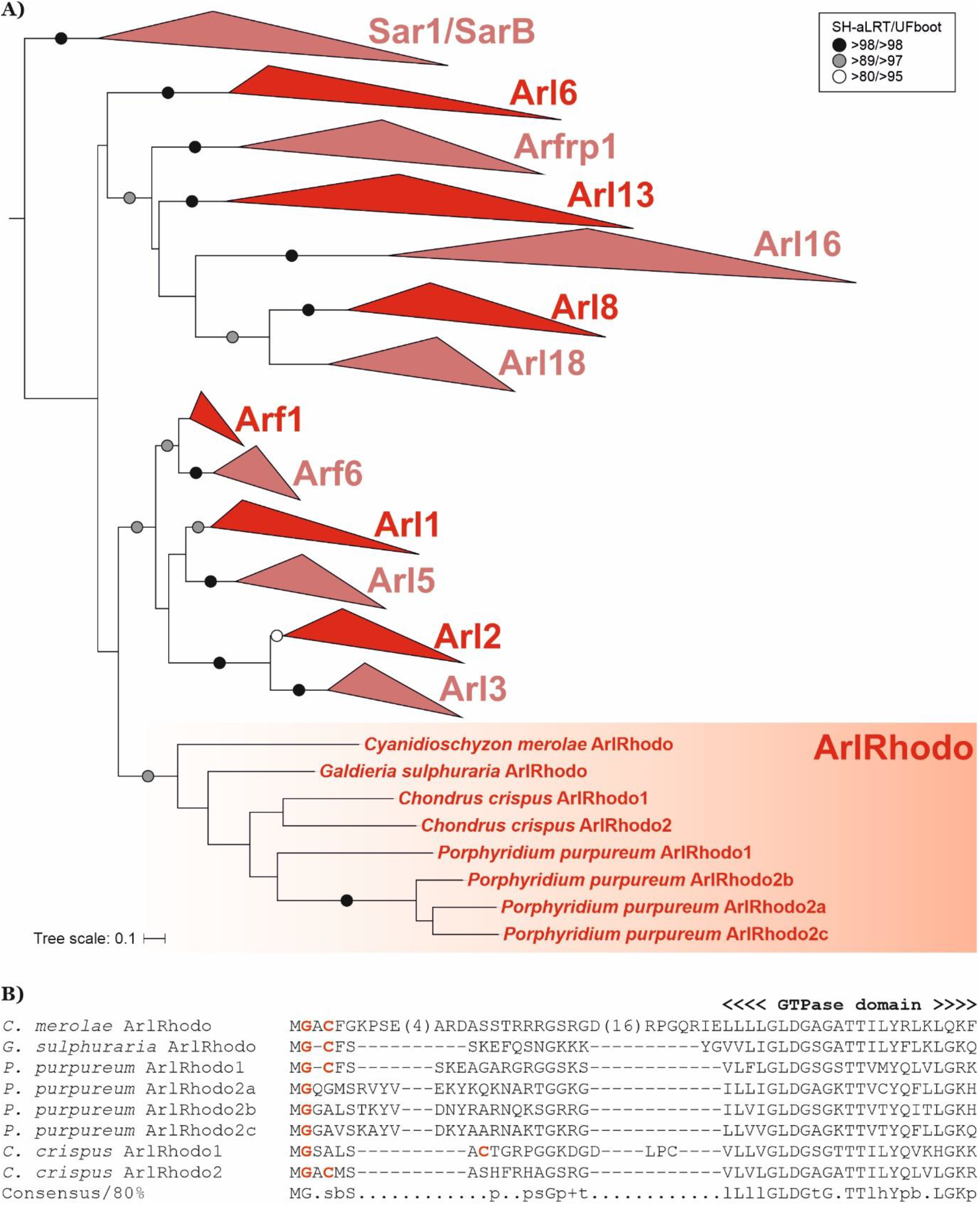
ArlRhodo, a novel ARF family member specific for red algae. (**A**) Phylogenetic analysis of the ArlRhodo group. The tree shown is a result of a ML analysis of all ArlRhodo sequences and the reduced “scrollsawed” dataset (altogether 356 sequences). The tree was inferred using IQ-TREE with LG+I+G4 model (the model selected by the program itself) with the ultrafast bootstrap algorithm and the SH-aLRT test (both 10,000 replicates). Dots at branches represent bootstrap values as indicated in the graphical legend (top right). (**B**) N-terminal region of ArlRhodo proteins with the characteristic configuration of glycine and cysteine residues (highlighted in red) predicted to be N-myristoylated and S-palmitoylated, respectively.

The only previously documented innovation of the ARF family specific for a major eukaryotic group other than metazoans is the plant ArfB (Vernoud et al. 2003). It was proposed to be an Arf6 ortholog (Li et al. 2004), and indeed our phylogenetic analysis places ArfB as sister group to Arf6 (supplementary fig. 15, Supplementary Material online). However, this topology is not statistically supported and can be an artefact resulting from the apparently rapid initial evolution of the ArfB gene reflected by the long stem branch subtending the ArfB subtree. Moreover, ArfB genes share one intron position with Arf1, but none with Arf6 (supplementary fig. 4, Supplementary Material online). Hence, we leave the origin of the ArfB group as unresolved. This not withstanding, the timing of the ArfB emergence coincides with a duplication of the ARF GEF BIG in the Chloroplastida (Pipaliya et al. 2019). The duplication of the ArfB paralogs in embryophytes also coincides with the duplication of GBF1 proteins in that same lineage. As both of these GEFs act on Arf1-derived paralogs in metazoans at least, this lends itself to the hypothesis that ArfB is derived from Arf1. It raises the further speculation that one of the BIG duplicates acts specifically on ArfB in green algae and suggests that the ArfB, BIG, and GBF1 duplicates should all be included in any activity assays aimed at understanding how this network functions in plant cells.

### Extensive molecular tinkering in the evolution of membrane attachment mechanisms in the ARF family

It is currently understood that a large fraction of ARF family members act within endomembrane traffic pathways through their actions on the surface of source membranes (Gillingham and Munro 2007). This necessitates specific, and (typically) transient, membrane attachment, typically relying on specific PTMs, employed by different ARF family members. Our analyses illuminate the origins of the previously described means of membrane association, but also finds evidence consistent with diversity in the mechanisms involved in membrane association (summarized in fig. 7A-F).

**Fig. 7.**
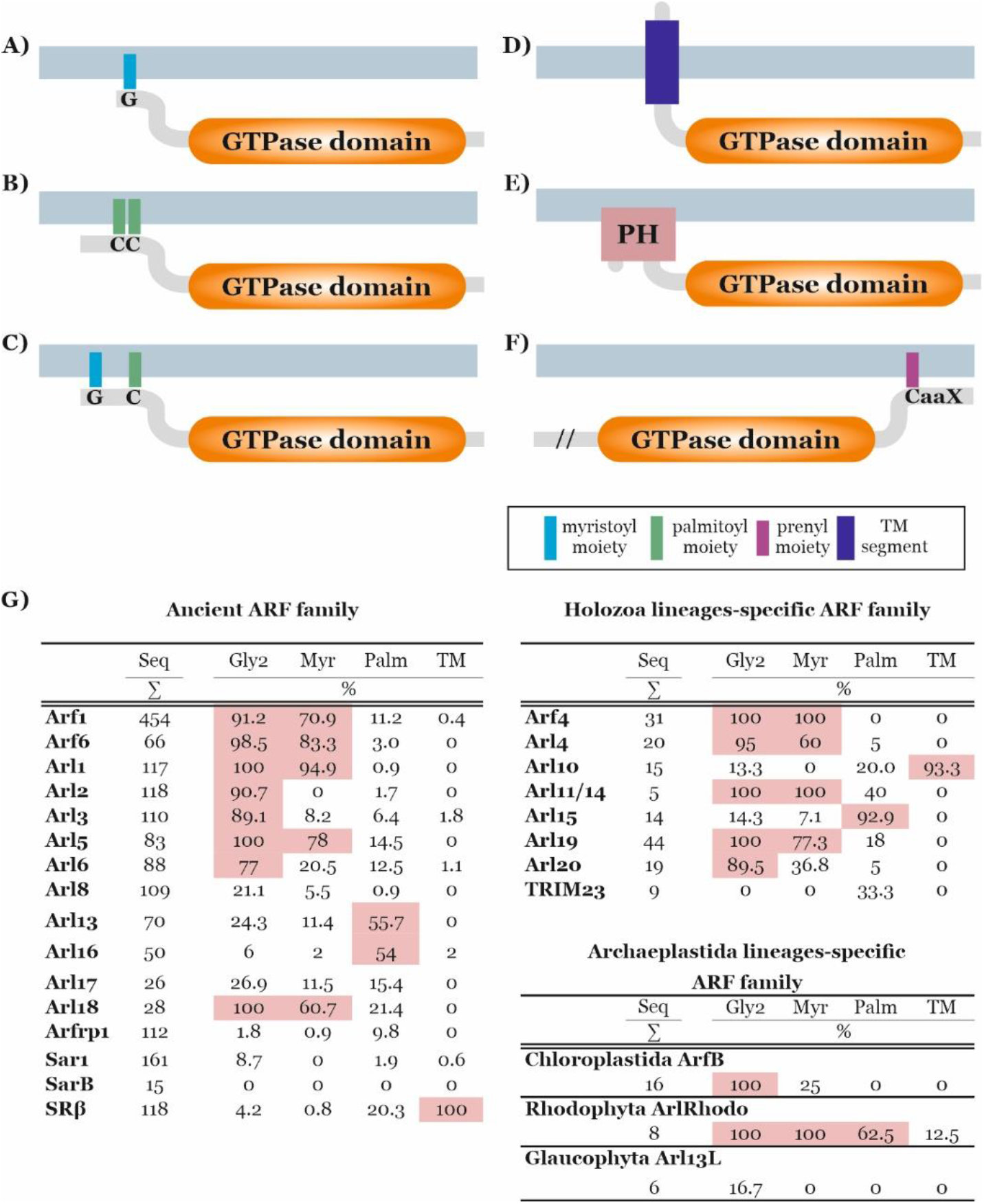
Membrane attachment mechanisms of ARF family proteins. Examples of different broadly conserved mechanisms of membrane attachments of ARF family members are depicted. (**A**) N-terminally myristoylated glycine residues, common for Arfs and several Arf-like proteins. (**B**) One or two S-palmitoylated cysteine residues near the N-terminus, typical for Arl16 and also common in Arl13. (**C**) N-terminally myristoylated glycine residue coupled with S-palmitoylated cysteine residue near the N-terminus, typical for ArlRhodo. (**D**) N-terminal transmembrane region, typical for SRβ and Arl10. (**E**) N-terminally accreted PH domain, present in divergent Arf-like proteins in kinetoplastids and choanoflagellates. (**F**) Prenylation motif (CaaX) at the C-terminus of certain eustigmatophyte-specific ARF family members (characterized also by a long N-terminal extension, in the figure marked with “//”). supplementary table 1, Supplementary Material online lists all identified ARF family proteins predicted to be N-myristoylated or S-palmitoylated, or to contain a transmembrane region or PH domain. (**G**) Summary of the results of prediction of N-myristoylation, S-palmitoylation and presence of the transmembrane (TM) region in particular subgroups of the ARF family. For each subgroup (group of orthologs), the number of sequences (Seq) and the percentages of sequences with glycine residues at the second position (Gly2), sequences predicted as N-myristoylated (Myr), sequences predicted as S-palmitoylated on at least one cysteine residue (Palm), and sequences with predicted transmembrane region(s) (TM) are given. Values above 50% are highlighted in pink. For complete data see supplementary tables 1 and 5, Supplementary Material online. These predictions were done as described under Materials and Methods.

N-terminal myristoylation (N-myristoylation) is the most common lipid modification mediating the reversible membrane attachment of ARF family proteins (Kahn et al. 1988; Liu et al. 2009). Two necessary prerequisites for N-myristoylation are the glycine residue at the second position of the protein and specific sequence motif downstream that is recognised by the myristoyl transferase catalysing the addition of the myristate moiety to the N-terminal glycine (Duronio et al. 1991; Resh 1999). Once acted upon by N-myristoyl transferase, the myristate group is attached through an amide bond that is permanent for the life of the protein. Reversibility in membrane association is tightly linked to the activation status of the ARF family protein, as the myristoylated N-terminal α-helix is accommodated in a hydrophobic channel when the protein is inactive (GDP-bound) but becomes solvent exposed in response to activation (GTP-binding), resulting in its propensity to bury the freed myristate in a lipid bilayer (Pasqualato et al. 2002; Seidel et al. 2004; Liu et al. 2009; Liu et al. 2010).

Using dedicated bioinformatic tools (see Materials and Methods), we predicted this post-translational modification for the majority of the proteins representing the ancestral eukaryotic paralogs Arf1, Arf6, Arl1, and Arl5 (fig. 7G; supplementary tables 1 and 5, Supplementary Material online), in keeping with previous experimental data from yeast and mammalian proteins (Kahn et al. 1988; D’Souza-Schorey and Stahl 1995; Lee et al. 1997; Lin et al. 2002). Virtually all Arf6, Arl1 and Arl5 proteins possess the conserved glycine residue at the second position, and the negligible minority of those not predicted as N-myristoylation targets may be false negatives. From almost 450 Arf1 genes investigated, 40 do not possess the expected glycine residue and cannot be modified by myristoylation in a standard manner. We note that a recent study found N-myristoyltransferase capable of acylating lysine in the third position (Dian et al. 2020), though the predicting algorithms employed here did not consider this possibility. Regardless, only three of the 40 Arf1 proteins without a myristoylatable glycine have a lysine residue at the third position. All of them are accompanied by two or more Arf1 genes that are N-myristoylated in the given organism (supplementary table 1, Supplementary Material online), so they apparently represent lineage-specific paralogs with a changed behaviour towards membranes. The newly recognised Arl18 paralog, though not closely related to the previous four paralogs, also is predicted to be ancestrally myristoylated, as all genes contain a glycine residue at the second position and the majority of them are predicted as N-myristoylated (fig. 7G; supplementary tables 1 and 5, Supplementary Material online). Interestingly, the Arl18 sister group Arl8 seems to ancestrally lack glycine at the second position (fig. 7G; supplementary table 1, Supplementary Material online) and the only putatively N-myristoylated Arl8 can be found in rhizarians, suggesting secondary acquisition of the myristoylation motif in this lineage. The majority of Arl2 and Arl3 proteins do harbour a glycine residue at the second position, but N-terminal myristoylation is predicted only for a few Arl3 proteins (fig. 7G; supplementary tables 1 and 5, Supplementary Material online) and these may be false positives, considering the experimental evidence for the lack of N-myristoylation in representative Arl3 proteins (Sharer et al. 2002; Setty et al. 2004). The Arl6 group is clearly heterogeneous, including members that certainly are not myristoylated as well as members that likely have this modification. Thus, the evolutionary course leading to the distribution of N-myristoylation in different ARF family members is not always clear. One possibility is an early origin of this modification in an ancestor of all the clades with N-myristoylated members, followed by its multiple secondary losses. However, multiple independent acquisitions is certainly a likely, and mutually non-exclusive, alternative.

S-palmitoylation (i.e., addition of a palmitoyl moiety to one or more cysteine residues) also mediates protein association with membranes, though unlike N-myristoylation there are enzymes capable of reversing this acylation making it a more transient modification (Zhou and Cox 2014). We again employed a suite of dedicated algorithms to predict the presence of this modification in ARF family members, as described under Materials and Methods. Arl15 proteins typically harbour several N-terminal cysteine residues, usually predicted as S-palmitoylated (supplementary fig. 16, Supplementary Material online), and approximately half of the Arl13 and Arl16 sequences analysed also contain one or more putative S-palmitoylated cysteine residues in their N-terminal region (fig. 7G; supplementary tables 1 and 5, Supplementary Material online). S-palmitoylation of Arl13 from *C. elegans* and mammals has been confirmed experimentally and demonstrated as crucial not only for the proper localization of the proteins, but also for stability and function (Cevik et al. 2010; Roy et al. 2017). In a few cases, such as in the red algae-specific paralog ArlRhodo, S-palmitoylation seems to accompany N-myristoylation (figs 6B and 7G; supplementary table 1, Supplementary Material online), similar to various other proteins, including GTPases (e.g., some Gα proteins; Zhou and Cox 2014).

In addition to employing covalently attached saturated fatty acids, proteins also can be permanently (absent proteolytic cleavage) anchored in the membrane via a transmembrane domain. Of the proteins investigated here, this was previously demonstrated for SRβ, a protein anchored in the ER membrane via its N-terminal transmembrane region (Keenan et al. 2001) that appears to be conserved in all SRβ sequences investigated (fig. 7G). An N-terminal transmembrane region was independently acquired by the Metazoa-specific Arl10 (see above) and several other ARF family members in various eukaryotes (fig. 7G; supplementary table 1, Supplementary Material online). In some cases, we could confirm conservation of such putative N-terminally anchored GTPases in a broader organism clade beyond the species primarily targeted by our analysis, as is the case of divergent putative Arf1 paralogs from *Bigelowiella natans* and other chlorarachniophytes (supplementary fig. 17A, Supplementary Material online) and from *Pavlova pinguis* and other haptophytes of the class Pavlovophyceae (supplementary fig. 17B, Supplementary Material online). Another mode of membrane attachment utilized by some ARF family members is accretion of specific membrane-binding domains. This is exemplified by unusual proteins from choanoflagellates and trypanosomatids that contain an N-terminal phosphoinositide-binding PH domain (Lemmon 2007) connected to the ARF family GTPase domain by a long linker region (fig. 7E; supplementary table 1, Supplementary Material online). Finally, the eustigmatophyte *Vischeria* sp. encodes a unique ARF family protein (VisArlX2 in supplementary table 1, Supplementary Material online) with a long N-terminal extension lacking any detectable conserved protein domain or functional motif and with a C-terminal tail ending with the amino acid sequence CSIM (fig. 7F), which is reminiscent of the so-called CaaX motif (or box) directing prenylation of the cysteine residue in diverse proteins (Fu and Casey 1999). A similar protein, including this motif, is encoded by additional eustigmatophytes (not shown), and two different prediction programs proposed the cysteine residue to be prenylated (see Material and Methods for details). C-terminal prenylation is a common modification ensuring membrane attachment of GTPases belonging to Rab, Ras and Rho families (Zhou and Cox 2014), but to our knowledge it has not been reported previously for an ARF family protein.

The well-studied mammalian members of the ARF family are subject to other post-translational modifications (e.g., see Phosphosite Plus; https://www.phosphosite.org/), though these either lack consensus motifs that prevent predicting their existence in other organisms or have no known functional consequences, or both. One exception to this is N-terminal acetylation of Arl8, which has been shown to be important for its association with lysosomal membranes (Hofmann and Munro 2006). Similarly, in *S. cerevisiae* the Arfrp1 protein (unfortunately named Arl3p only in this organism) is also acetylated and this is required for its association with Golgi membranes (Behnia et al. 2004). Future development of appropriate prediction tools, perhaps combined with dedicated biochemical investigations, will be instrumental in grasping the full breath and evolutionary conservation of PTMs in the ARF family.

In summary, the use of several different means of membrane attachment is consistent with ARF family proteins acting predominantly on a membrane surface, and the diversity of various membrane attachment mechanisms exhibited by this family is surprisingly extensive and reminiscent of what has been described for the distantly related GTPase Rheb (Záhonová et al. 2018). It is perhaps worth noting that eukaryotic organisms can vary widely in their lipid composition and the same is true of different organelles in an organism, making different means of membrane association likely important for this family of cell regulators that most often act on membrane surfaces and can even modify the lipid composition via direct activation of lipid kinases and lipases.

### Extensive diversity of multi-domain ARF family members

The existence of the PH domain-containing ARF family proteins or the aforementioned multi-domain TRIM23 protein (Vichi et al. 2005) counter the paradigm of ARF family members being limited to single (GTPase) domain proteins with only short N- and C-terminal extensions. In fact, our analyses challenge this dogma further. Although they represent a minority (75 out of >2,000 sequences in our dataset), multi-domain ARF family members represent a much greater number of different protein architectures involving combinations of the GTPase domain of the ARF family with other functional domains than thought previously (see column S in Supplementary table 1, Supplementary Material online).

The novel, presumably ancestral eukaryotic, Arl17 group characterized by combining an Arf-related domain with varying numbers of tandemly arrayed copies of a novel uncharacterized domain (fig. 2) was introduced above. Additional domain architectures are found in proteins that generally seem to be lineage-specific innovations restricted to particular taxa; some examples are provided in fig. 8. Similar to TRIM23, some include domains linked to ubiquitination, namely the BTB domain or the F-box domain (see Genschik et al. 2013), indicating recurrent recruitment of ARF family members into ubiquitin-dependent regulatory circuits. Ciliates exhibit a unique protein with an ARF family GTPase domain fused to a segment homologous to radial spoke protein 3 (RSP3), a component of radial spokes in the axoneme (see Wirschell et al. 2008). This predicts ciliary localization of this protein, and indeed, it is among the proteins detected in the ciliary proteome of *T. thermophila* (Smith et al. 2005). *Entamoeba histolytica* possesses a protein with a divergent C-terminal ARF family domain preceded by the VPS9 domain. The latter domain is known to act as a GEF of the endosomal Rab GTPase Rab5 (Ishida et al. 2016), so this protein may be part of a pathway with multiple sequentially acting GTPases similar to regulatory GTPase cascades known from mammalian or yeast cells (Jones et al. 1999; Mizuno-Yamasaki et al. 2012). Another unique domain combination occurs in one of the Arf paralogs in the haptophyte *Emiliania huxleyi*, which is fused to the C-terminus of a block including a domain of the 2OG-Fe(II) oxygenase superfamily. It is possible that the GTPase domain regulates the enzyme activity of the N-terminal part of the protein. The ARF family domain can combine also with other Ras superfamily GTPase domains, as demonstrated by a protein from *Malawimonas californiana* with an N-terminal Rab domain and a C-terminal Arf domain linked by a region containing detectable BTB and BACK domains (fig. 8A).

**Fig. 8.**
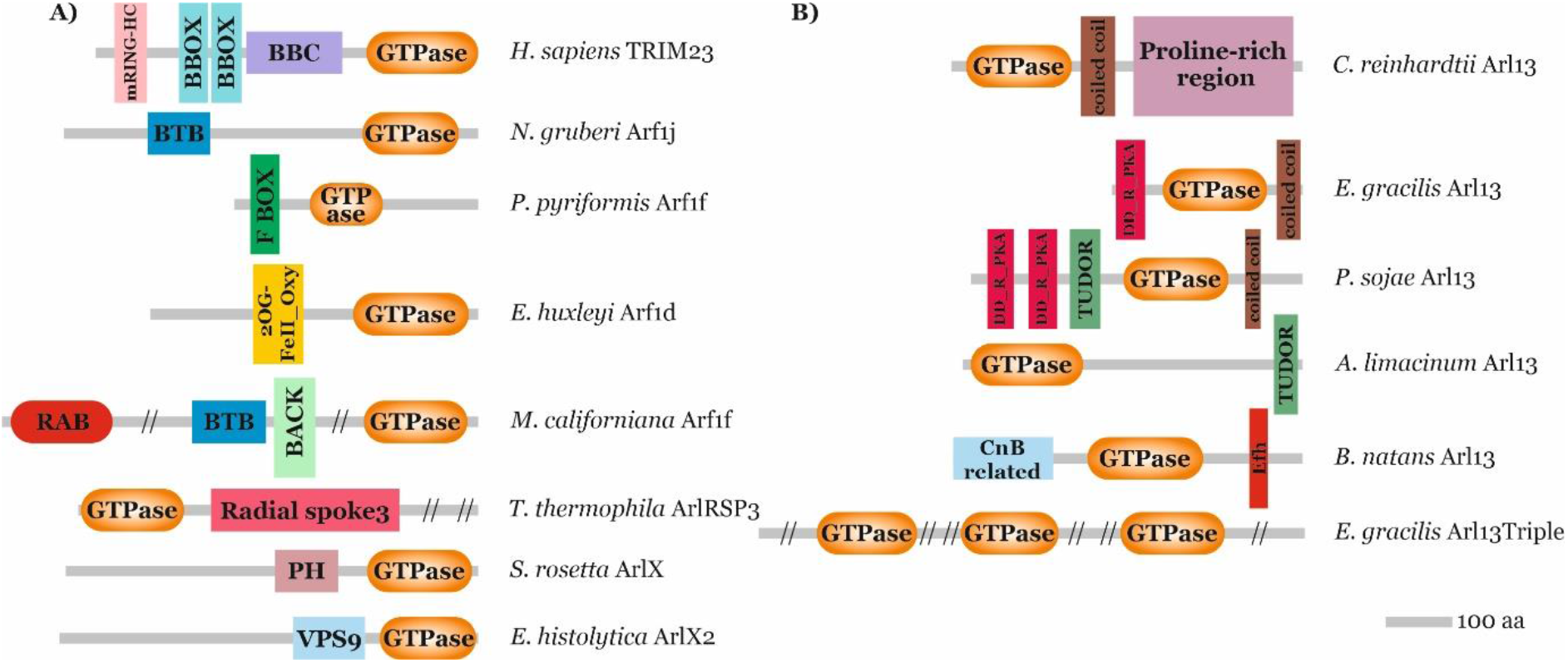
Multi-domain architectures of ARF family proteins. (**A**) Examples of lineage-specific ARF family proteins with extra domains accreted to the GTPase domain. Sequence IDs of the proteins listed are provided in supplementary table 1, Supplementary Material online. (**B**) Variation in the domain architecture of Arl13 proteins across the eukaryote diversity. The Arl13 from *Chlamydomonas reinhardtii* represents the most common and presumably ancestral state.

Tinkering with protein domains in ARF family proteins can be encountered in a different evolutionary context than the emergence of lineage-specific paralogs. In the case of Arl13, domains were acquired or lost without gene duplication, resulting in differences in domain architectures between orthologous Arl13 genes. Previously characterized orthologs from mammals and *Chlamydomonas reinhardtii* exhibit a poorly conserved C-terminal extension that includes a region forming a coiled-coil followed by a proline-rich region (Hori et al. 2008; Miertzschke et al. 2014; fig. 8B). Inspection of the large collection of Arl13 sequences amassed for this study revealed that this arrangement is distributed broadly across the eukaryote phylogeny and likely ancestral. However, some species (represented by eleven Arl13 genes out of 70 included in our dataset) depart in various way from this structure, e.g. by lacking the proline-rich region or the coiled-coil. Recently, Zhang et al. (2018) identified a non-canonical Arl13 gene from *Trypanosoma brucei* containing the DD_RI_PKA domain (Dimerization/Docking domain of the Regulatory subunit of protein kinase A (PKA)) that is essential for targeting of *T. brucei* Arl13 to the cilium. Our analysis revealed that the same protein architecture is present also in *Euglena gracilis*, suggesting it is a synapomorphic character for the whole Euglenozoa phylum (fig. 8B). Meanwhile, a subset of Stramenopiles (oomycetes and ochrophytes) independently acquired DD_RI_PKA domain as two tandemly arrayed copies (fig. 8B). DD_RI_PKA mediates interaction of PKA with A-kinase-anchoring proteins (AKAPs), which regulate PKA localization in the cell (Sarma et al. 2010). Given the ciliary function of Arl13 (see above), we speculate that the DD_RI_PKA domain in some Arl13 proteins interacts with a cilium-localized AKAP, such as the aforementioned RSP3 protein (Gaillard et al. 2001; Jivan et al. 2009). In contrast, mammalian Arl13b contains the simpler VxP motif in the large C-terminal domain that is required for ciliary localization (Higginbotham et al. 2012; Cevik et al. 2013; Gigante et al. 2020). DD_RI_PKA domains in *Phytophthora sojae* Arl13 are followed by the TUDOR domain, known for the ability to bind to the methylated lysine and/or arginine residues (Botuyan and Mer 2016). The TUDOR domain was independently accreted also to the C-terminus of the Arl13 from *Aurantiochytrium limacinum* (fig. 8B). Another notable variant is encountered in Arl13 from *B. natans* (fig. 8B) and other chlorarachniophytes (supplementary fig. 18, Supplementary Material online), which exhibit a novel form of the C-terminal extension including the Ca^2+^-binding EF-hand motif. Interestingly, the N-terminus of chlorarachniophyte Arl13 proteins appears to be related to calcineurin B, a Ca^2+^-binding regulatory subunit of the protein phosphatase calcineurin (Guerini 1997). It thus seems likely that Arl13 function is regulated by Ca^2+^ in chlorarachniophytes. Exceptional is an *E. gracilis* gene (co-occurring in this species with a typical Arl13 gene) that we named Arl13Triple, as it is composed of a tandem triplication of a divergent Arl13-reated GTPase domain (fig. 8B). The varying domain architecture of Arl13 in different eukaryotes points to a substantial degree of functional divergence of this key ciliary component.

### Insights into the early radiation of the ARF family

In the analysis of protein family evolution, resolution between the paralogs is a tremendously informative result as it allows the inference of cellular evolution of the associated organellar compartments. However, such resolution has been difficult to obtain for many families. The ScrollSaw methodology was a step forward in obtaining resolution for datasets with many paralogs and short sequence length; e.g., Rabs and TBC proteins (Rab GAPs; Elias et al. 2012; Gabernet-Castello et al. 2013). Here, our application of the ScrollSaw methodology also yielded a partially resolved backbone topology (fig. 1). We observed the robust sisterhood of Arl8 and Arl18 and of these both to Arl16. We also observed the sisterhood of Arl2 and Arl3 plus the moderately supported node uniting Arf1 with Arf6. Most notably, there was a strongly resolved node grouping together Arf1, Arf6, Arl1, Arl2, Arl3 and Arl5 and separating them from the remainder of the paralogs.

This resolution provides the basis for several key inferences about the ancestral role of the ARF family progenitor and some implications about the role of these proteins during eukaryogenesis. Taking only the most broadly conserved biochemical and cellular features of the various ARF family members, and assuming basic functional homology in orthologs to their roles in LECA (Klinger et al. 2016), what is likely ancestral is a GTPase that changes conformation to relocate from the cytosol to a membrane and which binds other proteins as effector(s). Given the widespread role of ARF family members, this may mean a role in membrane-traffic. However, with at least one resolved node separating the best-known family members Arf1 and Sar1, a simple scenario of a single primordial GTPase that nucleates a primordial vesicle coat-forming complex is ruled out. This suggests that the proto-coatomer hypothesis (Devos et al. 2004) may well need to be modified to take a more complicated scenario, including possible convergence, parallel evolution, and even merging of architectures into account (Dacks and Robinson 2017; Field and Rout 2019).

### Conclusions

Our comprehensive analysis of an extensive, well-curated dataset of ARF family proteins has provided evolutionary insights and raised questions to be addressed by future molecular cell biological exploration. The identification of 16 ancient ARF family paralogs both extends the inferred complexity of LECA and sets a framework of what components can be expected to be acting when delving into cellular function in diverse eukaryotes. By contrast the identification of expanded complements, including novel paralogs, e.g. the metazoan Arl19 and Arl20, provide specific new candidates for investigation in some of the best explored and heavily utilized cell biological model systems. The diversity of domain architecture challenges the paradigm of this family strictly as small GTPases and begs probing of new protein-protein interactions. Altogether, our work thus establishes a solid basis for future more detailed investigations into the biology of ARF family proteins at a eukaryote-wide scale.

## Materials and Methods

### Building and curation of the ARF family dataset

ARF family GTPases were searched in genome and/or transcriptome assemblies from 114 eukaryotic species selected such as to cover as many main eukaryotic lineages as possible (sequence identifiers, source databases, and further comments are provided in supplementary table 1, Supplementary Material online). The selection of taxa reflected the availability of relevant data as of 2018, when the sampling was frozen to obtain a final sequence dataset for all the subsequent analyses. As a result, several main eukaryote lineages, for which genome or transcriptome data became available more recently (e.g. the CRuMs supergroup, Telonemia, Rhodelphidia etc.), are not represented in our dataset. ARF family sequences were identified using BLAST and its variants (Altschul et al. 1997). Each organism-specific dataset was queried with reference members of the family and significant hits were evaluated by reverse BLAST searches against an in-house extensively curated taxonomically-rich database of GTPases. Query sequences being more similar to previously annotated members of the ARF family (including SRβ) were kept for further analysis. Existing protein sequence predictions were carefully evaluated and in a many cases revised by modifying the predicted exon-intron structure of the underlying gene model based on information from transcriptomic data or comparison to homologous sequences. To identify genes potentially missing from existing genome annotations, tblastn searches of nucleotide sequence data were carried out and gene models were created anew for previously missed genes. If possible, truncated sequences were completed using EST/TSA data or by iteratively recruiting raw genomic/transcriptomic sequencing reads. Revised, newly predicted or extended sequences are provided in supplementary table 1, Supplementary Material online. Putative pseudogenes (except for the human Arf2 pseudogene sequence, which can be reconstructed) as well as extremely divergent sequences with disrupted ARF family motif(s) were not included into the dataset and are not listed in supplementary table 1, Supplementary Material online.

Each gene was initially annotated by considering results of blastp searches against our comprehensive database of Ras superfamily proteins (iteratively updated by adding sequences newly annotated in the course of the study). In most cases, the blastp output enabled unambiguous assignment of the query sequence into one of the previously delineated ortholog groups or into novel orthogroups that emerged during the study. Sequences most similar to true Arfs, yet difficult to assign into the Arf1 or Arf6 groups or being visibly divergent were provisionally annotated as “ArfX”. Still more divergent ARF family members that did not show an apparently consistent affinity to a particular ARF family orthogroup when examined by BLAST searches were provisionally annotated as “ArlX”. The annotation of some of the ArfX and ArlX sequences was subsequently revised after the employment of the ScrollSaw protocol described below.

Sequence data from the glaucophyte *Gloeochaete wittrockiana* were included despite the fact that we noticed contamination of both available transcriptome assemblies (MMETSP0308 and MMETSP1089; https://www.imicrobe.us/#/projects/104) by sequences from an amoebozoan. The putative contaminants were identified by careful examination of individual sequences and excluded from the dataset. Another potential contaminant (the contig PCB_a545736;2 K: 25), showing a high similarity to Arl4 genes from primates, was noticed in the transcriptome assembly from the breviate *Pygsuia biforma* and removed from analyses.

### The ScrollSaw protocol and phylogenetic analyses

A master multiple alignment was built for the identified ARF family members, excluding short incomplete sequences and also all SRβ sequences, as this group is noticeably different from the core of the ARF family and many of its members tend to be rather divergent in their sequence. Altogether, the master alignment included 1931 sequences. It was built iteratively, starting with separate alignments for each group of sequences initially assigned to the same (potential) orthologous group using the on-line program MAFFT (version 7), with default parameters (Katoh and Standley 2013). All alignments were checked by eye and further edited manually using BioEdit (Hall 1999). The set of separate alignments was merged into one large alignment using the on-line Merge function of MAFFT. Divergent (ArlX) sequences were added to the alignment at the end. The final alignment was then manually trimmed to remove poorly conserved and unreliably aligned positions. After removing redundancies, the alignment comprised 1891 non-identical sequences and 148 aligned positions (all falling within the GTPase domain shared across the family).

The alignment was subjected to an analysis essentially following the previously published ScrollSaw protocol (Elias et al. 2012). The sequences were divided according to the source species into 13 taxonomic groups covering the known diversity of eukaryotes: Holozoa, Holomycota, Apusomonadida, Breviatea, Amoebozoa, Malawimonadida, Planomonadida, Discoba, Metamonada, Archaeplastida, Cryptista, Haptista, and SAR. The sizes of the groups differ substantially, since many evolutionarily important lineages were represented only by a small number of species with genomic or transcriptomic data available at the time when we initiated the study (in the case of Breviatea and Planomonadida by only a single species). The master alignment was then subsampled by keeping only sequences from each possible pair of the taxa listed above, corresponding to 78 combinations. For each of the 78 alignments, genetic distances between the sequences were inferred using the maximum likelihood (ML) method (with the WAG+Γ+I substitution model) implemented in Tree-Puzzle 5.3 (Schmidt et al. 2002). Each resulting distance matrix was analysed using a custom Python script to identify the so-called minimal-distance pairs. A minimal-distance pair consists of two sequences from the two different taxonomic group compared that have mutually minimal distances when distances to sequences from the other taxon are considered. Minimal-distance pairs from all 78 pairwise taxon comparisons were gathered and redundancies were removed, resulting in a set of 568 sequences. To further reduce the complexity of the dataset we then removed all sequences that formed only one minimal-distance pair in all 78 pairwise taxon comparisons combined. This step yielded the final full ScrollSaw dataset comprising 354 sequences. A reduced ScrollSaw variant was prepared by removing sequences from the majority of metamonads exhibiting generally divergent genes, including *Monocercomonoides exilis*, *Giardia intestinalis*, *Spironucleus* spp. and *Trichomonas vaginalis*.

The two variants of the final ScrollSaw dataset were used for inferring the ML phylogenetic trees using the program IQ-TREE (Nguyen et al. 2015). The substitution model (LG+I+G4) was selected by the program itself based on specific optimality criteria. Branch support was assessed by the SH-aLRT test (Guindon et al. 2010) and the ultrafast bootstrap approximation (Minh et al. 2013). The branch support of the reduced dataset was further examined by MrBayes 3.2 (Ronquist et al. 2011) using the CIPRES Science Gateway (Miller et al. 2010) with the following settings: prset aamodelpr=fixed(WAG); lset rates=gamma Ngammacat=4 mcmc ngen=1000000 printfreq=10000 samplefreq=1000 nchains=4 burnin=80. A number of additional alignments for specific dedicated analyses, derived by subsampling the master alignment or aligning the selected sequences *de novo* (using MAFFT with subsequent manual editing as described above) were used for ML phylogenetic inference using the same or similar approach. The alignments were in most cases trimmed according to the mask applied to the master alignment. Smaller phylogenetic analysis with only a subset of paralogous groups were trimmed either manually or by stand-alone version of trimAl (version 1.2rev57; option automated1; Capella-Gutierrez et al. 2009) in order to retrieve more positions for the ML phylogenetic analysis. The stand-alone version of IQ-TREE or the IQ-TREE web server (http://iqtree.cibiv.univie.ac.at/; Trifinopoulos et al. 2016) were used for the analyses. The substitution models were selected by the model selection program implemented in the IQ-TREE. Branch support was assessed by the SH-aLRT test and the ultrafast bootstrap approximation.

Tree topology testing was employed to test the hypothesis that SarB sequences form a monophyletic group sister to the Sar1 group. ML trees were inferred from the reduced SrollSaw alignment with a different topological constraints (specified in supplementary table 2, Supplementary Material online) using IQ-TREE and the same procedure as used for computing the unconstrained tree (shown in fig. 1). The unconstrained and constrained trees, together with a sample of 1,000 trees obtained as ultrafast bootstrap replicates in the unconstrained ML search on the alignment, were then compared in IQ-TREE (-au option) with the substitution models and its parameters optimized from the original alignment (-m TEST) and using 10,000 RELL replicates. The p-values of the alternative topologies obtained with the Kishino-Hasegawa (KH), Shimodaira-Hasegawa (SH), and approximately unbiased (AU) tests were considered.

### Annotation of sequences

The full and reduced ScrollSaw datasets were used as a basis for annotation of the rest of the sequences. The identity of individual sequences or their groups was tested by adding them to the reduced ScrollSaw dataset and inferring a ML tree with IQ-TREE. The scrutinized sequences were assigned to a particular ancestral eukaryotic paralog and annotated accordingly if they clustered together with reference representatives of the given paralog group and the relationship was supported by SH-aLRT and ultrafast bootstrap values of ≥80 and 95, respectively. Not all genes could be annotated by this approach, hence the HMMER package (stand alone version 3.0; hmmer.org) was employed as an alternative. The aligned full ScrollSaw dataset was divided into 14 separate alignments, each representing one ancestral paralog (Arf1, 6, Arl1, 2, 3, 5, 6, 8, 13, 16, 18, Arfrp1, Sar1, and SarB). A profile HMM was constructed for each alignment using hmmbuild and a database of profile HMMs was created using hmmpress. The unannotated sequences were then used as queries in hmmscan searches against the database and the “best 1 domain” score difference between the first and the second best hits was determined. If this difference was equal to or higher than 20, the sequence was annotated according to the best hit. Sequences annotated based on the phylogenetic analyses or hmmscan searches are marked by asterisk (*) in the column “Conclusively annotated” in the supplementary table 1, Supplementary Material online. Proteins representing SRβ and Arl17, which were not represented by reference sequences in the ScrollSaw dataset, were unequivocally identified owing to the distinct characters of these sequence groups, which makes them easy to recognise by BLAST-based similarity searches (SRβ) or by considering the presence of the novel conserved C-terminal domain (Arl17; see the main text). All SRβ and Arl17 proteins are therefore also considered as conclusively annotated. A single truncated sequence (Arl17b gene from *Chromera velia*) lacked the C-terminal extension with the characteristic C-terminal domain, but was assigned to the Arl17 group based on its close sequence similarity to undisputed Arl17 sequences. A combination of BLAST searches, ML phylogenetic analyses and comparison of exon-intron structure was used to obtain the most likely annotation of the sequences that could not be conclusively annotated by the aforementioned approaches. Several sequences were annotated as Arf1/6, as they showed affinity to the Arf1/6 clade, but it was impossible to decide whether they originated from ancestral Arf1 or Arf6 paralogs. Only 160 out of more than 2000 ARF family sequences analysed could not be annotated with any confidence, so they remained unassigned to any ancestral paralog (supplementary tables 1 and 3, Supplementary Material online).

### Taxon-specific ScrollSaw analyses and annotation of lineage-specific paralogs

To detect lineage-specific paralogs, we applied the ScrollSaw protocol separately to sets of sequences from the following main eukaryote taxa: Chloroplastida, Rhodophyta, Glaucophyta, Cryptista, Haptista, SAR, Discoba, Metamonada, Amoebozoa, Holomycota and Holozoa. Lineages represented by only one or two species (Apusomonadida, Breviatea, Planomonadida, and Malawimonadida) were not included. The ScrollSaw protocol and phylogenetic analyses of the resulted datasets were performed generally as described above for the whole dataset. For each main eukaryote taxon analysed, species representing it were assigned to predefined monophyletic subgroups specified in supplementary table 4, Supplementary Material online. Sequences from these species were extracted from the trimmed master alignment of the ARF family protein (except for sequences from Holozoa, which were aligned *de novo* using MAFFT and then trimmed according to the mask used for the whole dataset), the ScrollSaw protocol was applied to identify minimal-distance pairs, and ML phylogenetic trees were calculated on the filtered sequences. In contrast to the pan-eukaryotic ScrollSaw analysis, sequences that formed only one minimal-distance pair were not omitted (except for the analysis of the Holozoa dataset, where the criterion of the sequence belonging to at least two minimal-distance pairs was kept). The ML trees were inspected to identify robustly supported clades that would define conserved paralogs ancestral for the focal eukaryotic taxon but different from the previously defined ancestral eukaryote paralogs. In the case of Holozoa, paralogs specific for individual subgroups were considered, too. Further representatives of these paralogs (i.e., specific orthologs of the constituent sequences identified in the ScrollSaw trees) were then identified among the sequences that did not pass the ScrollSaw step by a combination of BLAST searches, phylogenetic analyses and (in case of Holozoa) HMMER-based comparisons. Candidates for ancestral taxon-specific paralogs were detected only in Chloroplastida, Rhodophyta, and Glaucophyta, as described in detail in the main text.

### Prediction of transmembrane regions and post-translation modifications

The presence of transmembrane (TM) regions in ARF family proteins was examined using the online TMHMM Server v. 2.0 (http://www.cbs.dtu.dk/services/TMHMM/). In the case of sequences with suspicious TM absence or presence (i.e., when the result was untypical for the respective ARF family subgroup), the on-line tool TMpred (https://www.ch.embnet.org/software/TMPRED_form.html) was additionally employed. The predictions are listed in supplementary table 1, Supplementary Material online. N-terminal myristoylation of sequences with the glycine residue at the second position was evaluated using the on-line ExPASy Myristoylator tool (http://web.expasy.org/myristoylator/; Bologna et al. 2004), NMT - The MYR Predictor (http://mendel.imp.ac.at/myristate/SUPLpredictor.htm; Maurer-Stroh et al. 2002), and the stand-alone version of GPS-Lipid (v1.0, http://lipid.biocuckoo.org/index.php; Xie et al. 2016). Only those proteins predicted as N-terminally myristoylated by at least two tools were considered as significant candidates. Setting of all tools was default except for NMT - The MYR Predictor where only N-terminal glycine residues were considered, and fungal sequences were predicted with the “Fungi specific” option. In case of GPS-Lipid, the threshold was set to “low”. Possible S-palmitoylation was predicted using SeqPalm (http://lishuyan.lzu.edu.cn/seqpalm/; Li et al. 2015), the stand-alone version of CKSAAP-Palm programme (http://doc.aporc.org/wiki/CKSAAP-Palm; Wang et al. 2009), PalmPred (http://proteininformatics.org/mkumar/palmpred/index.html; Kumari et al. 2014), stand-alone version of GPS-Lipid, and WAP-Palm (http://bioinfo.ncu.edu.cn/WAP-Palm.aspx; Shi et al. 2013). Only those sites predicted as S-palmitoylated by at least three tools were considered as significant candidates. Setting of all tools was default except for GPS-Lipid with the threshold set to “high”. Complete results from all tools are showed in supplementary table 5, Supplementary Material online, consensual results are included in supplementary table 1, Supplementary Material online. Possible prenylation was assessed only for VisArlX2 from the alga *Vischeria* sp., as it is the only protein from our dataset with a typical C-terminal prenylation motif. The online programs iPreny-PseAAC (http://app.aporc.org/iPreny-PseAAC/index.html; Xu et al. 2017) and GPS-Lipid were used with default settings; both tools predicted VisArlX2 as a prenylated protein.

### Other sequence analyses

Intron positions were investigated in four groups of ARF family genes (Sar1/SarB; Arl8/Arl18; Arfs and the GTPase domain of Arl17; Arfs and selected Arf-like in Holozoa) as a means to illuminate the origin and relationships of these genes. The positions of introns (including their phases) were mapped onto a multiple alignment of respective protein sequences using a custom Java script. The multiple sequence alignments were constructed *de novo* using MAFFT, inspected visually and adjusted manually whenever necessary (Sar1/SarB, Arl8/Arl18, Arf, and Arf-like in holozoans). For the analysis of Arf and Arl17 genes, the respective protein sequences were extracted from the master alignment. Sequences with no introns in the coding sequence or represented only by transcriptomic data were omitted. A manually curated dataset of gene exon-intron structures was used as the input for the intron positions mapping. For presentation purposes, regions corresponding to unconserved N- and C-termini of the sequences were trimmed and long sequence-specific insertions were collapsed. To highlight the pattern of protein sequence conservation, CHROMA (ver. 1.0 Goodstadt and Ponting 2001) was used for processing some of the multiple sequence alignments presented. Sequence logos of the Walker B motif were obtained using the on-line tool WebLogo 3 (http://weblogo.threeplusone.com/create.cgi; Crooks et al. 2004) from the multiple sequence alignment of the respective sequences after removing sequence-specific insertions present in a few sequences.

Conserved protein domains and other structural features in ARF family proteins were identified using searches of Pfam (http://pfam.xfam.org/; Finn et al. 2016), the Conserved Domains database (https://www.ncbi.nlm.nih.gov/Structure/cdd/wrpsb.cgi; Marchler-Bauer et al. 2017), and the SMART database (http://smart.embl-heidelberg.de/; Letunic and Bork 2018). Domain predictions provided by the three tools were compared and spurious results (low-significance with only a single tool) were ignored. The identity of unusual N-terminal extensions present in some Arl13 proteins were evaluated using HHpred (https://toolkit.tuebingen.mpg.de/; Söding et al. 2005). Multiple sequence alignments of the different forms on the N-terminal extensions conserved within different taxa were used as queries in the HHpred searches. In case of the N-terminal extension conserved in Arl13 proteins from Euglenozoa, the sampling was expanded beyond the focal set of taxa (including only three euglenozoans) by adding to the alignment several additional euglenozoan Arl13 sequences to improve the representativeness of the alignment. Similarly, additional chlorarachniophyte Arl13 sequences were identified and aligned with the sole representative in the focal dataset (that from *B. natans*), and additional stramenopile (oomycete and ochrophyte) Arl13 sequences with the same conserved N-terminal extension as the stramenopile sequences in the focal set were included to increase the sensitivity of the analysis. Some TRIM23 sequences were predicted by the standard tools to contain only one BBOX domain rather than the two common in most members of this group, but inspection of a multiple sequence alignment revealed high similarity of all sequences in the respective region, suggesting that all TRIM23 sequences likely conform to the same domain architecture with two BBOX domains.

## Supplementary Material

Supplementary data are available at Genome Biology and Evolution online.

## Acknowledgements

We thank Eunsoo Kim (American Museum of Natural History, New York) for an access to a genome assembly of *Goniomonas avonlea* prior to publication and Vladimír Hampl (Charles University in Prague) for his permission to use sequences from an unpublished genome assembly of *Paratrimastix pyriformis*. This work was supported by Czech Science Foundation grant 20-27648S; ERD Funds, project OPVVV CZ.02.1.01/0.0/0.0/16_019/0000759 (Centre for research of pathogenicity and virulence of parasites); and the infrastructure grant CZ.1.05/2.1.00/19.0388 („Přístroje IET“). Work in the Kahn lab is supported by a grant from the National Institutes of Health (NIH R35GM122568). Work in the Dacks Lab is supported by grants from the Natural Sciences and Engineering Research Council of Canada (RES0021028, RES0043758, RES0046091). JBD is the Canada Research Chair (Tier Ii) in Evolutionary Cell Biology.

## Data availability

ARF family gene sequences extracted from unpublished genome assemblies of *Gefionella okellyi, Planomonas micra*, and *Paratrimastix pyriformis*, and from our unpublished transcriptome assembly of *Vicheria* sp. CAUP Q 202, were deposited at GenBank with accession numbers #####-#####. A complete set of manually curated eukaryotic ARF family protein sequences is available in supplementary dataset 1, Supplementary Material online.

## Competing interests

No competing interests declared.

## Notes

### Competing Interest Statement

The authors have declared no competing interest.

## Literature Cited

Al-Bassam, J. (2017). Revisiting the tubulin cofactors and Arl2 in the regulation of soluble αβ-tubulin pools and their effect on microtubule dynamics. Molecular Biology of the Cell, 28(3), 359–363. https://doi.org/10.1091/mbc.E15-10-0694

Al-Bassam J. 2017. Revisiting the tubulin cofactors and Arl2 in the regulation of soluble αβ-tubulin pools and their effect on microtubule dynamics. Mol Biol Cell. 28:359–363. doi: 10.1091/mbc.E15-10-0694.

Altschul SF et al. 1997. Gapped BLAST and PSI-BLAST: a new generation of protein database search programs. Nucleic Acids Res. 25:3389–3402. doi: 10.1093/nar/25.17.3389.

Anantharaman V, Abhiman S, de Souza RF, Aravind L. 2011. Comparative genomics uncovers novel structural and functional features of the heterotrimeric GTPase signaling system. Gene. 475:63–78. doi: 10.1016/j.gene.2010.12.001.

Barlow LD, Nývltová E, Aguilar M, Tachezy J, Dacks JB. 2018. A sophisticated, differentiated Golgi in the ancestor of eukaryotes. BMC Biol. 16:27. doi: 10.1186/s12915-018-0492-9.

Behnia R, Panic B, Whyte JRC, Munro S. 2004. Targeting of the Arf-like GTPase Arl3p to the Golgi requires N-terminal acetylation and the membrane protein Sys1p. Nat Cell Biol. 6:405–413. doi: 10.1038/ncb1120.

Bologna G, Yvon C, Duvaud S, Veuthey A-L. 2004. N-Terminal myristoylation predictions by ensembles of neural networks. Proteomics. 4:1626–1632. doi: 10.1002/pmic.200300783.

Bosgraaf L, Van Haastert PJM. 2003. Roc, a Ras/GTPase domain in complex proteins. Biochim Biophys Acta. 1643:5–10. doi: 10.1016/j.bbamcr.2003.08.008.

Botuyan MV, Mer G. 2016. Chapter 8 - Tudor Domains as Methyl-Lysine and Methyl-Arginine Readers. In: Chromatin Signaling and Diseases. Binda, O & Fernandez-Zapico, ME, editors. Academic Press: Boston pp. 149–165. doi: 10.1016/B978-0-12-802389-1.00008-3.

Capella-Gutiérrez S, Silla-Martínez JM, Gabaldón T. 2009. trimAl: a tool for automated alignment trimming in large-scale phylogenetic analyses. Bioinformatics. 25:1972–1973. doi: 10.1093/bioinformatics/btp348.

Casanova JE. 2007. Regulation of Arf activation: the Sec7 family of guanine nucleotide exchange factors. Traffic. 8:1476–1485. doi: 10.1111/j.1600-0854.2007.00634.x.

Cevik S et al. 2013. Active transport and diffusion barriers restrict Joubert Syndrome-associated ARL13B/ARL-13 to an Inv-like ciliary membrane subdomain. PLoS Genet. 9:e1003977. doi: 10.1371/journal.pgen.1003977.

Cevik S et al. 2010. Joubert syndrome Arl13b functions at ciliary membranes and stabilizes protein transport in *Caenorhabditis elegans*. J Cell Biol. 188:953–969. doi: 10.1083/jcb.200908133.

Clark J et al. 1993. Selective amplification of additional members of the ADP-ribosylation factor (ARF) family: cloning of additional human and *Drosophila* ARF-like genes. Proc Natl Acad Sci U S A. 90:8952–8956. doi: 10.1073/pnas.90.19.8952.

Colicelli J. 2004. Human RAS superfamily proteins and related GTPases. Sci STKE. 2004:RE13. doi: 10.1126/stke.2502004re13.

Cotton M et al. 2007. Endogenous ARF6 interacts with Rac1 upon angiotensin II stimulation to regulate membrane ruffling and cell migration. Mol Biol Cell. 18:501–511. doi: 10.1091/mbc.e06-06-0567.

Crooks GE, Hon G, Chandonia J-M, Brenner SE. 2004. WebLogo: A Sequence Logo Generator. Genome Res. 14:1188–1190. doi: 10.1101/gr.849004.

Dacks JB et al. 2016. The changing view of eukaryogenesis - fossils, cells, lineages and how they all come together. J. Cell. Sci. 129:3695–3703. doi: 10.1242/jcs.178566.

Dacks JB, Robinson MS. 2017. Outerwear through the ages: evolutionary cell biology of vesicle coats. Curr Opin Cell Biol. 47:108–116. doi: 10.1016/j.ceb.2017.04.001.

van Dam TJP, Bos JL, Snel B. 2011. Evolution of the Ras-like small GTPases and their regulators. Small GTPases. 2:4–16. doi: 10.4161/sgtp.2.1.15113.

Derelle R et al. 2015. Bacterial proteins pinpoint a single eukaryotic root. Proc Natl Acad Sci U S A. 112:E693–E699. doi: 10.1073/pnas.1420657112.

Devos D et al. 2004. Components of coated vesicles and nuclear pore complexes share a common molecular architecture. PLoS Biol. 2:e380. doi: 10.1371/journal.pbio.0020380.

Dian C et al. 2020. High-resolution snapshots of human N-myristoyltransferase in action illuminate a mechanism promoting N-terminal Lys and Gly myristoylation. Nat Commun. 11:1132. doi: 10.1038/s41467-020-14847-3.

Diekmann Y et al. 2011. Thousands of Rab GTPases for the Cell Biologist. PLOS Comput Biol. 7:e1002217. doi: 10.1371/journal.pcbi.1002217.

Donaldson JG, Jackson CL. 2011. Arf Family G Proteins and their regulators: roles in membrane transport, development and disease. Nat Rev Mol Cell Biol. 12:362–375. doi: 10.1038/nrm3117.

D’Souza-Schorey C, Chavrier P. 2006. ARF proteins: roles in membrane traffic and beyond. Nat Rev Mol Cell Biol. 7:347–358. doi: 10.1038/nrm1910.

D’Souza-Schorey C, Stahl PD. 1995. Myristoylation is required for the intracellular localization and endocytic function of ARF6. Exp Cell Res. 221:153–159. doi: 10.1006/excr.1995.1362.

Duronio RJ, Rudnick DA, Adams SP, Towler DA, Gordon JI. 1991. Analyzing the substrate specificity of *Saccharomyces cerevisiae* myristoyl-CoA:protein N-myristoyltransferase by co-expressing it with mammalian G protein α subunits in *Escherichia coli*. J Biol Chem. 266:10498–10504. doi: 10.1016/S0021-9258(18)99252-5.

Elias M, Archibald JM. 2009. The RJL family of small GTPases is an ancient eukaryotic invention probably functionally associated with the flagellar apparatus. Gene. 442:63–72. doi: 10.1016/j.gene.2009.04.011.

Elias M, Brighouse A, Gabernet-Castello C, Field MC, Dacks JB. 2012. Sculpting the endomembrane system in deep time: high resolution phylogenetics of Rab GTPases. J Cell Sci. 125:2500–2508. doi: 10.1242/jcs.101378.

Eliáš M, Klimeš V, Derelle R, Petrželková R, Tachezy J. 2016. A paneukaryotic genomic analysis of the small GTPase RABL2 underscores the significance of recurrent gene loss in eukaryote evolution. Biol Direct. 11:5. doi: 10.1186/s13062-016-0107-8.

Eme L, Spang A, Lombard J, Stairs CW, Ettema TJG. 2017. Archaea and the origin of eukaryotes. Nat Rev Microbiol. 15:711–723. doi: 10.1038/nrmicro.2017.133.

Fansa EK, Wittinghofer A. 2016. Sorting of lipidated cargo by the Arl2/Arl3 system. Small GTPases. 7:222–230. doi: 10.1080/21541248.2016.1224454.

Field MC, Rout MP. 2019. Pore timing: the evolutionary origins of the nucleus and nuclear pore complex. F1000Res. 8. doi: 10.12688/f1000research.16402.1.

Fielding AB et al. 2005. Rab11-FIP3 and FIP4 interact with Arf6 and the exocyst to control membrane traffic in cytokinesis. EMBO J. 24:3389–3399. doi: 10.1038/sj.emboj.7600803.

Finn RD et al. 2016. The Pfam protein families database: towards a more sustainable future. Nucleic Acids Res. 44:D279–285. doi: 10.1093/nar/gkv1344.

Fisher S, Kuna D, Caspary T, Kahn RA, Sztul E. 2020. ARF family GTPases with links to cilia. Am J Physiol Cell Physiol. 319:C404–C418. doi: 10.1152/ajpcell.00188.2020.

Flot J-F et al. 2013. Genomic evidence for ameiotic evolution in the bdelloid rotifer *Adineta vaga*. Nature. 500:453–457. doi: 10.1038/nature12326.

Francis JW, Goswami D, et al. 2017. Nucleotide Binding to ARL2 in the TBCD·ARL2·β-Tubulin Complex Drives Conformational Changes in β-Tubulin. J Mol Biol. 429:3696–3716. doi: 10.1016/j.jmb.2017.09.016.

Francis JW, Newman LE, Cunningham LA, Kahn RA. 2017. A Trimer Consisting of the Tubulin-specific Chaperone D (TBCD), Regulatory GTPase ARL2, and β-Tubulin Is Required for Maintaining the Microtubule Network. J Biol Chem. 292:4336–4349. doi: 10.1074/jbc.M116.770909.

Francis JW, Turn RE, Newman LE, Schiavon C, Kahn RA. 2016. Higher order signaling: ARL2 as regulator of both mitochondrial fusion and microtubule dynamics allows integration of 2 essential cell functions. Small GTPases. 7:188–196. doi: 10.1080/21541248.2016.1211069.

Fu HW, Casey PJ. 1999. Enzymology and biology of CaaX protein prenylation. Recent Prog Horm Res. 54:315–342; discussion 342-343.

Funakoshi Y, Hasegawa H, Kanaho Y. 2011. Regulation of PIP5K activity by Arf6 and its physiological significance. J Cell Physiol. 226:888–895. doi: 10.1002/jcp.22482.

Gabernet-Castello C, O’Reilly AJ, Dacks JB, Field MC. 2013. Evolution of Tre-2/Bub2/Cdc16 (TBC) Rab GTPase-activating proteins. Mol Biol Cell. 24:1574–1583. doi: 10.1091/mbc.E12-07-0557.

Gaillard AR, Diener DR, Rosenbaum JL, Sale WS. 2001. Flagellar radial spoke protein 3 is an A-kinase anchoring protein (AKAP). J Cell Biol. 153:443–448. doi: 10.1083/jcb.153.2.443.

Genschik P, Sumara I, Lechner E. 2013. The emerging family of CULLIN3-RING ubiquitin ligases (CRL3s): cellular functions and disease implications. EMBO J. 32:2307–2320. doi: 10.1038/emboj.2013.173.

Gigante ED, Taylor MR, Ivanova AA, Kahn RA, Caspary T. 2020. ARL13B regulates Sonic hedgehog signaling from outside primary cilia. Elife. 9. doi: 10.7554/eLife.50434.

Gillingham AK, Munro S. 2007. The small G proteins of the Arf family and their regulators. Annu. Rev. Cell Dev. Biol. 23:579–611. doi: 10.1146/annurev.cellbio.23.090506.123209.

Goodstadt L, Ponting CP. 2001. CHROMA: consensus-based colouring of multiple alignments for publication. Bioinformatics. 17:845–846. doi: 10.1093/bioinformatics/17.9.845.

Gotthardt K et al. 2015. A G-protein activation cascade from Arl13B to Arl3 and implications for ciliary targeting of lipidated proteins. Elife. 4. doi: 10.7554/eLife.11859.

Guerini D. 1997. Calcineurin: not just a simple protein phosphatase. Biochem Biophys Res Commun. 235:271–275. doi: 10.1006/bbrc.1997.6802.

Guindon S et al. 2010. New Algorithms and Methods to Estimate Maximum-Likelihood Phylogenies: Assessing the Performance of PhyML 3.0. Syst Biol. 59:307–321. doi: 10.1093/sysbio/syq010.

Hall TA. 1999. BioEdit: a user-friendly biological sequence alignment editor and analysis program for Windows 95/98/NT. Nucl. Acids. Symp. Ser. 41:95–98.

Heazlewood JL, Verboom RE, Tonti-Filippini J, Small I, Millar AH. 2007. SUBA: the *Arabidopsis* Subcellular Database. Nucleic Acids Res. 35:D213–D218. doi: 10.1093/nar/gkl863.

Higginbotham H et al. 2012. Arl13b in primary cilia regulates the migration and placement of interneurons in the developing cerebral cortex. Dev Cell. 23:925–938. doi: 10.1016/j.devcel.2012.09.019.

Hofmann I, Munro S. 2006. An N-terminally acetylated Arf-like GTPase is localised to lysosomes and affects their motility. J Cell Sci. 119:1494–1503. doi: 10.1242/jcs.02958.

Hofmann I, Thompson A, Sanderson CM, Munro S. 2007. The Arl4 family of small G proteins can recruit the cytohesin Arf6 exchange factors to the plasma membrane. Curr Biol. 17:711–716. doi: 10.1016/j.cub.2007.03.007.

Hori Y, Kobayashi T, Kikko Y, Kontani K, Katada T. 2008. Domain architecture of the atypical Arf-family GTPase Arl13b involved in cilia formation. Biochem Biophys Res Commun. 373:119–124. doi: 10.1016/j.bbrc.2008.06.001.

Houghton FJ et al. 2012. Arl5b is a Golgi-localised small G protein involved in the regulation of retrograde transport. Exp Cell Res. 318:464–477. doi: 10.1016/j.yexcr.2011.12.023.

Insall R, Gaudet P, Weeks G. 2005. The small GTPase superfamily. In: Loomis WF, Kuspa A, editors. Dictyostelium Genomics. Norfolk: Horizon Bioscience. p. 173–210.

Ishida M, E Oguchi M, Fukuda M. 2016. Multiple Types of Guanine Nucleotide Exchange Factors (GEFs) for Rab Small GTPases. Cell Struct Funct. 41:61–79. doi: 10.1247/csf.16008.

Ivanova AA et al. 2017. Biochemical characterization of purified mammalian ARL13B protein indicates that it is an atypical GTPase and ARL3 guanine nucleotide exchange factor (GEF). J Biol Chem. 292:11091–11108. doi: 10.1074/jbc.M117.784025.

Jackson CL. 2014. Arf Proteins and Their Regulators: At the Interface Between Membrane Lipids and the Protein Trafficking Machinery. In: Ras Superfamily Small G Proteins: Biology and Mechanisms 2: Transport. Wittinghofer, A, editor. Springer International Publishing: Cham pp. 151–180. doi: 10.1007/978-3-319-07761-1_8.

Jackson CL, Bouvet S. 2014. Arfs at a glance. J Cell Sci. 127:4103–4109. doi: 10.1242/jcs.144899.

Jivan A, Earnest S, Juang Y-C, Cobb MH. 2009. Radial spoke protein 3 is a mammalian protein kinase A-anchoring protein that binds ERK1/2. J Biol Chem. 284:29437–29445. doi: 10.1074/jbc.M109.048181.

Jones S et al. 1999. Genetic interactions in yeast between Ypt GTPases and Arf guanine nucleotide exchangers. Genetics. 152:1543–1556.

Kaessmann H, Vinckenbosch N, Long M. 2009. RNA-based gene duplication: mechanistic and evolutionary insights. Nat Rev Genet. 10:19–31. doi: 10.1038/nrg2487.

Kahn RA et al. 2008. Consensus nomenclature for the human ArfGAP domain-containing proteins. J Cell Biol. 182:1039–1044. doi: 10.1083/jcb.200806041.

Kahn RA et al. 2006. Nomenclature for the human Arf family of GTP-binding proteins: ARF, ARL, and SAR proteins. J Cell Biol. 172:645–650. doi: 10.1083/jcb.200512057.

Kahn RA. 2009. Toward a model for Arf GTPases as regulators of traffic at the Golgi. FEBS Lett. 583:3872–3879. doi: 10.1016/j.febslet.2009.10.066.

Kahn RA, Goddard C, Newkirk M. 1988. Chemical and immunological characterization of the 21-kDa ADP-ribosylation factor of adenylate cyclase. J. Biol. Chem. 263:8282–8287.

Katoh K, Standley DM. 2013. MAFFT Multiple Sequence Alignment Software Version 7: Improvements in Performance and Usability. Mol Biol Evol. 30:772–780. doi: 10.1093/molbev/mst010.

Keenan RJ, Freymann DM, Stroud RM, Walter P. 2001. The signal recognition particle. Annu. Rev. Biochem. 70:755–775. doi: 10.1146/annurev.biochem.70.1.755.

Khatter D, Sindhwani A, Sharma M. 2015. Arf-like GTPase Arl8: Moving from the periphery to the center of lysosomal biology. Cell Logist. 5. doi: 10.1080/21592799.2015.1086501.

Klinger CM, et al. 2016. Resolving the homology-function relationship through comparative genomics of membrane-trafficking machinery and parasite cell biology. Mol Biochem Parasitol. 209:88–103. doi: 10.1016/j.molbiopara.2016.07.003.

Klinger CM, Spang A, Dacks JB, Ettema TJ. 2016. Tracing the Archaeal Origins of Eukaryotic Membrane-Trafficking System Building Blocks. Mol Biol Evol. 33:1528–1541. doi: 10.1093/molbev/msw034.

Klöpper TH, Kienle N, Fasshauer D, Munro S. 2012. Untangling the evolution of Rab G proteins: implications of a comprehensive genomic analysis. BMC Biol. 10:71. doi: 10.1186/1741-7007-10-71.

Kumari B, Kumar R, Kumar M. 2014. PalmPred: an SVM based palmitoylation prediction method using sequence profile information. PLoS ONE. 9:e89246. doi: 10.1371/journal.pone.0089246.

Lee FJ et al. 1997. Characterization of an ADP-ribosylation factor-like 1 protein in *Saccharomyces cerevisiae*. J Biol Chem. 272:30998–31005. doi: 10.1074/jbc.272.49.30998.

Leipe DD, Wolf YI, Koonin EV, Aravind L. 2002. Classification and evolution of P-loop GTPases and related ATPases. J. Mol. Biol. 317:41–72. doi: 10.1006/jmbi.2001.5378.

Lemmon MA. 2007. Pleckstrin homology (PH) domains and phosphoinositides. Biochem Soc Symp. 81–93. doi: 10.1042/BSS0740081.

Letunic I, Bork P. 2018. 20 years of the SMART protein domain annotation resource. Nucleic Acids Res. 46:D493–D496. doi: 10.1093/nar/gkx922.

Li C-C et al. 2007. ARL4D recruits cytohesin-2/ARNO to modulate actin remodeling. Mol Biol Cell. 18:4420–4437. doi: 10.1091/mbc.e07-02-0149.

Li S et al. 2015. In Silico Identification of Protein S-Palmitoylation Sites and Their Involvement in Human Inherited Disease. J Chem Inf Model. 55:2015–2025. doi: 10.1021/acs.jcim.5b00276.

Li Y et al. 2004. Functional genomic analysis of the ADP-ribosylation factor family of GTPases: phylogeny among diverse eukaryotes and function in *C. elegans*. FASEB J. 18:1834–1850. doi: 10.1096/fj.04-2273com.

Lin C-Y, Li C-C, Huang P-H, Lee F-JS. 2002. A developmentally regulated ARF-like 5 protein (ARL5), localized to nuclei and nucleoli, interacts with heterochromatin protein 1. J Cell Sci. 115:4433–4445. doi: 10.1242/jcs.00123.

Liu Y, Kahn RA, Prestegard JH. 2010. Dynamic structure of membrane-anchored Arf*GTP. Nat Struct Mol Biol. 17:876–881. doi: 10.1038/nsmb.1853.

Liu Y, Kahn RA, Prestegard JH. 2009. Structure and Membrane Interaction of Myristoylated ARF1. Structure. 17:79–87. doi: 10.1016/j.str.2008.10.020.

Makiuchi T, Nozaki T. 2014. Highly divergent mitochondrion-related organelles in anaerobic parasitic protozoa. Biochimie. 100:3–17. doi: 10.1016/j.biochi.2013.11.018.

Manolea F et al. 2010. Arf3 Is Activated Uniquely at the trans-Golgi Network by Brefeldin A-inhibited Guanine Nucleotide Exchange Factors. Mol Biol Cell. 21:1836–1849. doi: 10.1091/mbc.E10-01-0016.

Marchler-Bauer A et al. 2015. CDD: NCBI’s conserved domain database. Nucleic Acids Res. 43:D222-226. doi: 10.1093/nar/gku1221.

Maurer-Stroh S, Eisenhaber B, Eisenhaber F. 2002. N-terminal N-myristoylation of proteins: prediction of substrate proteins from amino acid sequence. J. Mol. Biol. 317:541–557. doi: 10.1006/jmbi.2002.5426.

Melville DB, Studer S, Schekman R. 2020. Small sequence variations between two mammalian paralogs of the small GTPase SAR1 underlie functional differences in coat protein complex II assembly. J Biol Chem. 295:8401–8412. doi: 10.1074/jbc.RA120.012964.

Meza I, Talamás-Rohana P, Vargas MA. 2006. The Cytoskeleton of *Entamoeba histolytica*: Structure, Function, and Regulation by Signaling Pathways. Arch Med Res. 37:234–243. doi: 10.1016/j.arcmed.2005.09.008.

Miertzschke M, Koerner C, Spoerner M, Wittinghofer A. 2014. Structural insights into the small G-protein Arl13B and implications for Joubert syndrome. Biochem J. 457:301–311. doi: 10.1042/BJ20131097.

Miller EA, Barlowe C. 2010. Regulation of coat assembly--sorting things out at the ER. Curr Opin Cell Biol. 22:447–453. doi: 10.1016/j.ceb.2010.04.003.

Miller MA, Pfeiffer W, Schwartz T. 2010. Creating the CIPRES Science Gateway for inference of large phylogenetic trees. In: 2010 Gateway Computing Environments Workshop (GCE). pp. 1–8. doi: 10.1109/GCE.2010.5676129.

Minh BQ, Nguyen MAT, von Haeseler A. 2013. Ultrafast approximation for phylogenetic bootstrap. Mol. Biol. Evol. 30:1188–1195. doi: 10.1093/molbev/mst024.

Mizuno-Yamasaki E, Rivera-Molina F, Novick P. 2012. GTPase networks in membrane traffic. Annu Rev Biochem. 81:637–659. doi: 10.1146/annurev-biochem-052810-093700.

More K, Klinger CM, Barlow LD, Dacks JB. 2020. Evolution and Natural History of Membrane Trafficking in Eukaryotes. Curr Biol. 30:R553–R564. doi: 10.1016/j.cub.2020.03.068.

Mourão A, Nager AR, Nachury MV, Lorentzen E. 2014. Structural basis for membrane targeting of the BBSome by ARL6. Nat Struct Mol Biol. 21:1035–1041. doi: 10.1038/nsmb.2920.

Nabais C, Peneda C, Bettencourt-Dias M. 2020. Evolution of centriole assembly. Curr Biol. 30:R494–R502. doi: 10.1016/j.cub.2020.02.036.

Neuwald AF. 2007. Gα-Gβγ dissociation may be due to retraction of a buried lysine and disruption of an aromatic cluster by a GTP-sensing Arg Trp pair. Protein Sci. 16:2570–2577. doi: 10.1110/ps.073098107.

Newman LE, Schiavon CR, Turn RE, Kahn RA. 2017. The ARL2 GTPase regulates mitochondrial fusion from the intermembrane space. Cell Logist. 7:e1340104. doi: 10.1080/21592799.2017.1340104.

Nguyen L-T, Schmidt HA, von Haeseler A, Minh BQ. 2015. IQ-TREE: a fast and effective stochastic algorithm for estimating maximum-likelihood phylogenies. Mol. Biol. Evol. 32:268–274. doi: 10.1093/molbev/msu300.

Panic B, Whyte JRC, Munro S. 2003. The ARF-like GTPases Arl1p and Arl3p Act in a Pathway that Interacts with Vesicle-Tethering Factors at the Golgi Apparatus. Curr Biol. 13:405–410. doi: 10.1016/S0960-9822(03)00091-5.

Pasqualato S, Renault L, Cherfils J. 2002. Arf, Arl, Arp and Sar proteins: a family of GTP-binding proteins with a structural device for ‘front–back’ communication. EMBO Rep. 3:1035–1041. doi: 10.1093/embo-reports/kvf221.

Patel M, Chiang T-C, Tran V, Lee F-JS, Côté J-F. 2011. The Arf family GTPase Arl4A complexes with ELMO proteins to promote actin cytoskeleton remodeling and reveals a versatile Ras-binding domain in the ELMO proteins family. J Biol Chem. 286:38969–38979. doi: 10.1074/jbc.M111.274191.

Pereira-Leal JB. 2008. The Ypt/Rab family and the evolution of trafficking in fungi. Traffic. 9:27–38. doi: 10.1111/j.1600-0854.2007.00667.x.

Petrželková R, Eliáš M. 2014. Contrasting patterns in the evolution of the Rab GTPase family in Archaeplastida. Acta Soc Bot Polon. 83:303–315. doi: 10.5586/asbp.2014.052.

Pipaliya SV, Schlacht A, Klinger CM, Kahn RA, Dacks J. 2019. Ancient complement and lineage-specific evolution of the Sec7 ARF GEF proteins in eukaryotes. Mol Biol Cell. 30:1846–1863. doi: 10.1091/mbc.E19-01-0073.

Resh MD. 1999. Fatty acylation of proteins: new insights into membrane targeting of myristoylated and palmitoylated proteins. Biochim Biophys Acta. 1451:1–16. doi: 10.1016/s0167-4889(99)00075-0.

Rojas AM, Fuentes G, Rausell A, Valencia A. 2012. The Ras protein superfamily: Evolutionary tree and role of conserved amino acids. J Cell Biol. 196:189–201. doi: 10.1083/jcb.201103008.

Ronquist F et al. 2012. MrBayes 3.2: efficient Bayesian phylogenetic inference and model choice across a large model space. Syst Biol. 61:539–542. doi: 10.1093/sysbio/sys029.

Rosa-Ferreira C, Christis C, Torres IL, Munro S. 2015. The small G protein Arl5 contributes to endosome-to-Golgi traffic by aiding the recruitment of the GARP complex to the Golgi. Biol Open. 4:474–481. doi: 10.1242/bio.201410975.

Roy D, Lohia A. 2004. Sequence divergence of *Entamoeba histolytica* tubulin is responsible for its altered tertiary structure. Biochem Biophys Res Commun. 319:1010–1016. doi: 10.1016/j.bbrc.2004.05.079.

Roy K et al. 2017. Palmitoylation of the ciliary GTPase ARL13b is necessary for its stability and its role in cilia formation. J. Biol. Chem. 292:17703–17717. doi: 10.1074/jbc.M117.792937.

Sarma GN et al. 2010. Structure of D-AKAP2:PKA RI complex: Insights into AKAP specificity and selectivity. Structure. 18:155–166. doi: 10.1016/j.str.2009.12.012.

Sato K, Nakano A. 2007. Mechanisms of COPII vesicle formation and protein sorting. FEBS Lett. 581:2076–2082. doi: 10.1016/j.febslet.2007.01.091.

Schlacht A, Dacks JB. 2015. Unexpected ancient paralogs and an evolutionary model for the COPII coat complex. Genome Biol Evol. 7:1098–1109. doi: 10.1093/gbe/evv045.

Schmidt HA, Strimmer K, Vingron M, von Haeseler A. 2002. TREE-PUZZLE: maximum likelihood phylogenetic analysis using quartets and parallel computing. Bioinformatics. 18:502–504. doi: 10.1093/bioinformatics/18.3.502.

Schwartz T, Blobel G. 2003. Structural Basis for the Function of the β Subunit of the Eukaryotic Signal Recognition Particle Receptor. Cell. 112:793–803. doi: 10.1016/S0092-8674(03)00161-2.

Schweitzer JK, Sedgwick AE, D’Souza-Schorey C. 2011. ARF6-mediated endocytic recycling impacts cell movement, cell division and lipid homeostasis. Semin Cell Dev Biol. 22:39–47. doi: 10.1016/j.semcdb.2010.09.002.

Seidel RD, Amor JC, Kahn RA, Prestegard JH. 2004. Conformational changes in human Arf1 on nucleotide exchange and deletion of membrane-binding elements. J Biol Chem. 279:48307–48318. doi: 10.1074/jbc.M402109200.

Setty SRG, Shin ME, Yoshino A, Marks MS, Burd CG. 2003. Golgi Recruitment of GRIP Domain Proteins by Arf-like GTPase 1 Is Regulated by Arf-like GTPase 3. Curr Biol. 13:401–404. doi: 10.1016/S0960-9822(03)00089-7.

Setty SRG, Strochlic TI, Tong AHY, Boone C, Burd CG. 2004. Golgi targeting of ARF-like GTPase Arl3p requires its Nα-acetylation and the integral membrane protein Sys1p. Nat Cell Biol. 6:414–419. doi: 10.1038/ncb1121.

Sharer JD, Shern JF, Van Valkenburgh H, Wallace DC, Kahn RA. 2002. ARL2 and BART Enter Mitochondria and Bind the Adenine Nucleotide Transporter. Mol Biol Cell. 13:71–83. doi: 10.1091/mbc.01-05-0245.

Shi S-P et al. 2013. The prediction of palmitoylation site locations using a multiple feature extraction method. J. Mol. Graph. Model. 40:125–130. doi: 10.1016/j.jmgm.2012.12.006.

Smith JC, Northey JGB, Garg J, Pearlman RE, Siu KWM. 2005. Robust Method for Proteome Analysis by MS/MS Using an Entire Translated Genome: Demonstration on the Ciliome of *Tetrahymena thermophila*. J. Proteome Res. 4:909–919. doi: 10.1021/pr050013h.

Söding J, Biegert A, Lupas AN. 2005. The HHpred interactive server for protein homology detection and structure prediction. Nucleic Acids Res. 33:W244–W248. doi: 10.1093/nar/gki408.

Stephen LA, Ismail S. 2016. Shuttling and sorting lipid-modified cargo into the cilia. Biochem Soc Trans. 44:1273–1280. doi: 10.1042/BST20160122.

Sztul E et al. 2019. ARF GTPases and their GEFs and GAPs: concepts and challenges. Mol Biol Cell. 30:1249–1271. doi: 10.1091/mbc.E18-12-0820.

Tamkun JW et al. 1991. The arflike gene encodes an essential GTP-binding protein in *Drosophila*. Proc Natl Acad Sci U S A. 88:3120–3124. doi: 10.1073/pnas.88.8.3120.

Tria FDK, Landan G, Dagan T. 2017. Phylogenetic rooting using minimal ancestor deviation. Nat Ecol Evol. 1:193. doi: 10.1038/s41559-017-0193.

Trifinopoulos J, Nguyen L-T, von Haeseler A, Minh BQ. 2016. W-IQ-TREE: a fast online phylogenetic tool for maximum likelihood analysis. Nucleic Acids Res. 44:W232–W235. doi: 10.1093/nar/gkw256.

Turn RE, East MP, Prekeris R, Kahn RA. 2020. The ARF GAP ELMOD2 acts with different GTPases to regulate centrosomal microtubule nucleation and cytokinesis. Mol Biol Cell. 31:2070–2091. doi: 10.1091/mbc.E20-01-0012.

Van Valkenburgh H, Shern JF, Sharer JD, Zhu X, Kahn RA. 2001. ADP-ribosylation factors (ARFs) and ARF-like 1 (ARL1) have both specific and shared effectors: characterizing ARL1-binding proteins. J Biol Chem. 276:22826–22837. doi: 10.1074/jbc.M102359200.

Vernoud V, Horton AC, Yang Z, Nielsen E. 2003. Analysis of the small GTPase gene superfamily of *Arabidopsis*. Plant Physiol. 131:1191–1208. doi: 10.1104/pp.013052.

Vetter IR. 2014. The Structure of the G Domain of the Ras Superfamily. In: Ras Superfamily Small G Proteins: Biology and Mechanisms 1: General Features, Signaling. Wittinghofer, A, editor. Springer: Vienna pp. 25–50. doi: 10.1007/978-3-7091-1806-1_2.

Vichi A, Moss J, Vaughan M. 2005. ADP-ribosylation factor domain protein 1 (ARD1), a multifunctional protein with ubiquitin E3 ligase, GAP, and ARF domains. Meth. Enzymol. 404:195–206. doi: 10.1016/S0076-6879(05)04019-X.

Vlahou G, Eliáš M, von Kleist-Retzow J-C, Wiesner RJ, Rivero F. 2011. The Ras related GTPase Miro is not required for mitochondrial transport in *Dictyostelium discoideum*. Eur J Cell Biol. 90:342–355. doi: 10.1016/j.ejcb.2010.10.012.

Vosseberg J et al. 2021. Timing the origin of eukaryotic cellular complexity with ancient duplications. Nat Ecol Evol. 5:92–100. doi: 10.1038/s41559-020-01320-z.

Wang X-B, Wu L-Y, Wang Y-C, Deng N-Y. 2009. Prediction of palmitoylation sites using the composition of k-spaced amino acid pairs. Protein Eng. Des. Sel. 22:707–712. doi: 10.1093/protein/gzp055.

Wilson GM et al. 2005. The FIP3-Rab11 protein complex regulates recycling endosome targeting to the cleavage furrow during late cytokinesis. Mol Biol Cell. 16:849–860. doi: 10.1091/mbc.e04-10-0927.

Wirschell M et al. 2008. Building a radial spoke: flagellar radial spoke protein 3 (RSP3) is a dimer. Cell Motil Cytoskeleton. 65:238–248. doi: 10.1002/cm.20257.

Wuichet K, Søgaard-Andersen L. 2014. Evolution and Diversity of the Ras Superfamily of Small GTPases in Prokaryotes. Genome Biol Evol. 7:57–70. doi: 10.1093/gbe/evu264.

Xie Y et al. 2016. GPS-Lipid: a robust tool for the prediction of multiple lipid modification sites. Sci Rep. 6:28249. doi: 10.1038/srep28249.

Xu Y, Wang Z, Li C, Chou K-C. 2017. iPreny-PseAAC: Identify C-terminal Cysteine Prenylation Sites in Proteins by Incorporating Two Tiers of Sequence Couplings into PseAAC. Med Chem. 13:544–551. doi: 10.2174/1573406413666170419150052.

Yang Y-K et al. 2011. ARF-like protein 16 (ARL16) inhibits RIG-I by binding with its C-terminal domain in a GTP-dependent manner. J Biol Chem. 286:10568–10580. doi: 10.1074/jbc.M110.206896.

Yoon HS et al. 2017. Rhodophyta. In: Handbook of the Protists. Archibald, JM, Simpson, AGB, & Slamovits, CH, editors. Springer International Publishing: Cham pp. 89–133. doi: 10.1007/978-3-319-28149-0_33.

Yu C-J, Lee F-JS. 2017. Multiple activities of Arl1 GTPase in the trans-Golgi network. J Cell Sci. 130:1691–1699. doi: 10.1242/jcs.201319.

Zahn C et al. 2006. Knockout of Arfrp1 leads to disruption of ARF-like1 (ARL1) targeting to the trans-Golgi in mouse embryos and HeLa cells. Mol Membr Biol. 23:475–485. doi: 10.1080/09687860600840100.

Záhonová K et al. 2018. Extensive molecular tinkering in the evolution of the membrane attachment mode of the Rheb GTPase. Sci Rep. 8:5239. doi: 10.1038/s41598-018-23575-0.

Zhang Y et al. 2018. The unusual flagellar-targeting mechanism and functions of the trypanosome ortholog of the ciliary GTPase Arl13b. J Cell Sci. 131. doi: 10.1242/jcs.219071.

Zhou B, Cox AD. 2014. Posttranslational Modifications of Small G Proteins. In: Ras Superfamily Small G Proteins: Biology and Mechanisms 1: General Features, Signaling. Wittinghofer, A, editor. Springer: Vienna pp. 99–131. doi: 10.1007/978-3-7091-1806-1_5.

