## Supplementary figures for "A eukaryote-wide perspective on the diversity and evolution of the ARF GTPase protein family"

**Supplementary figure 1. Maximum likelihood phylogenetic tree of the complete ScrollSaw dataset.** The tree was inferred from 354 protein sequences using IQ-TREE with LG+I+G4 model (the model selected by the program itself) with the ultrafast bootstrap algorithm and the SH-aLRT test (both 10000 replicates), as described under Materials and Methods. Dots at branches represent bootstrap values as indicated in the graphical legend (top right).

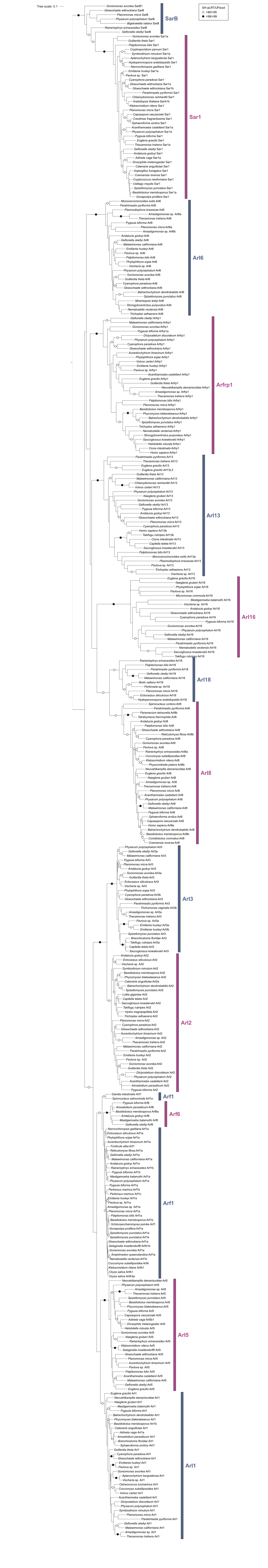

**Supplementary figure 2. Maximum likelihood phylogenetic tree of the ARF family based on a reduced ScrollSaw dataset**. The tree was inferred from 348 protein sequences using IQ-TREE with LG+I+G4 model (the model selected by the program itself) with the ultrafast bootstrap algorithm and the SH-aLRT test (both 10000 replicates). Dots at branches represent bootstrap values as indicated in the graphical legend (top right). Compared to the tree in Supplementary figure 1, this analysis omits most Metamonada species (except for *Paratrimastix pyriformis*), as their sequences are generally very divergent and may negatively impact the tree inference.

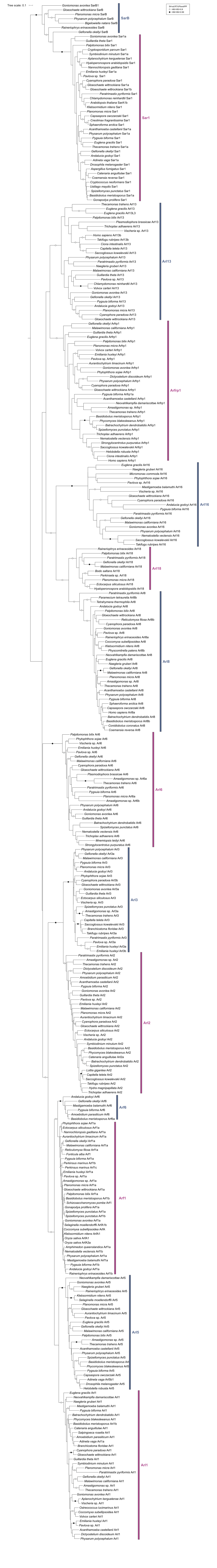

**Supplementary figure 3. Phylogenetic analysis of Arf1 and Arf6**. The tree shown is a result of a ML analysis of a subset of 101 sequences of the reduced "scrollsawed" dataset restricted to Arf1, Arf6, Arl1 and Arl5 sequences. Arl1 (34 sequences) and Arl5 (26 sequences) sequences were used as an outgroup according to the topology shown in Figure 1. The alignment was trimmed with trimAl. The tree was inferred using IQ-TREE with LG+I+G4 model (the model selected by the program itself) with the ultrafast bootstrap algorithm and the SH-aLRT test (both 10000 replicates). Dots at branches represent bootstrap values as indicated in the graphical legend (bottom left).

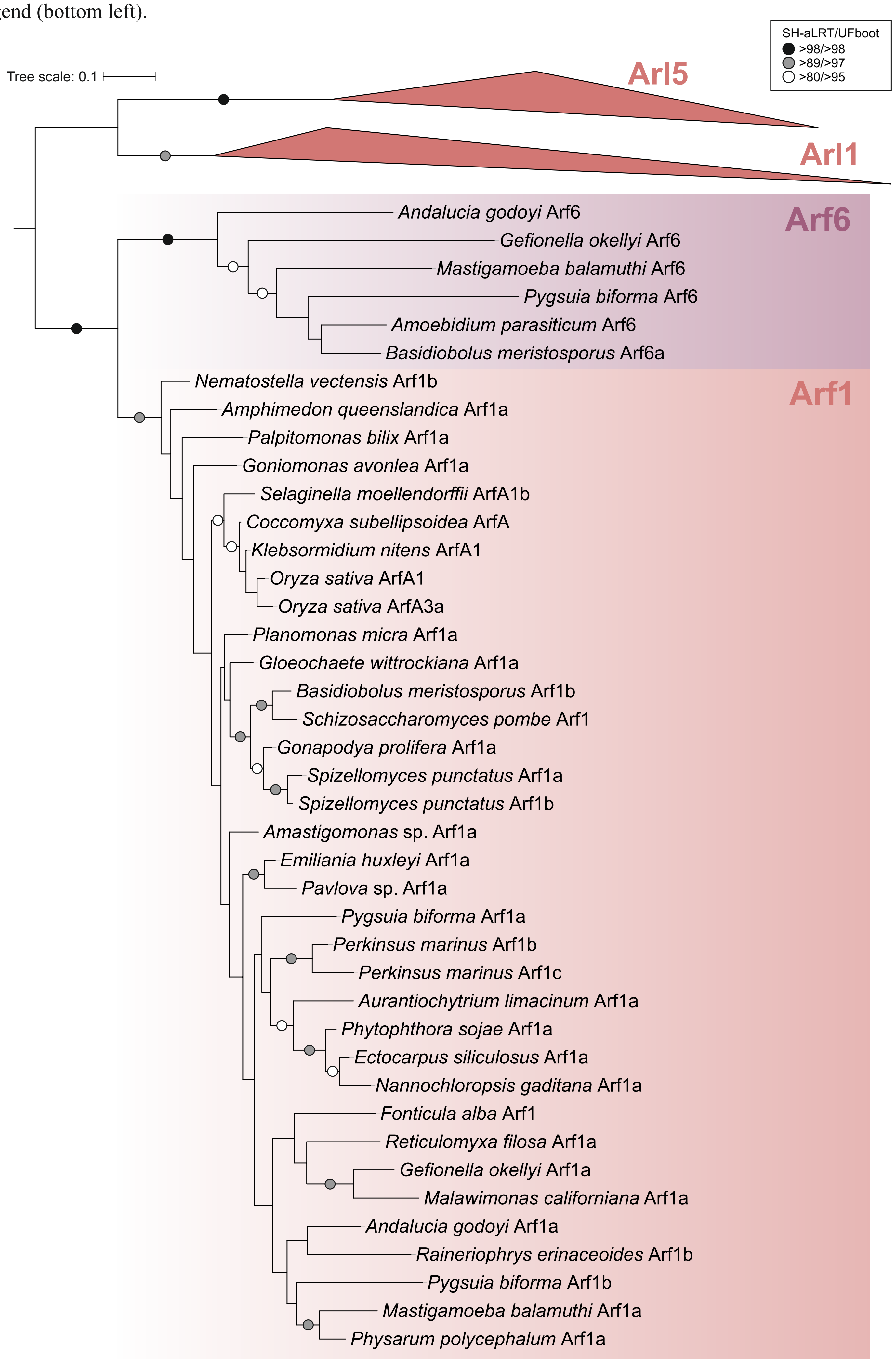

**Supplementary figure 4. Position and phase of introns in Arf genes mapped onto an alignment of Arf protein sequences.** The intron positions are marked by highlighting the amino acid residues encoded by a codon located immediately upstream of the intron (phase 0; in red) or interrupted by the intron at the second or third position (phases 2 and 3; in green or blue, respectively). The identity of the sequences included is provided in Supplementary table 1. Note that genes with the coding sequence contained in a single exon and sequences represented only by transcriptomic data are not included in the alignment as indicated in Supplementary table 1, column G. Note that for simplicity the alignment regions corresponding to the N- and C-termini of the protein sequences were trimmed and species-specific insertions were omitted (each is represented by three red dots).

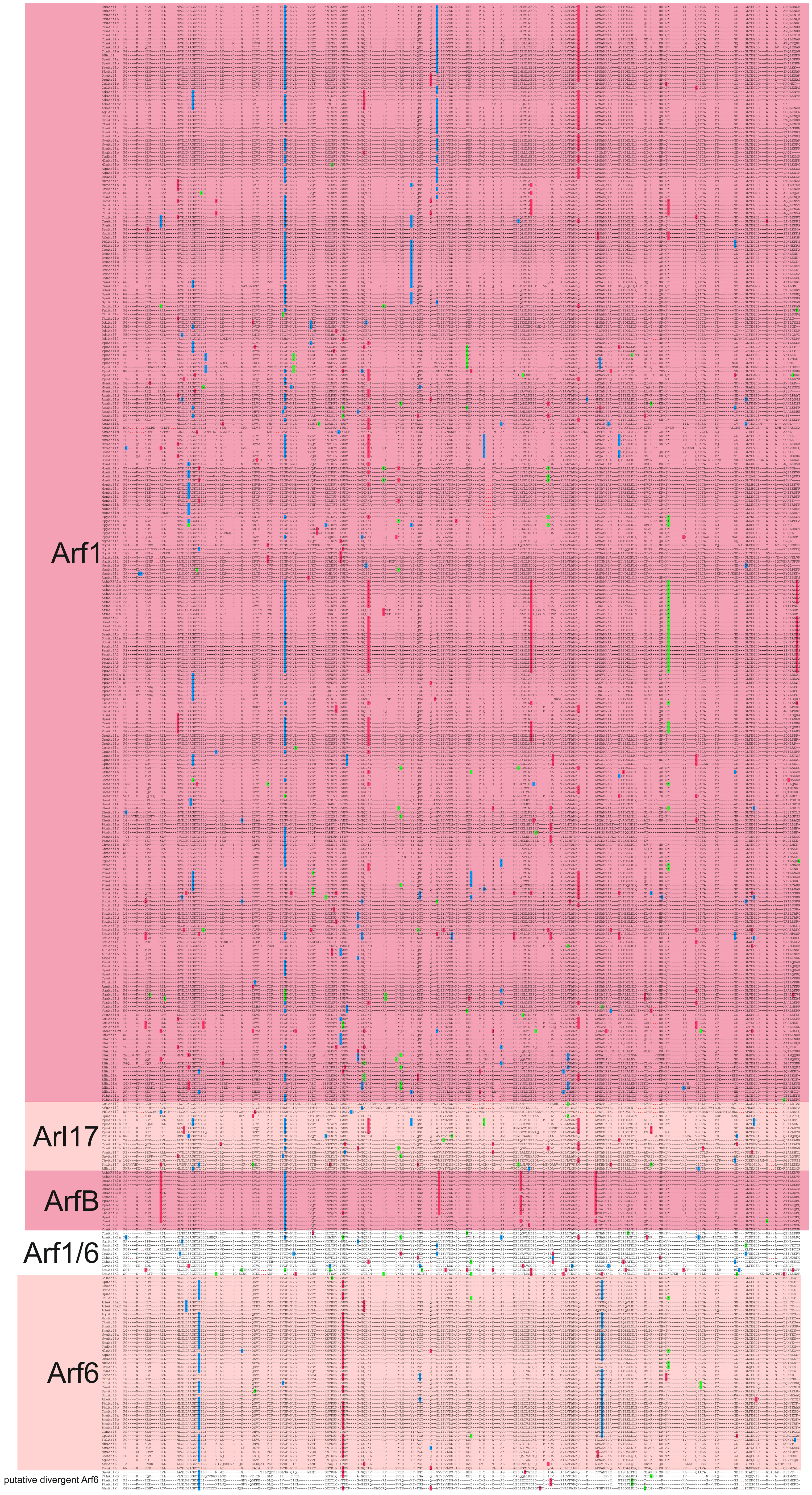

Intron position specific to:
- Arl8
- Arl18

Insertion in sequence

**Supplementary figure 5. Position and phase of introns in Arl8 and Arl18 genes mapped onto an alignment of Arl8 and Arl18 protein sequences.** The intron positions are marked by highlighting the amino acid residues encoded by a codon located immediately upstream of the intron (phase 0; in red) or interrupted by the intron at the second or third position (phases 2 and 3; in green or blue, respectively). The identity of the sequences included is provided in Supplementary table 1. Note that genes with the coding sequence contained in a single exon and sequences represented only by transcriptomic data are not included in the alignment as indicated in Supplementary table 1, column H. Note that for simplicity the alignment regions corresponding to the N- and C-termini of the protein sequences were trimmed and species-specific insertions were omitted (each is represented by three red dots). The specific, broadly conserved intron positions of Arl8 and Arl18 are indicated by violet (top) and pink (bottom) arrows, respectively. The two sequences from Metamonada species (MonArlX2 and TpyArlX2) are putative divergent Arl8 paralogs.

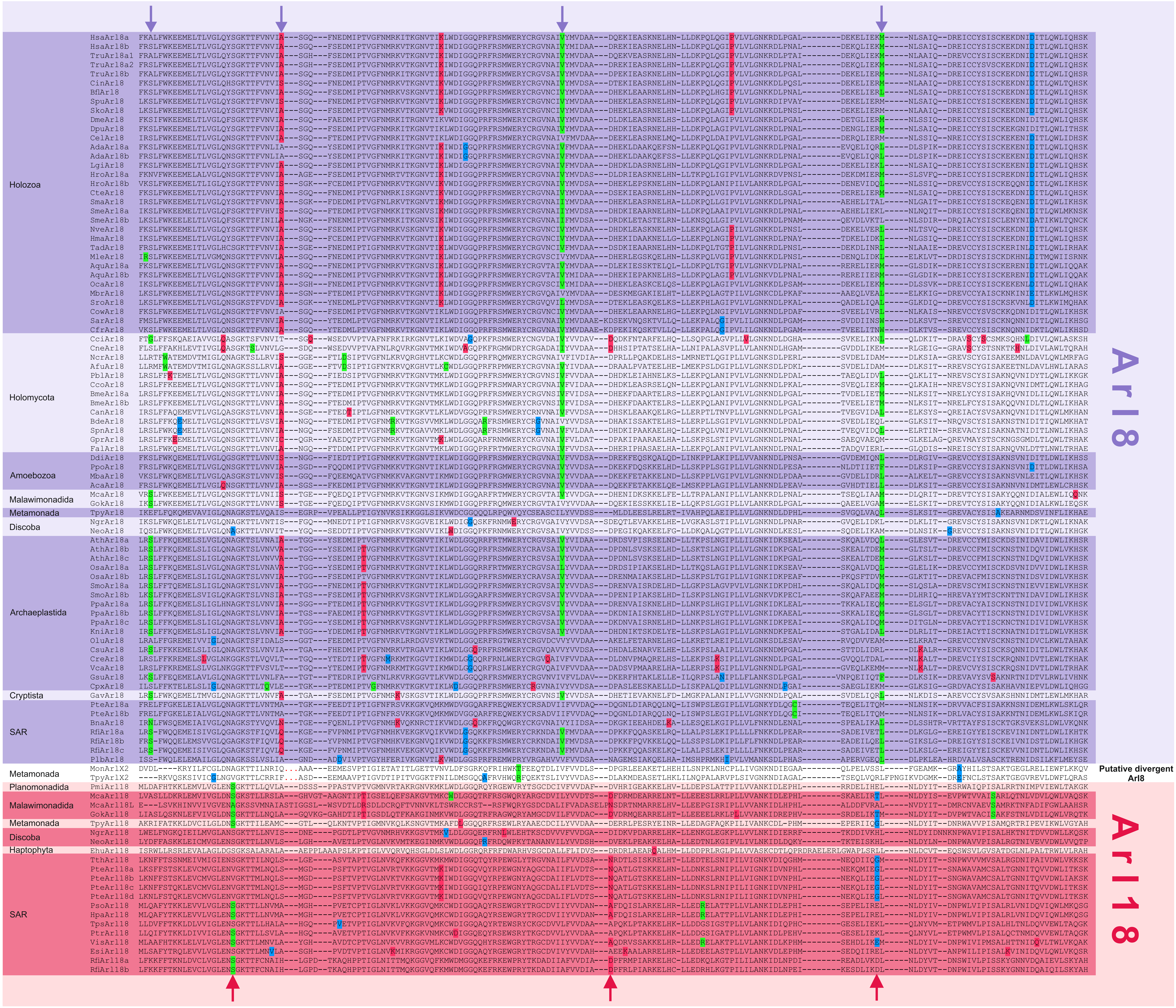

Intron position specific to:
Sar1
SarB
Insertion in sequence

**Supplementary figure 6. Position and phase of introns in Sar1 and SarB genes mapped onto an alignment of Sar1 and SarB protein sequences**. The intron positions are marked by highlighting the amino acid residues encoded by a codon located immediately upstream of the intron (phase 0; in red) or interrupted by the intron at the second or third position (phases 2 and 3; in green or blue, respectively). The identity of the sequences included is provided in Supplementary table 1. Note that genes with the coding sequence contained in a single exon and sequences represented only by transcriptomic data are not included in the alignment as indicated in Supplementary table 1, column I. Note that for simplicity the alignment regions corresponding to the N- and C-termini of the protein sequences were trimmed and species-specific insertions were omitted (each is represented by three red dots). The specific broadly conserved intron positions of Sar1 and SarB are indicated by violet and pink arrows, respectively. Some introns specific to SarB are marked even though some Sar1 genes share introns at the same position (most likely due to an independent gain).

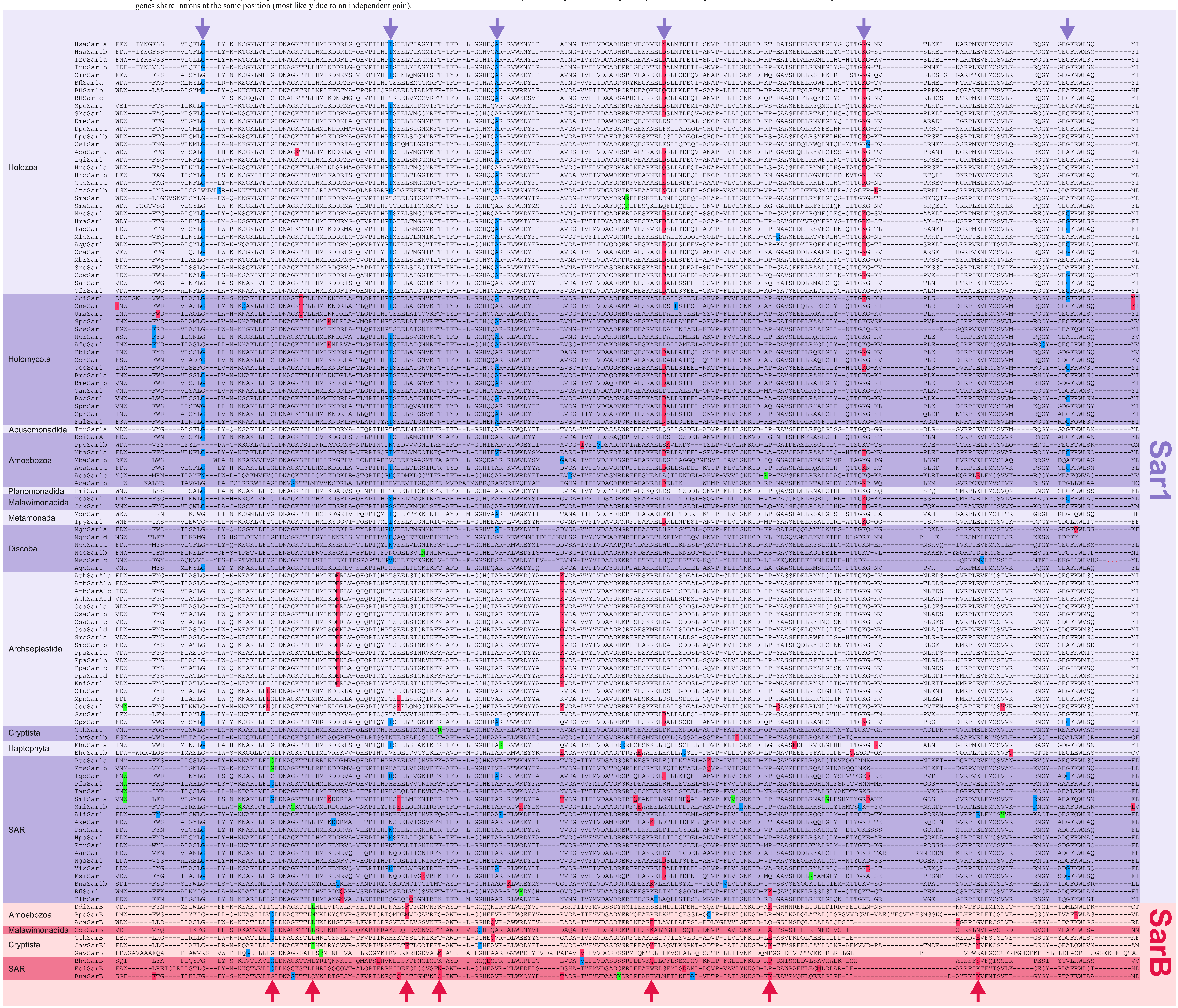

**Supplementary figure 7. Sequence logo of the Walker B (also known as the G3) motif of Sar1 and SarB proteins**. Sequence logo of Sar1 and SarB was obtained from multiple sequence alignments of all Sar1 and SarB protein sequences. The sequence logos are restricted to the Walker B motif as defined by Leipe et al., (2002), i.e. the motif hhhhDxxG (where h is a hydrophobic residue).

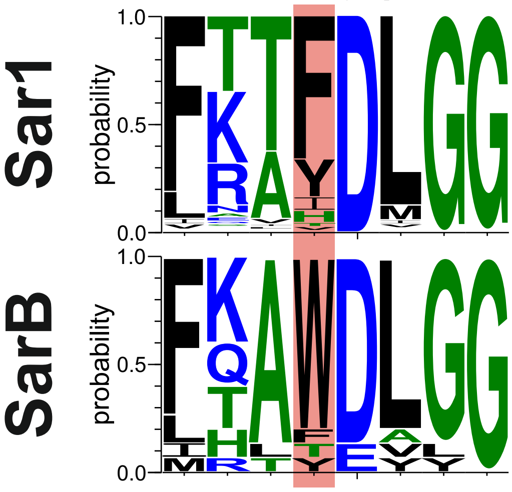

**Supplementary figure 8. Multiple sequence alignment of the novel domain occurring at the C-terminus of Arl17 proteins.** Depending on the protein, the domain occurs in one, two or three tandemly arrayed non-identical copies, labelled as novel domain (ND) 1 to 3. The alignment includes non-conserved regions flanking the domain copies (i.e. a part of the linker connecting the GTPase domain to the novel domain, linkers separating the domain copies, or the very C-terminal extension). The second paralog of Arl17 from *Chromera velia* (ChvArl17b) is not included due to incompleteness of the sequence. The consensus sequence, (residues conserved in at least 80% of the sequences included) is shown at the bottom.

```
Adineta vaga Arl17a_ND1                      ----ETNKSLQWLSQ--EDDDTDEEFID----KFQKH----Q----LLS---EQ-F--DHR----SFLRTIWSYLQL----HNRK---ETIKAIFNHLPTY
Adineta vaga Arl17a_ND2                      (1)-------------DEIDDDDDEFLK----QFESC----T----LT-----S-W--SHR----THVRMAWLYLTR----DGRR---TGVKKIFDGIKNF
Adineta vaga Arl17a2_ND1                     ----ETNKSLQWLSQ--EDDDTDEEFIG----KLQKH----Q----LLA---EQ-F--DHR----SLLRTIWSYLQL----HNRK---EMIKAVFDHLPTY
Adineta vaga Arl17a2_ND2                     (1)-------------DEIDNDDDEFLK----QFESC----I----LT-----S-W--SHK----THVRITWLYLTR----DGRR---TGVKKIFDGIKNF
Adineta vaga Arl17a3_ND1                     ----ETNKSLQWLSQ--EDDDTDEEFID----KFQKH----Q----LLA---EQ-F--DHR----SLLRTIWSYLQL----HNRK---EMIKAVFDHLPTY
Adineta vaga Arl17a3_ND2                     (1)-------------DEIDNDDDEFLK----QFESY----K----LT-----S-W--SHK----THVRITWLYLTR----DGRR---TGVKKMFDGIKNF
Adineta vaga Arl17b_ND1                      ----EAQKSLEWLSQ--VDDDTDEEFID----KFQKH----Q----LLS---EQ-F--DHR----SFLRTIWSYLQL----HNRK---ETIKAIFNHLPTY
Adineta vaga Arl17b_ND2                      VV-------------EIDDDDDDEFLK----QFESC----T----LT-----S-W--SHR----THVRMAWLYLTR----DGRR---TGVKKIFDGIKNF
Coprinopsis cinerea Arl17_ND1                ----LSKRLETWLHQ-TEVDSPPDEFIS----QFKSY----S----LP-----S-W--DHY----THIRIAYLLLMA----HGRQ---KGKDMVFSGLENY
Aspergillus fumigatus Arl17_ND1              ----LQRRIEKEVT---GDTIDPDAFWT----SFLEG----D----LP-----A-W--THY----TYLKALYFVILE----SAKK---KTFTEIANDFNIH
Aspergillus fumigatus Arl17_ND2              -----------------------------------------------------------DEG(2)-KLIRYAFAVMQY(4)GAARG---QVVTQALVALQQA
Aspergillus fumigatus Arl17_ND3              (17)PKLPSAEELTF--RARMACEELTP----TLQSS----G----PP(1)----L--SHT----HLLFYLHKRFTQSAEGGTRKKLERRARELFSEIA--
Phycomyces blakesleeanus Arl17_ND1           ----ARMPVAPWEDLPNPHHFKDKEFCE----WFLQAK---A----FL(1)----F--DHY----SLLRIAYLSILQ-DKQDTRRLL-YSLRTILRGIELL
Spizellomyces punctatus Arl17_ND1            ----LSTKLEEWLL---RQDDPDDVFLS----QLDAC----T----LD-----T-W--DHY----THLRIAYLYLTR----EGRQ---KGKNLIFDKIEHF
Physarum polycephalum Arl17a_ND1             ----QEALFLKWIV---PDKDTTDEFLA----KLEAG----E----IV-----L-F--DHR----TLLRTIWCYIST----SGRR---VGIKKIFEQLSKY
Physarum polycephalum Arl17a_ND2             YD-------------VDDKLTDDEFLA----NFEVC----N----LH-----K-W--DHY----THLRVAWSYITK----FGRK---EGSDKIFAGIENF
Physarum polycephalum Arl17b_ND1             ----NETVFLKWLS---DDKDTTEEFLA----KLEAG----E----LT-----E-W--DHR----TLLRTVWSHITV----DGRR---NGIKKIFANLEKY
Physarum polycephalum Arl17b_ND2             LD-------------EEDDFDDDEFLA----RFEKC----D----LK-----K-W--DHR----THIRMGWAYLTK----YGRK---EGSEKILKGIENF
Physarum polycephalum Arl17c_ND1             (5)GVRPPPQPPFTS-PSDALTNNEFLK----AFAEY----T----LP-----S-W--DHK----THIRIAYLNLKR----HGRR---EGVRKIRQGLQDF
Physarum polycephalum Arl17d_ND1             (7)GIRPPPQPPFTS-PSDVLTNTQFLK----AFEEY----T----LP-----S-W--DHK----THIRIAYLNLKR----YGRR---EGVRKIRQGLQDF
Planomonas micra Arl17a_ND1                  ----SAAVIDRWWAW--HRELDDATFVD----RIEHAAEA-D----LNR---EH-C--NHL----VLLRFIHAHLHS(10)SGRK---AAVDAIFAGLERF
Planomonas micra Arl17b_ND1                  ----TKETIAAWRKS--HADLSDAEFQL----RVERPQDVPD----LK-----R-WFMNHM----VLIRYIHMHLLA(8)HRGRK---TAVDAIFAGLQQF
Klebsormidium nitens Arl17_ND1               ----MEALLQQWLE---QEDEPDDEFLD----KFRTY----T----LD-----S-W--DHR----THLRIAWLYLTR----EGRR---EGMRLIFEGIKAF
Klebsormidium nitens Arl17_ND2               -------------------SENDADFLA----RFEGG----Q----LD-----R-W--NHE----CLIRVIFLYLIT----LGRA---QGKEKILTELRRH
Chlamydomonas reinhardtii Arl17a_ND1         ----EMDVLERWLQ---VEDEPDDEFLR----KLEDY----S----LD-----V-W--DHR----THLRLAWLYLTR----HGRR---EGLARIHSAIQSF
Chlamydomonas reinhardtii Arl17a_ND2         -------------AA-PPIALSDDEFLQ----RFLGR----G----LD-----R-W--SHA----TMLRAVYCCLRA----HGRR---HGGKLALDALAAL
Chlamydomonas reinhardtii Arl17b_ND1         ----AEERLSGWLAEPVAAGDSDDAFLA----ALEAC----E----PG----VR-W--DHF----AHLRLAWLYLVR----YGRS---QGAARIFQAVQRY
Chlamydomonas reinhardtii Arl17b_ND2         -------------AAAAA-AARDEDLLA(44)NFEGR----G(16)LT-----G-W--GSD----TLPRVAYAAIRH----HGYA---RASELLQAAAAAA
Volvox carteri Arl17_ND1                     ----ELDVLERWLS---AEDEPDEEFLM----KLENY----S----LD-----C-W--DHR----THLRLAWLYLTR----LGRR---EGLARIHASIRAF
Volvox carteri Arl17_ND2                     -------------------PLTDAEFLD----RFLGR----S----LD-----R-W--NHE----CLLRAVYCCLKR----HGRR---RGGDVALDGLRAL
Gloeochaete wittrockiana Arl17_ND1           ----SQAKFKQYLSV-PTFSMPDEDFLK----AFQEAS---S----LP---LDH-F--NHA----AKIRMMWLYLSK(4)SQGRR---TAVDLILNGLSNY
Chromera velia Arl17a_ND1                    ----IAEVLEDWLS---REDEEAEVFLA----HLENF----T----LE-----S-W--DHY----THLRIAWLLINT----HGRR---EGFRLIFARIEAF
Chromera velia Arl17a_ND2                    -------------------SVSDSLFLQ----LFEGN----R----LP-----S-W--GHA----AFIRLIYCRLKT----QGRR---KATDSLFASLKNL
Symbiodinium minutum Arl17_ND1               ---EPVQIEDRKLSE----SLEDEAFLQ----TFFDQ----T----LK(92)EATW--AHM(35)SLLRLSFILLKA----MPRK---EAIQRMDVILKRL
Symbiodinium minutum Arl17_ND2               -------------------QLTDEDFLQ----AVQKQ----S----LT-----S-W--GLP----SLARLAYVYLTT----MGRR---QAVRQLLEDVENL
Reticulomyxa filosa Arl17_ND1                ----KKSKLEEWVE---RKDEDDDVFLE----KFENV----T----LD----GP-F--DHY----VHLRLAYLYFTK----FGRR---QGLQKIFSNLQNF
Reticulomyxa filosa Arl17_ND2                ------------------EDTDDKTFIK----LFETK----Q----LQ-----G-W--NHL----YLLRAIWYYLET----IGRK---EGKNKIFSEIKRH
Raineriophrys erinaceoides Arl17_ND1         ----EKSLLEQWLD---RSDEPDDVFLS----KLDDF----S----LS-----S-W--DHY----THLRIAWLLLSR----HGRQ---VGMPMIFSKIKKF
Raineriophrys erinaceoides Arl17_ND2         -------------------AISDEEFLE----MFEKR----T----LK-----S-W--GHQ----SMIRLVFLLLSK----HGRQ---KGVQIILDTLQHF
Goniomonas avonlea Arl17_ND1                 ----RGLAKAQELS---APKLSDKELMT----GFTNGHA--A----LPN---PG-L--THE----TLLRFMWLELKQ(4)SRGRK---QAVDCILQGVEAA
Palpitomonas bilix Arl17_ND1                 ----IGEKMERWISE-CESDCSDADFIE----SVENY----T----LA-----N-W--DHR----VHLRLAWILLTT----LPRK---QAKDKIFELIGNF
Palpitomonas bilix Arl17_ND2                 LDL--------------GDPEDAESYIQ----AFKEK----R----LS-----A-W--GHR----PYLRLLYAFLET----RGFRGS-KAVDDAHAAISHG
Consensus/80%                                .....................ssp.Fl......bp.....p....L........a..sHb....sblRhhahblp......tR+...pthp.lb..lp.b
```

```
Adineta vaga Arl17a_ND1                    IND------------------INETLTYFWIQIVHY---AREATKNP----------------TND-----FPGFL----LMNPQILNEAELPLAYYKKETL-
Adineta vaga Arl17a_ND2                    LEN----TQIS-----SKPTFHFTMTYFWIQMIDL---AIAQSPK-----------------ETT-----FEEFL----RLNPQLL-KEDLYFDYYKKETI-
Adineta vaga Arl17a2_ND1                   IND------------------MNETLTYFWIQIVHY---AREATKNP----------------TND-----FPGFL----LMNPQILNETELPLVYYKKETL-
Adineta vaga Arl17a2_ND2                   LEN----TQIS-----SKPAFHFTMIYFWIQMIDL---AIAQTPK-----------------EIN-----FEEFL----RLNPQLL-KEDLYLDYYKKQTI-
Adineta vaga Arl17a3_ND1                   IND------------------MNETLTYFWIQIVHY---AREATKNP----------------TND-----FPGFL----LMNPQILNETELPLVYYKKETL-
Adineta vaga Arl17a3_ND2                   LEN----TQIS-----SKPAFHFTMIYFWIQMIDL---AIAQTPK-----------------EIN-----FEEFL----RLNPQLL-KEDLYLDYYKKQTI-
Adineta vaga Arl17b_ND1                    IND------------------INETLTYFWIQIVHY---AREATKNP----------------TND-----FPGFL----LMNPQILNEAELPLAYYKKETL-
Adineta vaga Arl17b_ND2                    IAN----SQVS-----RKTTFHFTMTYFWIQMIDL---AIAQSPK-----------------EIG-----FEEFL----RLNPQLM-NGGLFLEYYKKETM-
Coprinopsis cinerea Arl17_ND1              IAN----NSKT-----NVRGFHFTMTYFWIQIVHF---GIRNMPPQLLDHPLE(6)VRYPSSAD-----FFRFL----FINPHLV-DGSLWTDYYSKEVM-
Aspergillus fumigatus Arl17_ND1            LIR(11)SETPSAY--VHAPFNICLATFWTLQLQH---GIREYRMHSMSS--------HLPSREE-----FPQVL----RRSPSLM-STYLWKSYYSFNPV-
Aspergillus fumigatus Arl17_ND2            TMR----ARTADS---TVETYSETQAYFWIQIVHA---ALRSLDDKK-----------GSVDTSE---MSFEAFQ(5)----------KPTDWQEYYSKKLW-
Aspergillus fumigatus Arl17_ND3            -----------------GPIVAGATYRNFWIQQVGV---AVLNSDIGK-------------GRST-----FPEFI----TSNLHLV-FEELHGIYYGPGVW-
Phycomyces blakesleeanus Arl17_ND1         EQDALRDTGLANR(7)ESIEYSETQTLFWIQMVSF---ALLRHPVLD-------------GEERD-----FESFL----MGCPELW-DGQGWKKYYSPKIY-
Spizellomyces punctatus Arl17_ND1          IKH----SPRT-----NGKTFHLTLTYFWIQLVNY---GIRSMKPP-----------------PEE-----FKQFL----VMNPHLA-DGNLPLEYYSKETL-
Physarum polycephalum Arl17a_ND1           YTK----AP--------TIKNNETWTYFWIQMVHY---AMEVTRNP----------------SGD-----FIGFL----FMNPLMM-NQRMPYGYYSKKVI-
Physarum polycephalum Arl17a_ND2           IKN----SKIS-----RKTTFHLTMTFFWIQMVDI---AIQSSPP-----------------NIP-----FDEFM----KLNPQLM-DGGLFLQYYKKETM-
Physarum polycephalum Arl17b_ND1           YEK----FP-------LTLRNNETWTYFWIQLVHY---TMQTTRNP----------------SDD-----FVGFL----FMSPALM-NAHMPFGFYKKKTI-
Physarum polycephalum Arl17b_ND2           INN----SKIS-----RKTTFHLTMSLFWIQMLDY---YICSSPK-----------------GIS-----FEEFL----QKNPQVM-EGGLFLQYYKKETM-
Physarum polycephalum Arl17c_ND1           IAN----SSIT-----NKNRYHETMTYFWIQMTHY---AMSSCKIPLIWEPYD----DENGLPE-----FNAFL----DKNVHLM-NGGLFLEYYTRDLI-
Physarum polycephalum Arl17d_ND1           IAN----SAIT-----NKNRYHETMTYFWLQMTHY---AMSTCKIPLTWEPFD----DENGLPE-----FNTFL----DKNVQLM-NGGLFLEYYSRDLM-
Planomonas micra Arl17a_ND1                HAH-------------VQLVSHTTQIYFFIQMVDL---AMQLGNREL-------------ADAS-----FADLV----AARPWLL-DDQLIHSYYSPKLV-
Planomonas micra Arl17b_ND1                NSH-------------FDRVHHIT-SYVCPWLPTA---PVRNSDHPC-------------TAST-----FGS-----------------------------
Klebsormidium nitens Arl17_ND1             IAN----SPRTKRA--RGTTFHETMTYFWVHMVHY---ALATTSNP----------------DGG-----FKTFL----LLNPQLA-NGGMFLAYYSKKLM-
Klebsormidium nitens Arl17_ND2             -----------------EGSGFHMTINYFWIQMVDF---ARATWQKSRQGGPAPG---GVRTKKD(27)AFKSWCQGWKAGEGILT-NSRLYLDHYREKSI-
Chlamydomonas reinhardtii Arl17a_ND1       IAN----SPVTKRK--TGTTYHETMTYFWAHMVHF---CIASMKAPQ-----------QQGKEPD-----FKTFL----LFNPLLT-NGGLFLHYYSKDLM-
Chlamydomonas reinhardtii Arl17a_ND2       -----------------QGEHAHTTLNYFWLTMLTH---TLAAEHSAALFADRP(39)AAAAAAA(116)WSALL(6)SVRLQELVADESRYLAHYNKKTI-
Chlamydomonas reinhardtii Arl17b_ND1       IQH----GVAGGG---GGRTFHTTMTFFWMHMVHY---ALASSQLHT-----------PMVRTS-----FRAFL----IANPYLA-DSGLFLRHYSRARM-
Chlamydomonas reinhardtii Arl17b_ND2       ATT----RGQAST---EESSARGTLE---------SPPALAEAAAAASAETRP----HVKRAGD(71)EEGEWL(6)AWGVVTALNDKALRVD--------
Volvox carteri Arl17_ND1                   IQH----SPLTARR--SGTTYHETMTYFWAHMVHF---AIASQKVP-------------AGASPD-----FRVFL----LMNPQLT-NGGLFLHYYSKQLM-
Volvox carteri Arl17_ND2                   -----------------QGPDFHLTICYFWLCLLTH---TLASEYHAALFTDRP-------AKAA(22)EWPELL(6)AVRLKELVADERRYLHYYSNKVV-
Gloeochaete wittrockiana Arl17_ND1         YKS-------------KNLVYHMTLSYFWLQLVDL---AINAPYYNPPLTTRL(6)-SVPTVEA-----FETFV----LSNLFLL-NEDLVTDFYSINAI-
Chromera velia Arl17a_ND1                  IKN----SELTKRAD-RQTTFHQTMTYFWAHMIHY---ALEAARDPR-------------AKTD-----FKFFL----AFNPQLC-NSGMFLHFYAKERM-
Chromera velia Arl17a_ND2                  -----------------QKGAFHETKTYFWIQMVTH---ALATDFHACIFDSSSDK--EEKAGPE--MPEFEDLM(13)SKLRELTVDTQFFHAFYSKKLI-
Symbiodinium minutum Arl17_ND1             EPT----------------RPFHETRQYVALQFAHL---ALVQHPVL--------------QEKS-----FADLQ----ERCPELC-EEDCIYKYYSEQAL-
Symbiodinium minutum Arl17_ND2             RGS-----------QVVHGVFTHETLAYFTLHMIHY---FIASQKLS--------------MSEP-----FGGFV----KKYDKIL-DLSLYRSYYSDQAI-
Reticulomyxa filosa Arl17_ND1              LTK----SKN-------TRNTFSITITYFWAHMVWY---ALEATKIG---------------KDN-----FKTFL----VMNPRLS-DFGLYKEYYSDDLF-
Reticulomyxa filosa Arl17_ND2              -----------------DGDAYHETLTYFWLQMVDF(4)AVELNTNKT-------------TIST-----FTQWI(19)STSYDLE-DVLLWKLFYSESLL-
Raineriophrys erinaceoides Arl17_ND1       IEN----SSRTKRS--HGTTFHETLTYFWVHMVHY---AIVSTQNP----------------NGQ-----FKTFL----LLNPQLS-NGGMYLAYYSKKLI-
Raineriophrys erinaceoides Arl17_ND2       -----------------QGASFHFTLTYFWIQMVHI---CVATVSRSR----------SLNSFAD-----FSSAA----ETATVLN-DSNYYLQFYSVNVI-
Goniomonas avonlea Arl17_ND1               SRA-------------AGAIFHLTHTYFWLQLVDM---SLQNDIKA---------------ELGR-----FVEFL----GAAPWLL-DHELITRFYSRSLI-
Palpitomonas bilix Arl17_ND1               IAN----SKIA-----SKTRFHLTMTYFWIQMVHF---AIMTTENK---------------DGT-----FQSFI----ILNPQLA-NGSLFLEYYSADLM-
Palpitomonas bilix Arl17_ND2               YP--------------RTRQYYFTFIYFWIHFLRI(4)AMKERNGGA-------------REQD---PTFAEVWE---MGITTLG-NSNLLQDYYSTPIL(
Consensus/80%                              ...................bpbTbsYFWlpblpb...tb.......................s.....F..bl.......sblh.p..b.b.aYp.phb.
```

```
Adineta vaga Arl17a_ND1                  --Y-SNQAK-GSVILADV--KQLPSILPTANKAVSKVTSGRE------------(3)-
Adineta vaga Arl17a_ND2                  --LNNPTAR-QEMVLPDI--KPLPTLVVQQTKK-------------------------
Adineta vaga Arl17a2_ND1                 --F-SSQAK-ISVVLADV--KQLPSILPTAYNPVSKAISRRE------------(3)-
Adineta vaga Arl17a2_ND2                 --LNNSTAR-QEMILPDI--KPLPTLVIPQTKK-------------------------
Adineta vaga Arl17a3_ND1                 --F-SSQAK-ISVVLADV--KQLPSILPTAYNPVSKAISRTD------------(3)-
Adineta vaga Arl17a3_ND2                 --LNNPTAR-QEMILPDI--KPLPTLVIPQTKK-------------------------
Adineta vaga Arl17b_ND1                  --Y-SNEAK-ASVILADV--KQLPSILPTANKAVSKVTSGRE------------(3)-
Adineta vaga Arl17b_ND2                  --LNNPTAR-QEMVLPDI--KPLPTLIPATTGK-------------------------
Coprinopsis cinerea Arl17_ND1            --M-SPEAK-SSMVLPDK--KPLPSLVIRDAIAS------------------------
Aspergillus fumigatus Arl17_ND1          -----SRPR-DYWSIPNL--RKLPTQTDYLRDPATVPRK-------------------
Aspergillus fumigatus Arl17_ND2          ---NSVAAR-SQFVMPDL--KPLPNVIASLPSK-------------------------
Aspergillus fumigatus Arl17_ND3          ---TSADAA-EKILGPDR--RRMETIVNMADVNMSANTK-------------------
Phycomyces blakesleeanus Arl17_ND1       --L-SLKAA-QEFIPPDR--KPLPNAFKASSLALRG----------------------
Spizellomyces punctatus Arl17_ND1        --FMDPKAR-TEMVLPDK--KPLPSVIPRETAKRRR----------------------
Physarum polycephalum Arl17a_ND1         --L-GTEAV-KNVVLPDK--RPLPSIVPKAYNPAAHSPAPPQ-STNSTDQNSTT(17)
Physarum polycephalum Arl17a_ND2         --LNNIVAR-KEFVLPDI--KPLPTYVPKKQ---------------------------
Physarum polycephalum Arl17b_ND1         --L-SPEAA-KAVVLPDK--RPLPSIVPKALIPAS---------LKTIDQSLKKLE--
Physarum polycephalum Arl17b_ND2         --LNNPIAR-KEFVLPDV--KPLPSFVPGTK---------------------------
Physarum polycephalum Arl17c_ND1         --LNTQSSR-EQFAIPDV--KPLPNIVETN----------------------------
Physarum polycephalum Arl17d_ND1         --LNTQSTR-EQFTIPDI--KPLPNIVEAS----------------------------
Planomonas micra Arl17a_ND1              --FHTPDAV-SSFVLPDI--KQLPSVLNFNVPVSVQ----------------------
Planomonas micra Arl17b_ND1              ------------------------------------S---------------------
Klebsormidium nitens Arl17_ND1           --LHTPEAR-TSVVLPDK--APLPSLVSDVRERSSALEKTA--GISPAAESSST(6)-
Klebsormidium nitens Arl17_ND2           --F-DTSAA-EVFRLPDL--KPLPSLVG------------------------------
Chlamydomonas reinhardtii Arl17a_ND1     --LKNPEAR-KQVVLPDK--RPLPSLVTSVESIKQLQQNQARYGKKPAAGSGPA(8)-
Chlamydomonas reinhardtii Arl17a_ND2     --F-SDAAA-AGFVPPDK--KPLPTTV-------------------------------
Chlamydomonas reinhardtii Arl17b_ND1     --LHDPAAR-TALLLPDL--QPLPSLVTDVEARKREQAREKRVAVVAGAAAAAV(85)
Chlamydomonas reinhardtii Arl17b_ND2     ----GEDAELEKMLGSDR--VPLMRAALPQYR--------------------------
Volvox carteri Arl17_ND1                 --LQSPESR-IRVVLPDK--RPLPSIITDLSTLRR--------GTTTSAASQAPG---
Volvox carteri Arl17_ND2                 --F-LAAAA-TTFVPPDK--RPLPSTT-------------------------------
Gloeochaete wittrockiana Arl17_ND1       --FHKPESL-AEFVLPDK--KPLPSILSFSRDNISSTSKSL--GI-------------
Chromera velia Arl17a_ND1                --LNDPKAR-TEVVLPDR--KPLPSLISSIEREGPLP---------PAHEHLPVGD--
Chromera velia Arl17a_ND2                ---ESKEAE-ESFVLPDK--KPLPSFAAPR----------------------------
Symbiodinium minutum Arl17_ND1           -----SEGQ-SSFRAPDL--QPLPTKLDQPLPTWHE----------------------
Symbiodinium minutum Arl17_ND2           ---HCSEAR-VSFVLPDL--CPLPDLLPTR----------------------------
Reticulomyxa filosa Arl17_ND1            --LKNAKSR-EEFMFPDKKDKQLPSTVDTNLQELKEAKKGI--DVIAQLKQLAK----
Reticulomyxa filosa Arl17_ND2            --F---GKA-TKLMLAKIWSCQISCLCPACRSNKSYI---------------------
Raineriophrys erinaceoides Arl17_ND1     --METPEAR-TQVVLPDK--RALPSLLSDVSQIPAKP-----IEVRTAPK--------
Raineriophrys erinaceoides Arl17_ND2     ---DSPAAA-TQMVLPDK--KPLPSIL-------------------------------
Goniomonas avonlea Arl17_ND1             --MSNPSTL-TEFVLPDL--QPLPSIVNFGAAAR------------------------
Palpitomonas bilix Arl17_ND1             --LKNPSSR-KEFALPDK--KQLPSIVPSAALPWKNRERFE--EVVSAMQSGTKN---
Palpitomonas bilix Arl17_ND2             7)VSLQDVD-KTFVLPDL--KKLPQSI-------------------------------
Consensus/80%                            ....s..t...phhbPDb..+.LPshl...............................
```

**Supplementary figure 9. Multiple alignment of the GTPase domain of Arl17 protein sequences together with various “standard” ARF family protein sequences from *H. sapiens*.** Alignment of the N-terminal part of Arl17 proteins sequences (i.e. the region containing the GTPase domain) and several ARF family sequences from *H. sapiens* included for a reference. The GTPase domain is delineated by greater/less than symbols, conserved G-boxes are delineated by stars. Note that the available sequence of Arl17 from *R. erinaceoides* is truncated (incomplete) at the N-terminus.

```
                                                         G1 box                    G2 box                     G3 box
                                <<<                     ********                     *                        **  **
H. sapiens Arf1          ---MGNIFANLFK---GLFGK--KEMRILMVGLDAAGKTTILYKLKLG---EI--VTTIPTIGFNVETVEYK---NISFTVW---DV---GGQDKIRP---LW
H. sapiens Arf6          -------MGKVLS---KIFGN--KEMRILMLGLDAAGKTTILYKLKLG---QS--VTTIPTVGFNVETVTYK---NVKFNVW---DV---GGQDKIRP---LW
H. sapiens Arl1          ---MGGFFSSIFS---SLFGT--REMRILILGLDGAGKTTILYRLQVG---EV--VTTIPTIGFNVETVTYK---NLKFQVW---DL---GGQTSIRP---YW
H. sapiens Arl2          -----MGLLTILK---KMKQK-ERELRLLMLGLDNAGKTTILKKFNGE---DI--DTISPTLGFNIKTLEHR---GFKLNIW---DV---GGQKSLRS---YW
H. sapiens Arl6          -----MGLLDRLS---VLLGLKKKEVHVLCLGLDNSGKTTIINKLKPSN-AQS--QNILPTIGFSIEKFKSS---SLSFTVF---DM---SGQGRYRN---LW
H. sapiens Arl8a         MIALFNKLLDWFK---ALFWK--EEMELTLVGLQYSGKTTFVNVIASG---QFN-EDMIPTVGFNMRKITKG---NVTIKLW---DI---GGQPRFRS---MW
H. sapiens Sar1a         (7)IYNGFSSVLQ----FLGLYKKSGKLVFLGLDNAGKTTLLHMLKDD---RL--GQHVPTLHPTSEELTIA---GMTFTTF---DL---GGHEQARR---VW
A. vaga Arl17a           ---MSSYIRSMLN---KIFGK--PEYRVLIIGLDAAGKTTILYRLKLG---EP--VTTIPTIGFNVESLTYK---SVPLTFW---DI---GGRDKIRP---LF
A. vaga Arl17a2          ---MSSYISSMLN---KIFGK--YEYRALIFGLDASGKTTILYRLKLG---EV--VTTIPTIGFNVESVTYK---SASLTFW---DI---GGRDRIRS---LM
A. vaga Arl17a3          ---MSSYVSSMLN---KIFGK--YEYRALIFGLDASGKTTILYRLKLG---EV--VSTIPTIGFNVESVTYK---SASLTFW---DI---GGRDRIRS---LM
A. vaga Arl17b           ---MASHIRIMLN---KIFGK--PEYRALFIGLDASGKTTILYRFKLG---EV--VTAIPTIGFNVETIEYN---DTKLTIW---DV---GGCDKIRP---LL
C. cinerea Arl17         -----MSIRNLLE---RFFPSNVGEHKVVISGTDWAGKTTLLYYLKRG---EI--IQTIPTIGFNIEHLTLS(5)KLDLELW---DL(4)GGFRMARM---ML
A. fumigatus Arl17       -------MDKLSK---WIFGE--KYHRVLLLGEDSCGKTTFLRRLTFG---EIGEHEPLPTRDFDIETTNYP--ATYKWSIW---EF---RNADRQAS(6)IW
P. blakesleeanus Arl17   MKRFSEIVSQLWS(6)EEQGR--QEIPIRILGARGAGKTTFLYKLYFK(6)ST--FQVFPTESHNVEVIPYR---QFAFQIWEFADI---GAS----------
S. punctatus Arl17       ----METLRTLFG---KWLPA-RADRTLLILGLDACGKTTILYRLVLDP-TEI--ITTIPTIGFNVEKTSIK(4)TVSVSLW---DV---GGCDKIRP---LW
P. polycephalum Arl17a   ----MSFISQLLG---KFFGK--EEYRVLALGLDASGKTSVLYRLKLG---EI--TTTIPTIGFNVETVQIL---GHNVTIW---DV---GGCDKIRP---LY
P. polycephalum Arl17b   -------MNRIM----AMFKK--QACRLLMVGLDAAGKTTILYKLKLG---EI--VTTIPTIGFNVETVEYK---NTAFTVW---DV---GGCDKIRP---LW
P. polycephalum Arl17c   -------MKKFFA---SFFSP-KDETKIILLGLDAAGKTTIIYKMKLG---EV--QTVIPTIGFNLEIVKYK---DLNLNCW---DV---GGCDKIRP---LW
P. polycephalum Arl17d   -------MKRLWA---NFFPP-KDETKIILLGLDAVGKTTIIQKMKLG---DV--ETVIPTIGFSVEIVKYK---NLNLNCW---DV---GGCGQIRP---LW
P. micra Arl17a          ---MGNALGALSS---RMRGK--TEVRVIMNGLDASGKTTALYKLKLG---EV--VTTIPTIGFNVETVQYG---RLNLTLW---DI---GGRDKMRA---LS
P. micra Arl17b          ---MGNELKKARG---ALKGK--NETRILMNGLDAAGKTTILYKLRLG---EV--VCQIPTIGFNVETVSFK---NIQFTVW---DV---GGRDKIRP---LW
K. flaccidum Arl17       ----MSIIGRMIE---KIYGR--PEVRVLLLGLDAAGRTTTLYKLKLG---EI--VTTIPTIGFNVETVTHG---GTSFTMW---DV---GGCDKIRP---LW
C. reinhardtii Arl17a    ----MSRIADFLQ---SFFPS--QEGRVLCLGLDASGRTTWLYAQKLG---EI--VTTIPTIGFNVESVAKD---GVDVTFW---DV---GGCDKIRP---LW
C. reinhardtii Arl17b    -----------MRWRPCWSSK--PEFPILFLGLDAAGKSTVLYGLKLG---EV--VTTVPTIGFNLETIERN---GLKINCW---DV---GGCDKIRP---LW
V. carteri Arl17         ---MSSFVAGLLE---KLFPS--PEGRLLCVGLDACGRTSWLYYKLLQ---EV--VTTIPTIGFNVETIKHA---GCAMTIW---DV---GGCDKIRP---LW
G. wittrockiana Arl17    ---MGNAIKQAFK---KAIGQ--KEVRILMGGLDAAGKTTILYKLKLG---EV--VTTIPTIGFNVETVEYK---NISFTVW---DS---GGRTKMRA---LW
G. avonlea Arl17         ---MGQIPSTALS---SILGK-SEPRKVCLLGLDAAGKTTILYQLKLG---EV--ITTVPTIGFNVETIESG---GLDLTVW---DV---SSRGKLRA---LW
P. bilix Arl17           --------MGILD---VFRRK--KEKRVLLLGLDASGRTSALYRWKLN---DV--VTTVPTIGFNVETLKAG--PNTILTAW---DV---GGCSKIRP---LW
R. erinaceoides Arl17    -------------------------------------RTTTLYTWKLG---EV--VTTIPTIGFNVETIQHN---GVSLTIW---DV---GGCDKIRP---LW
C. velia Arl17a          (5)FLDNLKGVLG----LGGA--KQSRGLFLGLDASGRTTLLYRLVLG---EC--VPTIPTIGFNVETVTHR---NTNFTLW---DV---GGCDKIRP---LW
C. velia Arl17b          MG--------------QLGSREKKEIRTALVGLENAGKTTALYRLKFA---QE--SPAVPTIGFCVETFLYK---ELNLVVM---DMQKGRGMGKLRP---LW
S. minutum Arl17         ---MGQSIQILKK---TLAGA-NVNKRVLLVGPNAAGKTCLLYGLKLGF-DQN--LTTIPTVGFNVETFTHG---RKDFTVW---DL---GLRSKMRP---LV
R. filosa Arl17          ----MSSITETLN---RLFKRKDREIRSLMLGLDASGRTTALYKLKLG---ET--VTTIPTIGFNVETVEYK---NTTFTFW---DV---GGCDKIRP---LW
Consensus/80%            .......h..hb.....bbs....ph+hlbhGLDstG+TThLYbbpbt...ch..hsslPTlGFNlEplpb....shpbshW...Dl...GGps+bRs...lb
```

```
                                                                          G4 box
                                                                           ****
H. sapiens Arf1              RHYFQNTQGLIFVVDSNDRERV----NEAREELMR-MLAE--DEL---RDAVLL-VFANKQDLPNAM--NAAEITDK----LGLHSL-----RHR-NWYIQ--
H. sapiens Arf6              RHYYTGTQGLIFVVDCADRDRI----DEARQELHR-IIND--REM---RDAIIL-IFANKQDLPDAM--KPHEIQEK----LGLTRI-----RDR-NWYVQ--
H. sapiens Arl1              RCYYSNTDAVIYVVDSCDRDRI----GISKSELVA-MLEE--EEL---RKAILV-VFANKQDMEQAM--TSSEMANS----LGLPAL-----KDR-KWQIF--
H. sapiens Arl2              RNYFESTDGLIWVVDSADRQRM----QDCQRELQS-LLVE--ERL---AGATLL-IFANKQDLPGAL--SSNAIREV----LELDSI-----RSH-HWCIQ--
H. sapiens Arl6              EHYYKEGQAIIFVIDSSDRLRM----VVAKEELDT-LLNH--PDIK-HRRIPIL-FFANKMDLRDAV--TSVKVSQL----LCLENI-----KDK-PWHIC--
H. sapiens Arl8a             ERYCRGVSAIVYMVDAADQEKI----EASKNELHN-LLDK--PQL---QGIPVL-VLGNKRDLPGAL--DEKELIEK----MNLSAI-----QDR-EICCY--
H. sapiens Sar1a             KNYLPAINGIVFLVDCADHSRL----VESKVELNA-LMTD--ETI---SNVPIL-ILGNKIDRTDAI--SEEKLREI(12)VTLKEL-----NAR-PMEVF--
A. vaga Arl17a               RHYFHNTQALMFIVDSNDRERL----PEATDTLWR-LLCE--DEL---REIPVL-IYRNKMDLEHTL--SQEDIIEQ----MRLNDI-----RNR-PWHIQ--
A. vaga Arl17a2              RHYFQNTQVLIFIVDANDRERL----PEATDALWR-LFDE--DEL---REIPLL-IYMNKIDLEHSL--LREDLIEQ----MRLNHI-----RNR-PWYLQ--
A. vaga Arl17a3              RHYFQNTQALIFIVDANDRERL----PEATDALWR-LFDE--DEL---REIPLL-IYMNKIDLEHSL--LREDFIEQ----MRLNDI-----RNR-PWYLQ--
A. vaga Arl17b               RHYFQNTQAMIFVIDTNDRERL----HEAKDELSR-LVSE--EEM---YGVPIL-FYLNKTDLIHAL--PINEIVEQ----MRMSHM-----RNR-SYHVQ--
C. cinerea Arl17             RHIAYTAKAIIWVVDSTDTPEMM---AESAETLLI-ALEDA-EEM(6)RHLPVL-ILANKSDLPNAM--PLDDIRKV----FRHAT------AGR-IASIY(4
A. fumigatus Arl17           RELAPHT-LVLWLHDCSTEDDWP---PWKFRSLLE-QMVE--------RGCRYIWVGLNKQDSPDVSEESVQEARRK----YEEEFA---KYKDDLSWKVL-T
P. blakesleeanus Arl17       LNYIKDTRVIIYMVDAVEQSKP-VVASKARENMSW-ILKTFEEEL---RDAIVI-TVVNKIESEGAV--DIQDLGQQ(4)PLLTKGL-----RNH-QWRIF--
S. punctatus Arl17           RHYFSGLSGLMYIVDSNDRERF----PEAVEEFRA-MLTEI-AAL(5)QRIPVI-VIANKQDLPNAM--GTLEIREK----FAPTV------GNR-AFAVY--
P. polycephalum Arl17a       RHYFANTQGLIWMVDSNDHQRI----EETSEELAK-VLRD--DEM---RDVALL-VFANKQDLPGCM--SVSEITDK----LGLHAL-----RNR-KWFIK--
P. polycephalum Arl17b       RHYFQNTQGLIFVIDSNDKDRL----QETTEELER-FMKE--DEL---KDAVLL-LMANKQDLPNSL--KPDEIAKK----MDLAKT----CAGR-KWRIQ--
P. polycephalum Arl17c       RHYFDGMHALIFVVDSNDRERI----DEAKKEMDV-MLRELEDNP---HPIPVL-VFANKQDLPNCM--SPEEIADK----MGLKSI-----KNS---HIT--
P. polycephalum Arl17d       RYYFDGLHALIFVIDSNDPTGL----PSAKSELDF-MLREMAQSP---CPVPIL-VFANKQDLPNCM--SPEELADK----IDIKFV-----NNA---HIR--
P. micra Arl17a              RHYYQNTQAVIWVIDSNDRERL----PDAMDEMHR-AWNE--HEL---EAAVWL-VLCNKQDLPNAM--SRDEIVAS----LPTSMS-----RSP-RLRAV--
P. micra Arl17b              RHYYQGTNAVIWVIDSNDHDRM----DETLEEMVM-AAKE--DDLS-GSDIVWL-ILCNKQDLPGAL--SPTLIEAK----LPKEIK-----NRR-ACRVV--
K. flaccidum Arl17           RHYFQNTQALIWFAHSSDRDRV----DEAREEIQR-LLSE--EVL---QDVPFL-VWATKQDLPNVM--TPEEVSSK----LALQNL------DR-PWACI--
C. reinhardtii Arl17a        RHYFQNTDGVMLFVDSNDRERI----QEVRDEFNR-LLSE--EEL---RDAFCL-VIASKQDLPNVM--SLAEIRDA----LDLPRL----MKDR-HWVLM--
C. reinhardtii Arl17b        RHFTEGCVAVVFVVDSNDRERL----AAARQELQD-VLRDV------ASGVPVL-VLANKQDLPNAL--PAAEVARL----MWLAPP(66)LMSRHAVHVQ--
V. carteri Arl17             RHYFQNTHGVVAFVDSNDRDRL----DEVRKEVLA-LMDA--PEL---RDAFCL-VIANKQDLPCAM--PPREVSEK----LQLPRA----MGSR-LWTCL--
G. wittrockiana Arl17        RPYYANTNGVIWVVDAQDKDRI----SETAEELHH-LFRE--DEL---RDCVFL-ILINKGDLPNAM--NLREAEEL----LNLSDE-----RKRHKIEVR--
G. avonlea Arl17             RHYYLGSAASIFVIDSTDRDRL----EEALEELVI-VMEH--------ADGVFL-LLANKQDLSGAM--SPEEIAKK----VSEVAP-----PNT-LWGLF--
P. bilix Arl17               RHYFGNTSAVFLFWDI-ERERQ(16)ETTGQHLRS-LLRE--ADL---EHVPIL-LVVSKADL--AI--SEEEIGS---------------IRAK-FYAIL--
R. erinaceoides Arl17        RHYFQNTQLCVFHVDSNDRDRM----PEAVLGLQR-VLGE--DEL---KGAPIL-VLANKQDLPNAM--PAHEIKSL----FLASIH----PNDR-EFEVI--
C. velia Arl17a              RHYFQGTCVMFFFVDSADRERL----QDAFKEITKYVLCE--DAL---KGVPLV-LVMTKQDLPGCM--PPSEILDK----LQVEKN----LRGR-WWKHV--
C. velia Arl17b              KHYFEGLGAIIFMVDASDLEAI----GDARTALWQ-LVDE--LEG---RDVIVA-VLANKQDKAEAL--GPEEVALQ----LRFTEL----PQAR--KRAF--
S. minutum Arl17             RHYLPGTDMVIFMISGLSGVHS-ELDMEDAEDLLN-LVRD--GLMENQAAVPFL-FFVNFMHEPKHM--TMTEIIER----LQLNRL----ARQT-RVHLQ--
R. filosa Arl17              RHYYQNTGVIWYFVDSRDRERI(11)TSSVPGLND-SLTE---EL---RDAVFC-VLCNKQDLPNAM--SVNEISER----LQLHKV----LKGR-DWNIF--
Consensus/80%                RcYb.sspsllahlDtsDcp+b.....ps.ppl...hbp-....b...p.h.hl.lbhNKbDL..sb..s.p-l.pb....b.b.........s+..b.lb..
```

```
                                   G5 box
                                    ***                      >>>
H. sapiens Arf1                   -ATCAT---SGD--GLYEGLDWLSNQLRNQK----
H. sapiens Arf6                   -PSCAT---SGD--GLYEGLTWLTSNYKS------
H. sapiens Arl1                   -KTSAT---KGT--GLDEAMEWLVETLKSRQ----
H. sapiens Arl2                   -GCSAV---TGE--NLLPGIDWLLDDISSRIFTAD
H. sapiens Arl6                   -ASDAI---KGE--GLQEGVDWLQDQIQTVK---T
H. sapiens Arl8a                  -SISCK---EKD--NIDITLQWLIQHSKSRR---S
H. sapiens Sar1a                  -MCSVL---KRQ--GYGEGFRWLSQYID-------
A. vaga Arl17a                    -SCSAT---KGD--GLYEGLDWISRAVQSPS----
A. vaga Arl17a2                   -PCSAT---RGD--GLYEGLDWMLRAIRSPS----
A. vaga Arl17a3                   -PCSAT---RGD--GLYEGLDWMLRAIRSPS----
A. vaga Arl17b                    -PCSAI---NGD--GLYEGLDWLISVLKSSS----
C. cinerea Arl17                  )PVVTK---SVT--GLQEAIDWLALALDIAS----
A. fumigatus Arl17                HKLSAK---TGV--GVSEVLKDIYQAVKRAN----
P. blakesleeanus Arl17            -ECDAA---TEK--GFEKVLDYLSQKIEMKD----
S. punctatus Arl17                -PSRAL(4)SQT--GLPEAFEWLTEQIVTPQ----
P. polycephalum Arl17a            -GVCAT---RGDISEVYDGMEWLVNAIDSTPFTHT
P. polycephalum Arl17b            -GCCAT---SGD--GLYEGLDWLSRAIAENP-PPA
P. polycephalum Arl17c            -PSCAL---SGD--GLHKGFDWLSDAVAKYK----
P. polycephalum Arl17d            -GSIAT---TGE--GLYEGFEWLSDAVAKYK----
P. micra Arl17a                   -GCCAK---DGD--GLYDGLAWLQYAMQHQA----
P. micra Arl17b                   -GTSAI---RGD--GLYEGLDWLSFALDRNN----
K. flaccidum Arl17                -GCSAF---TGE--GLYEGLQWIQDALRAKR----
C. reinhardtii Arl17a             -PTRLP--VDKQ--ELDGHMDWLVASIRAERNQVA
C. reinhardtii Arl17b             -PTCAL---RLF--GVLEGFDWLASALDHARQQDS
V. carteri Arl17                  -PGQLG---SMQ--LTNAHFDWLASALRHTL----
G. wittrockiana Arl17             -TTCAT---SGE--GLLEGLQWLADALSEKA----
G. avonlea Arl17                  -PCSAT---KAE--SLHDALSWLGKVLQGKD----
P. bilix Arl17                    ---------EDE-C----GDDYVADHCRISM----
R. erinaceoides Arl17             -PTCAT---SRA--GLAEALDWIVDTIKHTK----
C. velia Arl17a                   -AVATP---EGI--GIEEVKEAMWEANEAFL----
C. velia Arl17b                   -GTAAADDRGSA--ELFEALDWILEVHRETK----
S. minutum Arl17                  -PCCAK---TGD--GVMQGLDFLASYSGESV----
R. filosa Arl17                   -AVSAI---KGD--GLYEALEWTQEAYFDPK----
Consensus/80%                     .ssss....p.p..sl.ctbcWl.p.hp.......
```

**Supplementary figure 10. Phylogenetic analysis of the novel C-terminal domain of Arl17 proteins**. The presented tree is a result of a ML analysis of all individual copies of the novel C-terminal domain of 42 Arl17 proteins. The sequences were aligned by the on-line MAFFT, the alignment was trimmed manually, and the tree was inferred using IQ-TREE with LG+G4 model (the model selected by the program itself) with the ultrafast bootstrap algorithm and the SH-aLRT test (both 10000 replicates). Dots at branches represent bootstrap values as indicated in the graphical legend (bottom left). The novel domain (ND) copies are numbered according to their position in the Arl17 protein and highlighted in different colours (first copy – blue; second copy – violet; third copy – black). Note that individual Arl17 proteins may have from one to three copies of the domain.

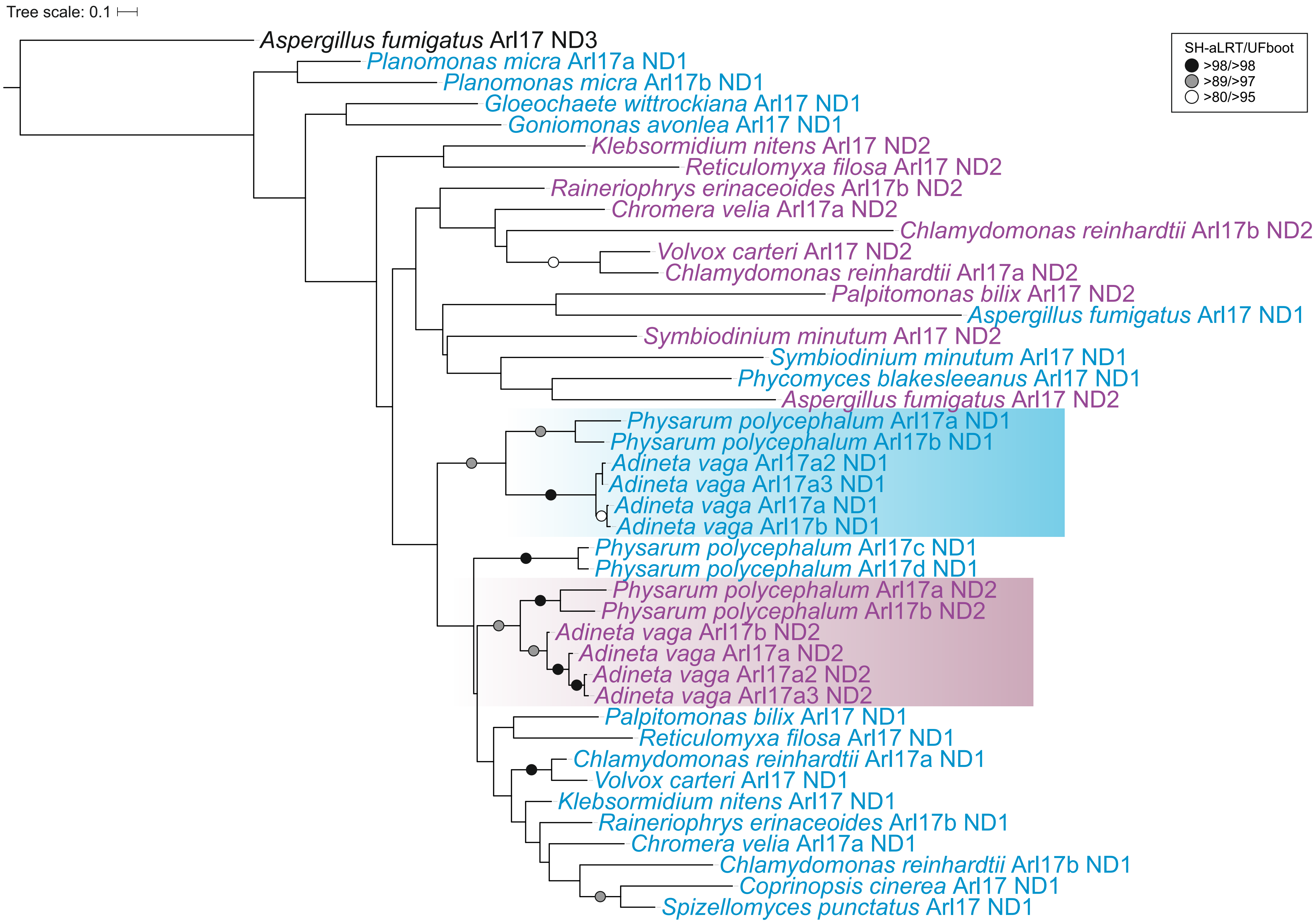

**Supplementary figure 11. Position and phase of introns in holozoan Arf1, 4, 6, Arl4, 19, 20 and TRIM23 genes mapped onto a multiple alignment of protein sequences**. The intron positions are marked by highlighting the amino acid residues encoded by a codon located immediately upstream of the intron (phase 0; in red) or interrupted by the intron at the second or third position (phases 2 and 3; in green or blue, respectively). The identity of the sequences included is provided in Supplementary table 1. Note that genes with the coding sequence contained in a single exon and sequences represented only by transcriptomic data are not included in the alignment as indicated in Supplementary table 1, column J. Note that for simplicity the alignment regions corresponding to the N- and C-termini of the protein sequences were trimmed. The sequence on a white background is a divergent *C. intestinalis* gene annotated as Arf1/6. A large insert in the Arl4 sequence from *D. melanogaster* was replaced by three red dots for simplicity. Note that only the GTPase domain of TRIM23 sequences is included in the analysis.

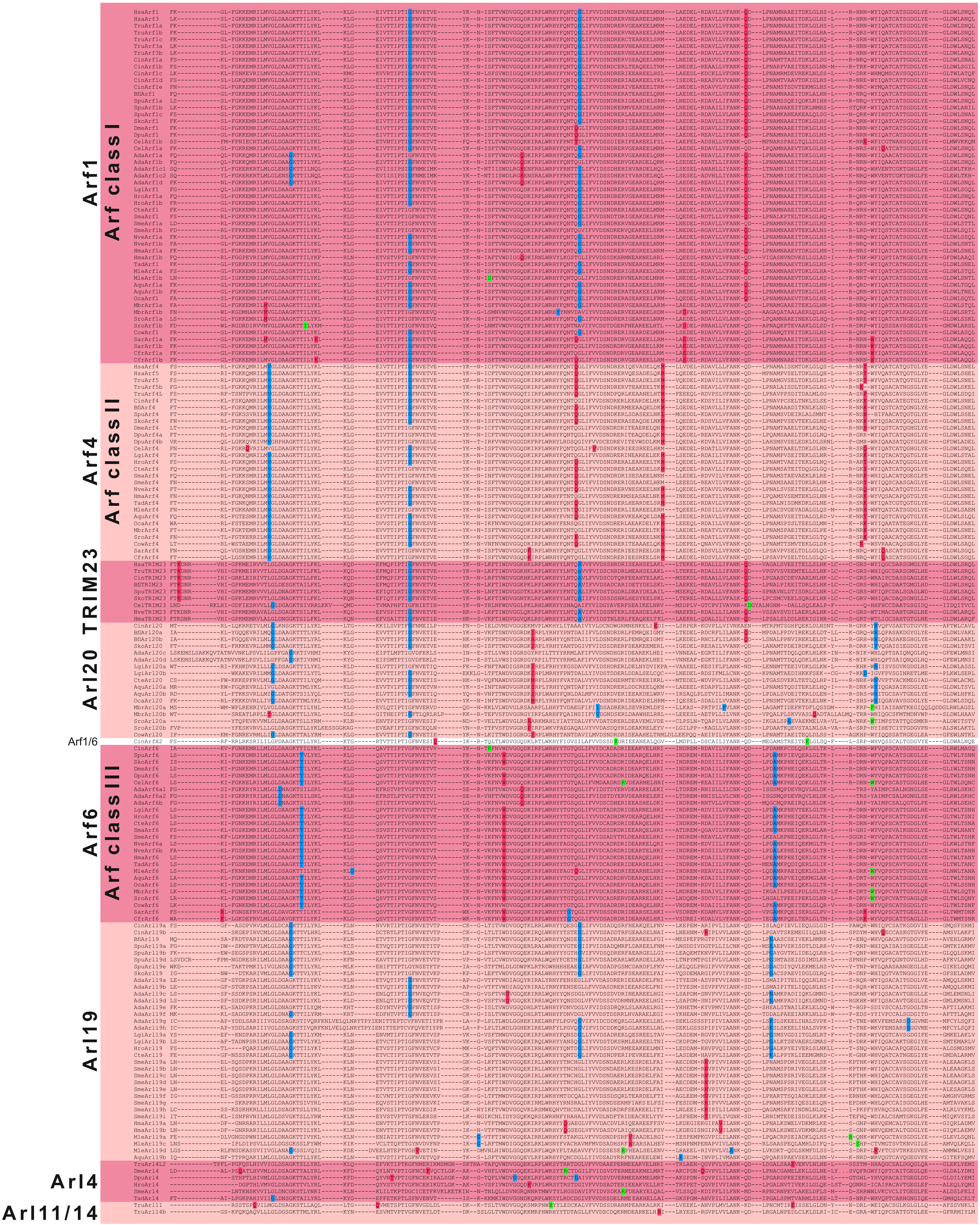

**Supplementary figure 12. Phylogenetic analysis of Arl20 and Arl10**. The trees shown are results of ML analyses of (A) Arl20 sequences and a subset of the holozoan "scrollsawed" dataset restricted to TRIM23, Arl1, 4, 5, 19 and Arf1, 4, 6 sequences (152 sequences in total), and (B) of Arl10 sequences with the complete holozoan "scrollsawed" dataset (328 sequences in total). The trees were inferred using IQ-TREE with LG+I+G4 model (the model selected by the program itself) with the ultrafast bootstrap algorithm and the SH-aLRT test (both 10000 replicates). Dots at branches represent bootstrap values as indicated in the graphical legend (top right).

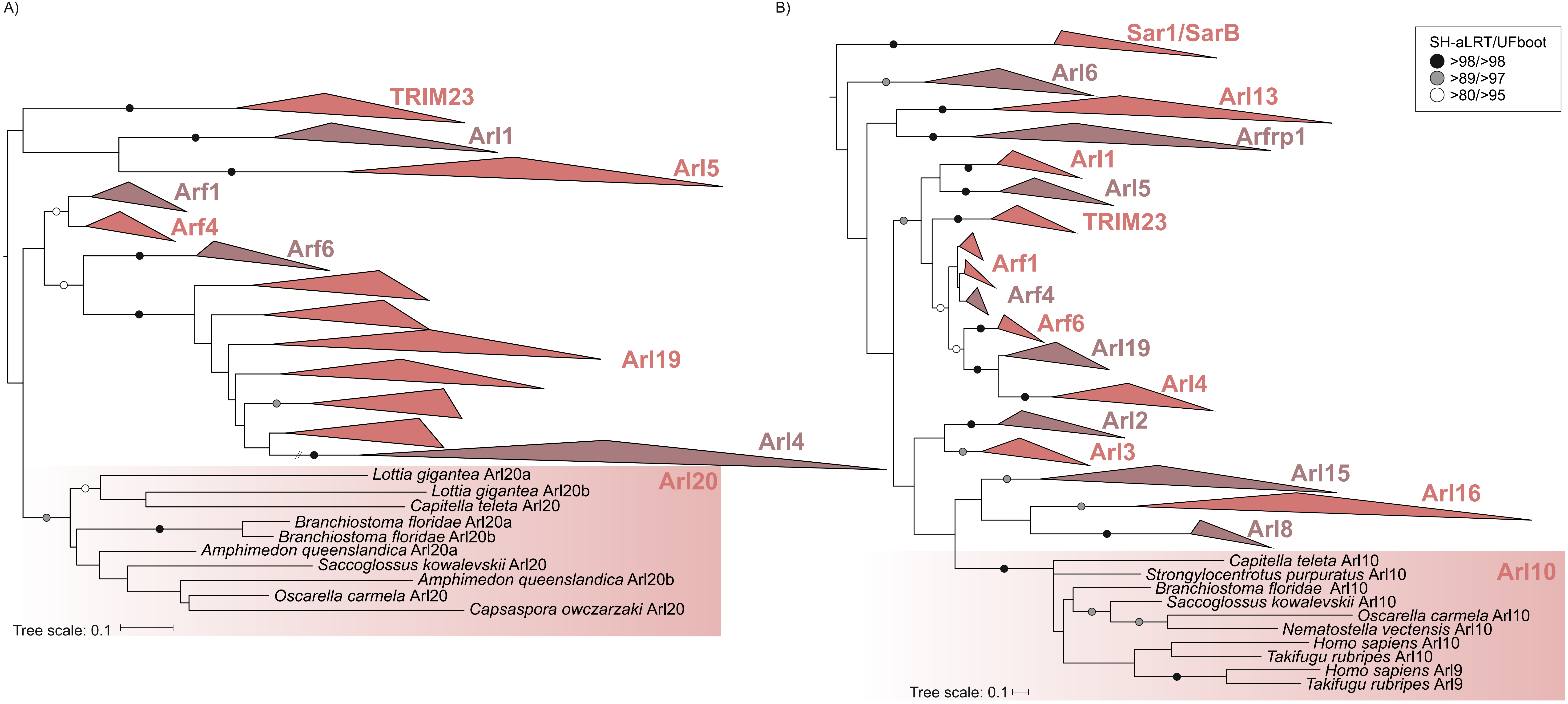

**Supplementary figure 13. Phylogenetic analysis of Arl11/14**. The tree shown is a result of a ML analysis of vertebrate Arl11 and Arl14 sequences and a subset of the holozoan "scrollsawed" dataset restricted to TRIM23, Arl1, 4, 5, 19 and Arf1, 4, 6 sequences (147 sequences in total). The alignment was trimmed manually and the tree was inferred using IQ-TREE with LG+I+G4 model (the model selected by the program itself) with the ultrafast bootstrap algorithm and the SH-aLRT test (both 10000 replicates). Dots at branches represent bootstrap values as indicated in the graphical legend (top right).

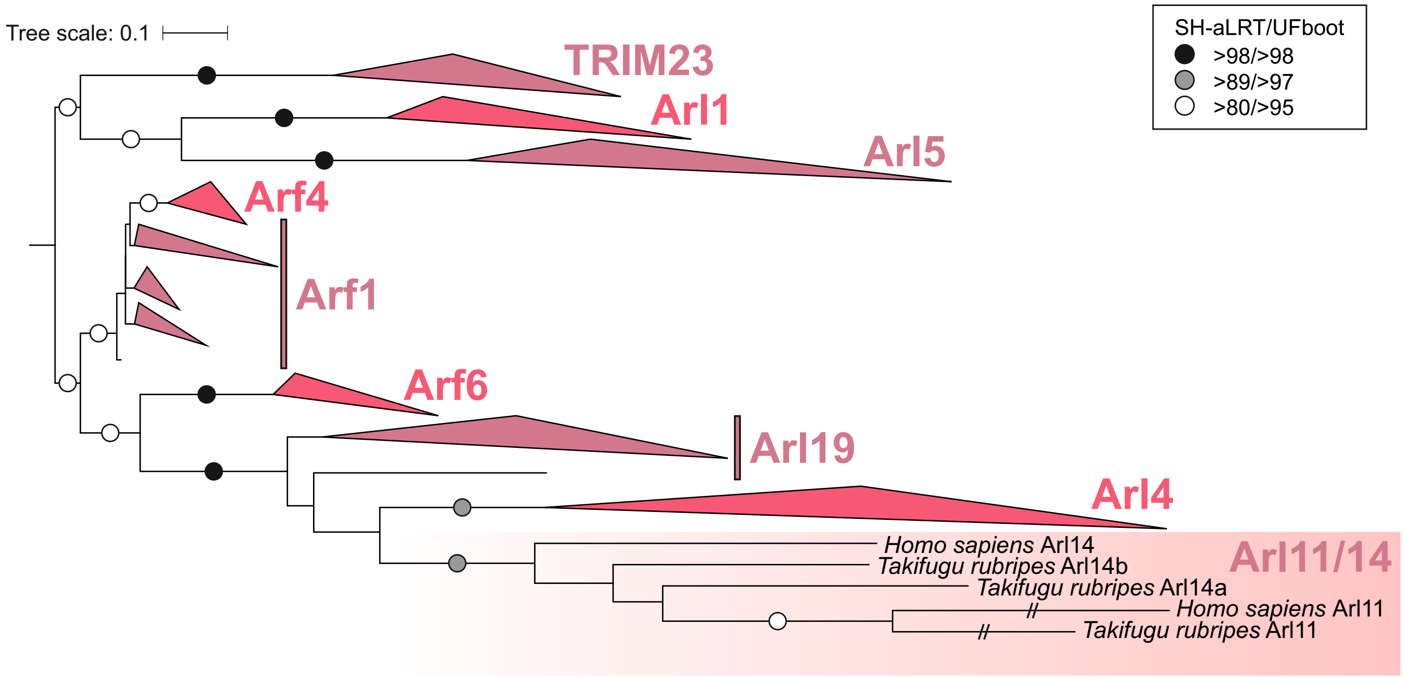

**Supplementary figure 14. Phylogenetic analysis of the glaucophyte-specific Arl13L group**. The tree shown is a result of a ML analysis of all glaucophyte Arl13L sequences and the reduced "scrollsawed" dataset (354 sequences in total). The tree was inferred using IQ-TREE with LG+I+G4 model (the model selected by the program itself) with the ultrafast bootstrap algorithm and the SH-aLRT test (both 10000 replicates). Dots at branches represent bootstrap values as indicated in the graphical legend (top right). Typical Arl13 proteins from *C. paradoxa* and *G. wittrockiana* are highlighted in bold.

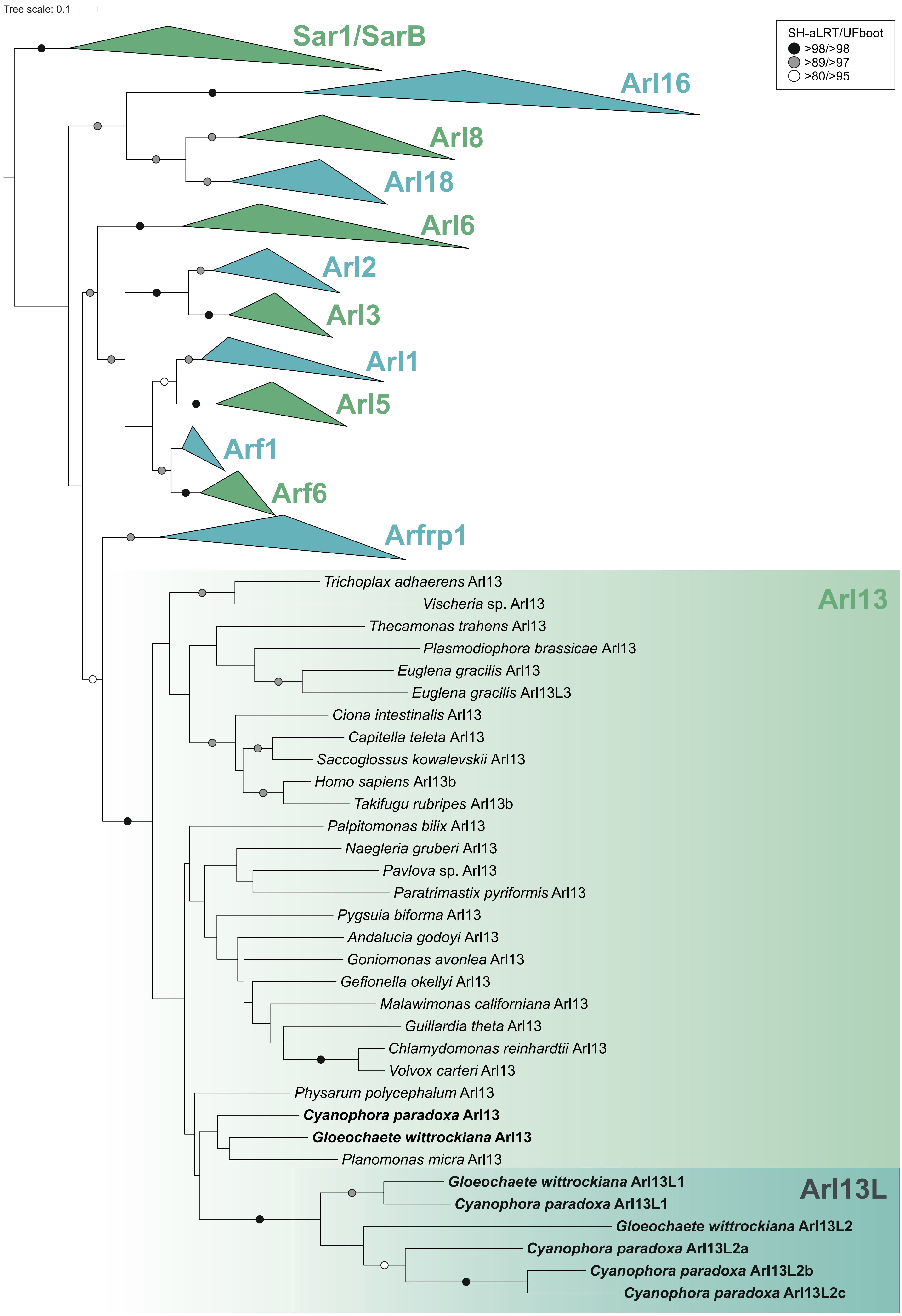

**Supplementary figure 15. Phylogenetic analysis of ArfB.** The tree shown is a result of a ML analysis of all ArfB sequences and a subset of the reduced "scrollsawed" dataset restricted to Arl1, Arl5, Arf1 and Arf6 sequences (117 sequences in total). The alignment was trimmed manually and the tree was inferred using IQ-TREE with LG+I+G4 model (the model selected by the program itself) with the ultrafast bootstrap algorithm and the SH-aLRT test (both 10000 replicates). Dots at branches represent bootstrap values as indicated in the graphical legend (bottom left).

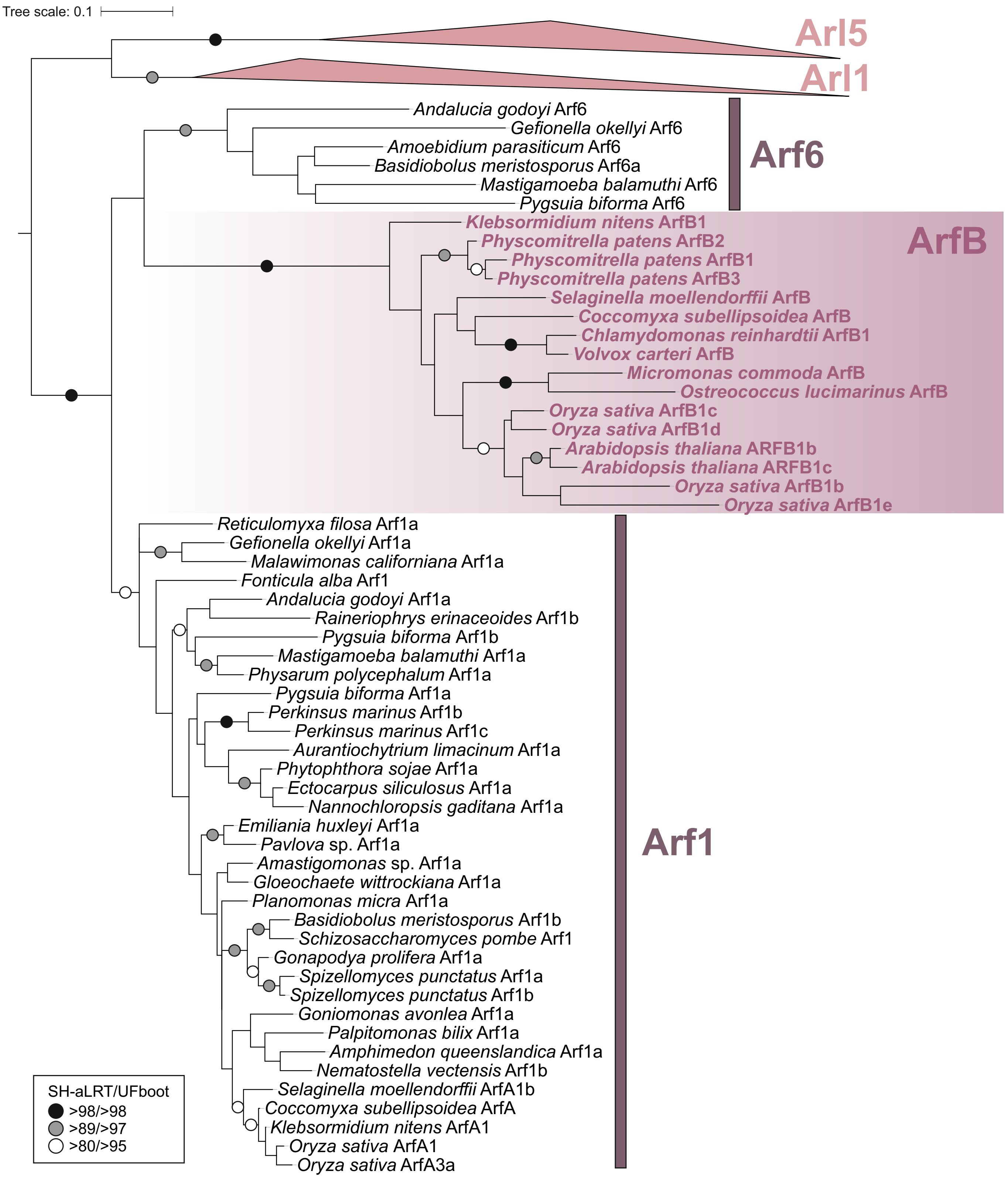

**Supplementary figure 16. Annotated multiple sequence alignment of the N-terminal region of Arl15 proteins.** Amino acid residues of special interest and residues predicted to carry a specific lipidic modification are highlighted, as indicated in the graphical legend below the alignment. The unique absence of cysteine residue(s) in the N-terminal extension of the *S. rosetta* Arl15 protein suggests an alternative, yet unknown, means of membrane association or a lineage-specific functional shift of the protein.

```
                                                               <<<<<<<<<<<<<<<<<<<<<<< GTPase domain >>>>>>>>>>>>
Homo sapiens Arl15               --MSDLRITEAFLY-------MDY----LCFRALCCKGPPPARPEYDLVCIGLTGSGKTSLLSKL-CSES-PDNVVSTTGFSIKAVPFQNAILNVKE
Takifugu rubripes Arl15          ---------------------MDY----LCFRTLCCKGPPPPPRPEYDVVCIGLTGAGKTSLLSRL-CSEA-SDGIVPTTGFSIKAVPFPNAILNVKE
Ciona intestinalis Arl15         -------MAEHGSVCCGACRFLCC----SMFHCICCKDSPPELPVYNVICIGITNSGKSTLLKVL-THEN-SDTVVPTVGFNVKDIRLPNAVLKVKE
Branchiostoma floridae Arl15     ------MCSERCQLAIGACRIICF----NLCRGLCCCTPPTPRPEYHVVCLGMSGAGKTTLLTLLQNGET-TDDMEPTSGFNIKAIQFEEAILNVKE
Saccoglossus kowalevskii Arl15   ----MPSIGEIFGLCCGACRIGCY----TLYVKLCCKSAPSQRQEYQLICLGLSKAGKTTLLTML-CGEK-ADDIQPTQGFNIKAIPFKEAILNVKE
Lottia gigantea Arl15            -------MCENIRLMCVVCRIGVY----SLCRRMCCMGPPSIRPNFTVLCLGLTKAGKSSLLAVL-SGDD-IHKIEPTIGFSIKAIMFDNCILDVKE
Capitella teleta Arl15           (4)DAAHRFESLEVKFEIC---CF(15)QIYKRVFCIKEPAPRPVFKPICIGLDDAGKSTLLSSL-AGES-TDDIQPTKGFVIKAIQFDDCILNVKE
Schmidtea mediterannea Arl15     -------MGQNFSFAGGICRIFCY----KTYKTICCVPDIKYLPVCNALCVGLANVGKSRMLAAL-NKEN--LNTEPTIGFNVKAIKLDNCILSIKE
Nematostella vectensis Arl15     ----MSSKCECCGLLCAVLRIGCF----RCCQKLRGKKPDAPRPEYSVLMIGLSGAGKSVMLAKL-DSEP-TDDIKPTQGFSVKPVQLPTAILNVKE
Hydra magnipapillata Arl15       ----MTGFCLNCQFCFAASRIFCY----EFCC-CCIHKKKDLKRNLNVLVLGLESCGKSYLLASL-CGES-TKDLTPTNGFSIKDVSFKNYTMHMKE
Trichoplax adhaerens Arl15       ----MTTVRECVLLAFAFCRIGLY----TCYRRIFYGPPPPPRPGFSAICIGITGAGKSKVLAVL-CSEK--DEIIPTQGFSIKDVPFENAILHVKE
Monosiga brevicollis Arl15       -------MGNALALCWGVVRIACY----QVCK-ICLPAEPPPRPIYDVLCLGMAGTGKSALLAAL-NHEP-YTDLEPTKGFAIKPVLTSKVQFNVKE
Salpingoeca rosetta Arl15        ---MLDAIWQRLSFAWGVIRIASY----SAWRFFFPADEQQQRPTVSALVLGLQGVGKTTMLTRL-ANETPQQDIPPTKGFAIKAIQTPACIINAME
Consensus/80%                    .................................................................................................
```

C - cysteine residue predicted to be S-palmitoylated
C - cysteine residue
G - glycine residue predicted to be N-myristoylated

**Supplementary figure 17. Annotated sequence alignments of taxon-specific Arf1-like proteins carrying a predicted N-terminal transmembrane domain.** The figure includes two examples of such proteins, representing taxon-specific paralogs from chlorarachniophytes (**A**) and the haptophyte class Pavlovophyceae (**B**). The approximate position of the predicted transmembrane domain is indicated above each alignment. Arf1a from *Bigelowiella natans* (**A**) and Arf1 from *Pavlova pinguis* (**B**) represent a conventional Arf protein without an N- terminal transmembrane domain. Accession numbers of the sequences coming from organisms that are not part of the core set of taxa analysed in this study are provided in Supplementary table 6.

**A)**

```
                                                                     <<<<<<<<<  TM  >>>>>>>>>                                <<<<<<<<<<<<<<<<<<<<<<<<
Bigelowiella natans Arf1TM            -------MG---------------ADRPRIKLRLAGVLLVAVACAA----LRSLDMRRDAPRRRPVLQSQAVKGLAITFRNMWREFFARRTPTNVVM
Bigelowiella longifila Arf1TM         -------AG------------------RRLKLRLAGVLLVVVVCTA----LRSLDMRRHSPRHRPVLRSQAMKGLAITLKNMWRDFFVRRIPTNVVM
Chlorarachnion reptans Arf1TM         M(18)VQEG(8)MTRLTIPIRAQRVKLLRI--YLLFAVLVALVHVG(12)KLAIRASTSPSSAPAVASLQAIRGLTASLGSMWRSVFSRRVRTNVVM
Lotharella globosa Arf1TM             M(1)-VRFG(5)RTR----RRLRHRTRRRL--WLGVAAAVAI--------AVFPQAQLVEEGRRGRLNSTIVRGVAEAVGGMWRSLFDRRIRTNVVM
Lotharella oceanica Arf1TM            M(6)WSRIG---------------GARSRL--WMGLAAVLAI--------AVFPQARIVEEGRRGRLNSTIVRGVAEAVGGIWRTIFDRRVRTNVVM
Lotharella amoebiformis Arf1TM        -------MG-----------LLSHRRGRPLSLLLAAVVLAST--------PNCPSKSPYRSDRSNRLQSQMLRGVAEVMGSMWKRWFERRILTNVVM
Gymnochlora sp. Arf1TM                -------MA---------------SRFRRVKNLIVVVSIVAL--------LVHASFMTRMSKSKSRLRSQALRGFTEAVGNLWTRVFSRRVLTNVVM
Norrisiella sphaerica Arf1TM          -------MA---------PHLASQRLCRRL--AFRFCLLGCICMAMMI--TCSGKQNTYGSHSRRDLRSEAIRGLAKSVRDLWTQTFWRKIPTNVVM
Partenskyella glossopodia Arf1TM      ------------MFR--PPHVASRG-------VLRALPVVTV--------LVLGLALSATPPPRQSLRSEAIRGFTQVVSGFWKSIFCRKTPTNVVM
Bigelowiella natans Arf1a             --------------------------------------------------------------------------MGMTFSNFWKRMFGNK-EMRILM
Consensus/80%                         ........t....................l...b.hs.lssl............p........p..LpSphl+Ghsbsh.sbW+phFsR+h.TNVVM

                                      <<<<<<<<<<<<<<<<<<<<<<<<<<<<<<<<<<<<<<<  GTPase domain  >>>>>>>>>>>>>>>>>>>>>>>>>>>>>>>>>>>>>>>>
Bigelowiella natans Arf1TM            LGLDSAGKTAVLYKLRLGKTVATIPTMGFNTEPIPYKDFEFRIWDVGGQNRIRTLWQHHLNGAKVVIFVVDSADKERLPQAAEELHRLMTDPRLRGA
Bigelowiella longifila Arf1TM         LGLDSAGKTAVLYKLRLGKTVATIPTMGFNTEPIPYKDFEFRIWDVGGQNRIRTLWPHHLNGAKVVIFVVDSADKERLPQAAEELHKLMGDGRLRNA
Chlorarachnion reptans Arf1TM         LGLDSAGKTAVLYKLRLGQAVQTIPTMGFNTESIPYKKFEFRIWDVGGQDRIRSLWQHHLDGAKVVIFVVDAADRTRIPKAANELHKLMQDPRLSDA
Lotharella globosa Arf1TM             LGLDSAGKTAVLYKLRLGTNVQTIPTMGFNTEAIPYKNFEFRIWDVGGQDRIRALWQHHLDGARVVIFVVDSADRARVPKAASELHRLLKDPRLRYA
Lotharella oceanica Arf1TM            LGLDSAGKTAVLYKLRLGTNVQTIPTMGFNTEAIPYKNFEFRIWDVGGQDRIRALWQHHLDGARVVIFVVDSADRARVPKAASELHRLLKDPRLRYA
Lotharella amoebiformis Arf1TM        LGLDAAGKTAVLYKLRLGTTPQTIPTMGFNTEAIAYKNYEFRIWDVGGQDRIRTLWDYHLDGARVVIFVVDSADRARLPKARMELHRLMKDERLKQA
Gymnochlora sp. Arf1TM                LGLDAAGKTAVLYKLRLGTTVQTIPTMGFNTEAISYKNYEFRIWDVGGQDRIRSLWQYHLEGARVMIFVVDSADRPRLQQAKSELQKLLTDPRLNGA
Norrisiella sphaerica Arf1TM          LGLDAAGKTAVLYKLRLGKAVQTIPTMGFNTETIPYKNFEFRIWDVGGQDRIRALWQHHLDGARVVIFVVDSADKDRLPQAATELHKLMRDPRLRDA
Partenskyella glossopodia Arf1TM      LGLDAAGKTAVLYKLRLGQAVQTIPTLGFNAEAIPYKNFEFRVWDVGGQDKIRTLWQYHLSGAKVVIFVVDSADKSRLPVAAEELHKLMRDPRLSNA
Bigelowiella natans Arf1a             VGLDAAGKTTVLYKLKLGEVVTTIPTIGFNVETVEYKNINFTVWDVGGQDKIRPLWRHYYQNTQGIIFVVDSNDRERLDMAKEELDKMLNEDELRDA
Consensus/80%                         LGLDtAGKTAVLYKLRLGpsVpTIPTMGFNTEsIsYKsaEFRIWDVGGQDRIRsLWpaHLsGA+VVIFVVDSAD+.RlPpA.pELH+LbpDsRLp.A
```

```
                                      >>>>>>>>>>>>>>>>>>>>>>>>>>>>>>>>>>>>>>>>>>>>>>>>>>>>>>>>>>>
Bigelowiella natans Arf1TM            HLLVFANKQDQEHAVSAQEMVNELRLGHVTNRSWFLQGSNARTGEGLYEGLDWLVSELRLEQEQKRLEQS(8)GGN-GGGRM---
Bigelowiella longifila Arf1TM         YLLVYANKQDQDHAVTAEEMVKELRLRRVTSRPWFLQGSNARTGEGLYEGLDWLVSELRLEQEQKRLEQS(8)DGNDGTGRI---
Chlorarachnion reptans Arf1TM         HLLVYANKQDQDGAMTAKEMVEALHLNMITNRSWFLQGSCARTGEGLYEGLDWVCSELAVEQELQERQS------------LRQY
Lotharella globosa Arf1TM             HLLVYANKQDQRSALTAKEMAEALSLGNITNRSWYLQGSCARTGEGLYEGLDWLVSELDAEPDGP--------------------
Lotharella oceanica Arf1TM            HLLVYANKQDQRSALTAKEMAEALSLNNITNRSWYLQGSCARTGEGLYEGLDWLVSELDAEPVEP--------------------
Lotharella amoebiformis Arf1TM        HLLVYANKQDQANAISAKNLATQLNLEAIRNRSWYLQGSCARTGEGLYEGLDWLVSELRASDEPT-------------------
Gymnochlora sp. Arf1TM                HLLVYANKQDQPNALSAKEIAVGLGLEDLKNRNWYLQGSCAQTGEGLYEGLDWLVSELRAAAEAAAQV-----------------
Norrisiella sphaerica Arf1TM          YLLVYANKQDQRHAITAREMVERLDLSSISDRSWFLQGSNARTGEGLYEGLDWLVSELLIEKDLAEAAA---AATQGSSRMRFF
Partenskyella glossopodia Arf1TM      HLLVYANKQDQNQALNVRQMVDALRLRAITNRSWYLQGSNARTGEGLYEGLDWVVNELDAERAALSSR-----------------
Bigelowiella natans Arf1a             VVLVFANKQDLPKALKVKNITEALGLDKM-RRQWFVQGTCATSGDGLYEGLDWLSSTLSKKK----------------------
Consensus/80%                         aLLVYANKQDQ.pAl*A+pbsp.LpLp.lpsRsWaLQGSsARTGEGLYEGLDWLVSELphc.......................
```

**B)**

```
                                         <<<<<<<  TM  >>>>>>>          <<<<<<<<<<<<<<<<<<<<<<<<<<<<<<<<<<<<<<<<<<<<<<<<<<<<<<<<<<<<
Pavlova pinguis Arf1TM               M-VARRALALALCVFACLANVAEGGAAFSKILGSLILNRNVRILMLGLDGSGKTTVLYRLKLNKYITTAPTIGFNAESLKYRNLEMTVWDVGGQE
Pavlova gyrans Arf1TM                MGAVRVCLCVALC-LACFAGSAEGGSAFSKILGSLLLNRNVRILMLGLDGSGKTTVLYRLKLNKYITTAPTIGFNAESLKYRNLEMTVWDVGGQE
Pavlovales sp. CCMP2436 Arf1TM       MAVSRVLRFALLAALACTFSVIEGGAAFSKILGSLLLNRHVRILMLGLDGAGKTTVLYRLKLNKYITTAPTIGFNAESLKYRNLEMTVWDVGGQE
Diacronema lutheri Arf1TM            MALARIGRLLALVLVACAVG-VEGGAAFSKILGSLLLNRNVRILMLGLDGAGKTTVLYRLKLNKYITTAPTIGFNAESLKYRNLEMTVWDVGGQE
Exanthemachrysis gayraliae Arf1TM    MGVVRRALCAALLALALVGTPVEGGSAFSKILGAILLNRNVRILMLGLDGAGKTTVLYRLKLNKYITTAPTIGFNAESLKYRNLEMTIWDVGGQE
Pavlova pinguis Arf1                 M-----------------------GLTFSKVFGRLFGKKEVRILMVGLDAAGKTTILYKLKLGEIVTTIPTIGFNVETVEYKNIQFTVWDVGGQD
Consensus/80%                        M.hsR.hbhhhLh.hAhhhs.hEGGtAFSKILGtLlLNRpVRILMLGLDGtGKTTVLYRLKLNKYITTAPTIGFNAESLKYRNLEMTVWDVGGQE

                                     <<<<<<<<<<<<<<<<<<<<<<<<<<<<<<<<<<<<<<  GTPase domain  >>>>>>>>>>>>>>>>>>>>>>>>>>>>>>>>>>>>>>>
Pavlova pinguis Arf1TM               KIRPLWRHYCRNTDALIFVVDSNDRDRLEEAGSELQQMLALPEMEAASVLVLANKQDMAGAADVYAICESLGLEEIQGHAWHIQACSAATGDGLF
Pavlova gyrans Arf1TM                KIRPLWRHYCRNTDALIFVVDSNDKDRLEEAGSELQQMLALPEMEAASVLVLANKQDMAGAADVYAICEALGLEEIQGHAWHIQACSAATGDGLF
Pavlovales sp. CCMP2436 Arf1TM       KIRPLWRHYCRNTDALIFVVDSNDRDRLEEAGSELQQMLALPEMEAASVLVLANKQDMAGAADVYAICEALGLEEIQGHAWHIQACSAATGDGLF
Diacronema lutheri Arf1TM            KIRPLWRHYCRNTDALIFVVDSNDRDRLDEAGSELQQMLALPEMETASVLVLANKQDMAGAADVYNICEALALEDIQGHAWHIQACSAATGDGLF
Exanthemachrysis gayraliae Arf1TM    KIRPLWRHYCRNTDALVFVVDSNDKDRLEEAGSELQQMLALPEMEAASVLVLANKQDMAGAADVYTICEALALDEIQGHAWHIQACSAATGDGLF
Pavlova pinguis Arf1                 KIRPLWRHYYQNTQGVIFVVDSNDRERVTEARDELHRMLNEDELRDAILLVFANKQDLPNAMNAAEITDKLGLHSLRQRQWYIQSTCATTGDGLY
Consensus/80%                        KIRPLWRHYCRNTDALIFVVDSND+DRL-EAGSELQQMLALPEMEsASVLVLANKQDMAGAADVYsICEtLtL--IQGHAWHIQACSAATGDGLF

                                     >>>>>>>>>
Pavlova pinguis Arf1TM               EGLDWLCKQL---KG
Pavlova gyrans Arf1TM                EGLDWMCKQL---KG
Pavlovales sp. CCMP2436 Arf1TM       EGLDWLCKQL---KG
Diacronema lutheri Arf1TM            EGLDWLCKQL---KG
Exanthemachrysis gayraliae Arf1TM    EGLDWLCKQL---KG
Pavlova pinguis Arf1                 EGLDWLSAQLAKSSK
Consensus/80%                        EGLDWLCKQL...KG
```

**Supplementary figure 18. Annotated alignment of Arl13 protein sequences from chlorarachniophytes.** Chlorarachniophytes exhibit an unusual domain architecture of the Arl13 protein, conserved across the group and comprised of an N-terminal CnB-related domain and a C-terminal EF-hand domain sandwiching the core GTPase domain. Arl13 from *Chlamydomonas reinhardtii* included in the alignment represents the standard domain architecture of Arl13 proteins. Accession numbers of the sequences coming from organisms that are not part of the core set of taxa analysed in this study are provided in Supplementary table 6.

```
                                                 <<<<<<<<<<<<<<<<<<<<<<<<<<<<<<<<<<<<<<<<<<<<<<   CnB-related domain   >>>>>>>>>>>>>>>>>>>>>>>
Bigelowiella natans Arl13           M---AAEPPGPKFDLKVLKDGAEQTSSDASKEEI-VSLLIKAGLSSEVASQHCRDFFDFFASGSSAKLVTAIEFYQEFVRMSCFRAVFQIRKMCRQMLDTSEVQKVLE
Lotharella amoebiformis Arl13       M--AALKPAGEKFKLPMLVKLAESGKNGSIDREMLVELTFEAGLSEEAAESHSSSFFSFFDHDNSGR-INLEDFVQEFIRMGCFRLVNAFKKRKEASYSTKDIQKELS
Lotharella globosa Arl13            MPVEAKHVPGPAFDISVLQKAAD-GDVAANEMEM-RLLCTGAGMSEDLASTHVKTFFRFFDVQKAGK-IKVAEFVKELIRMSCFKAVFALRKQGERKFPGERVRSELE
Partenskyella glossopodia Arl13     -----------------------------------------------------------------GS-VSLNDFVNEFVRMSCYRTVHALRQCGKMSLEPDTVQSELS
Chlamydomonas reinhardtii Arl13     M-----------FGLLV-------------------------------------NFYRF-----------------------C-------------------------
Consensus/80%                       M...........F.l.h.....................................sFapF......tp.lph.-FhpEblRMtCa+hV..b+p..c..b.spplpp.Lp

                                    >>>>>>>>>>>>>>>>>>>>>>>>>>>>>>>>>>>>>>>>>>>>>
Bigelowiella natans Arl13           NHLGARGGKDQYASMMKYAEKNMKAGGS-PQEKNKVDVKILEQWYWDKARSIIRSRKHFLEQKGYQV------------------GDLVGEGSSS----SNSDSKPED
Lotharella amoebiformis Arl13       LQLGAAGGREQFARLLAFAEEHXDPCSS----MGQVSIKTIERWYWTQARRLLLKRKDLLKKKGIDIG--KLLNVS-----------------------GSQIASKEY
Lotharella globosa Arl13            QHLGAIGGRTQYAKLLGFVEEKSEDGAS----AGSISVTMLERWYWSKAKALMEQRQALLNKKALTT-------------PATPSTDAPATTDEG----AAGRASPEE
Partenskyella glossopodia Arl13     IHLGSHGAKVYYKNLLEFLEKLGTPETGTALPHKKIPLESLMQWYLNMSNQLLXXXXELLTRKGFDVTCVTLVNIP(69)PSTPTTPTPTPTTPT(30)ASPSQNPAT
Chlamydomonas reinhardtii Arl13     ------------------------------------------------------------------------------------------------------------
Consensus/80%                       .pLGt.Gt+.ba.pbb.ahEc..ps.st.....plslp.lbpWYbsbtp.lb.....bLppKtbph.................................ts...p...

                                              <<<<<<<<<<<<<<<<<<<<<<<<<<<<<<<<<<<<<<<<<<<<<<<<<<<<<<<<<    Arl13 GTPase domain    <<<<<<<<<<<<<<
Bigelowiella natans Arl13           RIQKRTKREATICLLGIDNAGKSTFQCALLGETDPSVMPTFGFDMANFDAKFPSSNPFSCEVYDLPGGEDFGRSD-FWQNYYGDSHAFAFVVDSTD-RTRTSEALSEF
Lotharella amoebiformis Arl13       K---GERRKVSLLLVGLDNAGKTTLSKTFQGEEDPLIIPTCGFEFDSFYADYPKENPFECEIYDLAGDAGFGRAEGLWHNYFGDAHGIVFVVDSTD-TKRIKEVADEF
Lotharella globosa Arl13            R-----GRKATILLVGLDNAGKTTFSKAMIGEENPFIMTTFGFEIEEFESSFPEQNPFKCEFFDLAGGESFGRHD-FWSYYYGSAHAIAFIVDSTD-TTRMPEVIQQY
Partenskyella glossopodia Arl13     A-----TRRVVCVLTGLDNAGKTSVSKALIGEEDPFIMPTCGFDTEEFCAKYPASSPFRVTLYDLAGGDGFGRSEDLWSNYYEDAHFITFVIDSNDIESKQHEAIKEF
Chlamydomonas reinhardtii Arl13     --RKKTERKITIALLGLDNAGKTTLLNSIQGEVDRDTTPTFGFNSTTL-----NEGKYKIEVFDLGGGKNI---RGVWKKYLAEVHAIVYVVDAAD-PGRFEESKMTM
Consensus/80%                       ......pR+s*lhLlGLDNAGKTTbpptbbGEpDP.lbPThGF-hppF.tpaPppsPFcsElaDLtGGcsFGR.-.bWppYat-tHtIsFVVDSsD.psRbpEsbppa

                                    >>>>>>>>>>>>>>>>>>>>>>>>>>>>>>>>>>>>>>>>>>>>>>>>>>>>>>>>>>>>>>>>>>>>>>>>>>>>>>>>
Bigelowiella natans Arl13           SHLIGHSATRGKPIAIIANKQDIKEPQPYSGSDLKAMLGVPEKIPADRHCKLFECVATKGCKESKGVEGGVARALKWLLETLDPIYEKIDNEI--KEFKIKDRERADA
Lotharella amoebiformis Arl13       RRVSDHKGTAKKPLCILANKQDL--KGAMSGGDVAKALGS----SGDHPCKLFECVAREATG-----ERGVGRALKWMLTQLDPLFDDIDKRV--QKYKIADAKRAES
Lotharella globosa Arl13            QEVSQHPEAKGKPMVIIANKQDL--PDAKKVSEISKILNL-----KNPSQRIFECVSKPGS------IKGVSAALEWTLNTLHPMFATINKRV--EQYKIEERKKQAR
Partenskyella glossopodia Arl13     NHIIRHPKTNGKPILILANKQDL--PNSMSGPELCSLLNV-KDITGDRPCRIFECVAVEGGSKSK--PKGFDEAMEWVLGEVDSRFGDLDRMVTKEAEEVENREKAVK
Chlamydomonas reinhardtii Arl13     AEVLENQFMRDKPICIFANKQDL--PTAAPAAEVVKGLGL---ATCRNSHNVFPCTAKMPAGQDV--DHRLRDGLKWLVGTVDREFGRLDPRV--QTEAEEVRQEEAR
Consensus/80%                       pcl.pH..spsKPlsIlANKQDL..Pstbstt-lsphLsl....sscpsp+lFECVApbts.......+GlscALcWhLsplDsbFsclDpcV..ppbclcsRc+.sp

                                                                                     <<<<<<<<<<<<<<<<<<<<<<    EF-hand domain    >>>>>>>>>>>>>>>
Bigelowiella natans Arl13           EIKAQRKRTEE----WKRKEREENGEVENEGGD-----GKGKGDSALDSRGKVAAIFKMFDVDKDGYLNFKEFQSLL----SKT---------DGAEISEDMYRNEFC
Lotharella amoebiformis Arl13       ERKEKRERLE--------KQRKEVGTEEE----------KKXEEXPLDARGKVAKIFSEFDRDKDGFLNFDEFQALM----SAT-DP------DGEDLTDENFRDA-C
Lotharella globosa Arl13            EKEAARKRSEQ(25)AEKKKRVEKEEAQRRADEAA---MQAKQKSAMDAREKVAKIFERFDADKDGRLNFKEFKTLI----AAT-DP-----ENAAGITPSFFQFQIC
Partenskyella glossopodia Arl13     DQTAQREK----------NQKMEMATKQ-----------SAETHLDLDARGKTAIIFERFDDDKDGFLSLDEFQALM----DATTDP-----GDRQKISEEDFKDQIC
Chlamydomonas reinhardtii Arl13     KKKEREERLR--------KQREERLRQQKEEEERA(9)HDGKAPSLLAAGGGVVGAAA---AGVNGVM-VDEQQELR(58)SASGGP(110)GSGSGMTPD------A
Consensus/80%                       -pc.pRcR.c........KpRbE.hp.pp.........ptc.c.sLDARGKVA.IFpbFDsDKDGbLsbcEFQsLb....tAT.sP......st..l*.-.ap...C
```

```
                                   >>>>>>>>>>>
Bigelowiella natans Arl13          HELQ----VDPDKGVGVSALFVLLNQDEIALDKTYMELCC---G
Lotharella amoebiformis Arl13      DTLG----ADHKNGXFLSXLSGLCTQRTRARLTHTTRF------
Lotharella globosa Arl13           ERLR----CDAEKGVGLTDLYTVYERNLSDVDAHFELLGL---S
Partenskyella glossopodia Arl13    LALD----ADAEKGVHLAALYTLYSQDEEQLNAHFKMLDL----
Chlamydomonas reinhardtii Arl13    RELG(21)SDGGHGNSKGSFSLVHTSNKVVPVAPDLRAGI(6)A
Consensus/80%                      cpL.....sDsc+GshltsL.slbspsp.s.ssphbbbsh....
```
